# Biosynthesis of a major plant immunity hormone, salicylate, underwent multiple drastic changes during evolution of flowering plants

**DOI:** 10.1101/2025.08.19.670929

**Authors:** Fumiaki Katagiri, Evan A. Larson, Kunqi Hong, Kenichi Tsuda, Jerry D. Cohen, Adrian D. Hegeman

## Abstract

Salicylate (SA) is a major plant immunity hormone that activates immune responses against biotrophic pathogens. Between two known SA biosynthetic routes, the isochorismate synthase (ICS) and phenylalanine ammonia-lyase (PAL) pathways, which predominates in particular flowering plants has remained uncertain. We established a straightforward approach that directly quantifies the relative contributions of the ICS and PAL pathways and can also reveal an alternative, non-ICS/PAL route. In this approach, SA and shikimate (ShA), the precursor of SA in the ICS and PAL pathways, were metabolically labeled with [^13^C_6_]glucose in leaf tissue. We analyzed the LC-MS/MS data of the compounds using a metabolism-guided statistical model (the ICS-PAL model). This analysis allowed tracking of the origins of their ring and carboxyl carbons and, consequently, determination of the main SA biosynthetic routes. We surveyed SA biosynthetic routes in plants of the order Brassicales, other diverse eudicots, a monocot, magnoliids, and basal angiosperms. We found that the main SA biosynthetic routes changed at least twice during evolution of flowering plants: from a non-ICS/PAL pathway to the PAL pathway around the divergence time of monocots, and from the PAL pathway to the ICS pathway within Brassicales. Application of another metabolism-guided statistical model (the PKS-PAL model) to the SA data suggested that the non-ICS/PAL pathway involves a type III polyketide synthase (PKS). Typically, once a biosynthetic pathway is established for an important molecule, it is conserved in the subsequent lineage. SA biosynthesis in flowering plants took an unusual evolutionary path in which the pathways changed drastically multiple times. This unconventional evolutionary mechanism was employed for a major immunity hormone, likely due to rapid evolution of biotic environments, such as pathogens and insects that attack plants.

## INTRODUCTION

Generally, microbes with large population sizes, short generation times, and frequent horizontal gene transfers can evolve much faster than complex multi-cellular organisms, such as animals and plants [1, 2]. This principle of evolutionary rate differences leads to a fundamental biological question of how complex host organisms can keep up with the fast evolution of their microbial pathogens. The situation is even worse for hosts: a single host species or a single host individual could be attacked by multiple diverse microbial pathogens equipped with diverse molecular arms.

Vertebrates have evolved adaptive immunity, which applies single-cell selection on pre-generated enormously diverse immune-specialized cells to rapidly innovate specific solutions to particular pathogens in each individual [3]. However, plants and many other complex host organisms have only innate immunity: the pathogen recognition repertoire is genome-encoded and limited. Furthermore, pathogens have typically evolved to deliver effector molecules that target and compromise components of the immune machinery of the host organisms [4]. How could such host organisms with only innate immunity keep up with the evolution of their microbial pathogens?

Plants use special mechanisms to apparently buy time for slow evolution of specific solutions. First, plants have a highly resilient innate immune signaling network, in which subnetworks with different molecular compositions can back up the functions of one another if some subnetworks are compromised by pathogen effectors [5–8]. Such resilient networks effectively conceal the underlying molecular mechanisms from pathogens. Second, plants employ unconventional evolutionary mechanisms. For example, structurally homologous but not too similar positive and negative transcription regulator proteins of induced immunity could coevolve to protect the positive regulatory function from targeting effectors [9]. It is of interest to investigate what other unconventional evolutionary mechanisms host organisms with only innate immunity have employed.

Like other phytohormones, salicylate (SA) is involved in various physiological and developmental processes of plants. However, the most prominent role of SA is as an immunity hormone [10]. Roughly speaking, SA induces immune responses effective against biotrophic pathogens and suppresses immune responses induced by another immunity hormone, jasmonate (JA), which is effective against necrotrophic pathogens and insects [11]. Many pathogens are biotrophic or hemibiotrophic (i.e., biotrophic in early infection stages and necrotrophic in late stages), and immune responses induced by SA are crucial for defense against these pathogens [12].

Despite its central role in immunity, biosynthesis of SA in plants has been well studied in only a limited number of species. In Arabidopsis, SA is primarily synthesized via the isochorismate synthase (ICS) pathway [13]. We recently demonstrated that the Arabidopsis-type ICS pathway, which requires the likely isochorismate (IC) transporter EDS5 and the IC-glutamate conjugating enzyme PBS3 in addition to ICS [14, 15], evolved after diversification of the order Brassicales [16]. The order Brassicales is just one of 44 extant eudicot orders [17]. In the monocot, rice, the phenylalanine ammonia-lyase (PAL) pathway is a major SA biosynthetic pathway [18], and the contribution of the ICS pathway to basal SA accumulation is negligible [19]. The ICS and PAL pathways bifurcate at chorismate, which is the product of the shikimate (ShA) pathway, i.e., both the ICS and PAL pathways act downstream of the ShA pathway. The first committed steps after the bifurcation point are mediated by ICS and chorismate mutase (CM) for the ICS and PAL pathways, respectively. We collectively define the PAL pathway as any SA biosynthetic pathway using CM at the bifurcation point downstream of the ShA pathway (Fig 1A).

**Fig 1.**
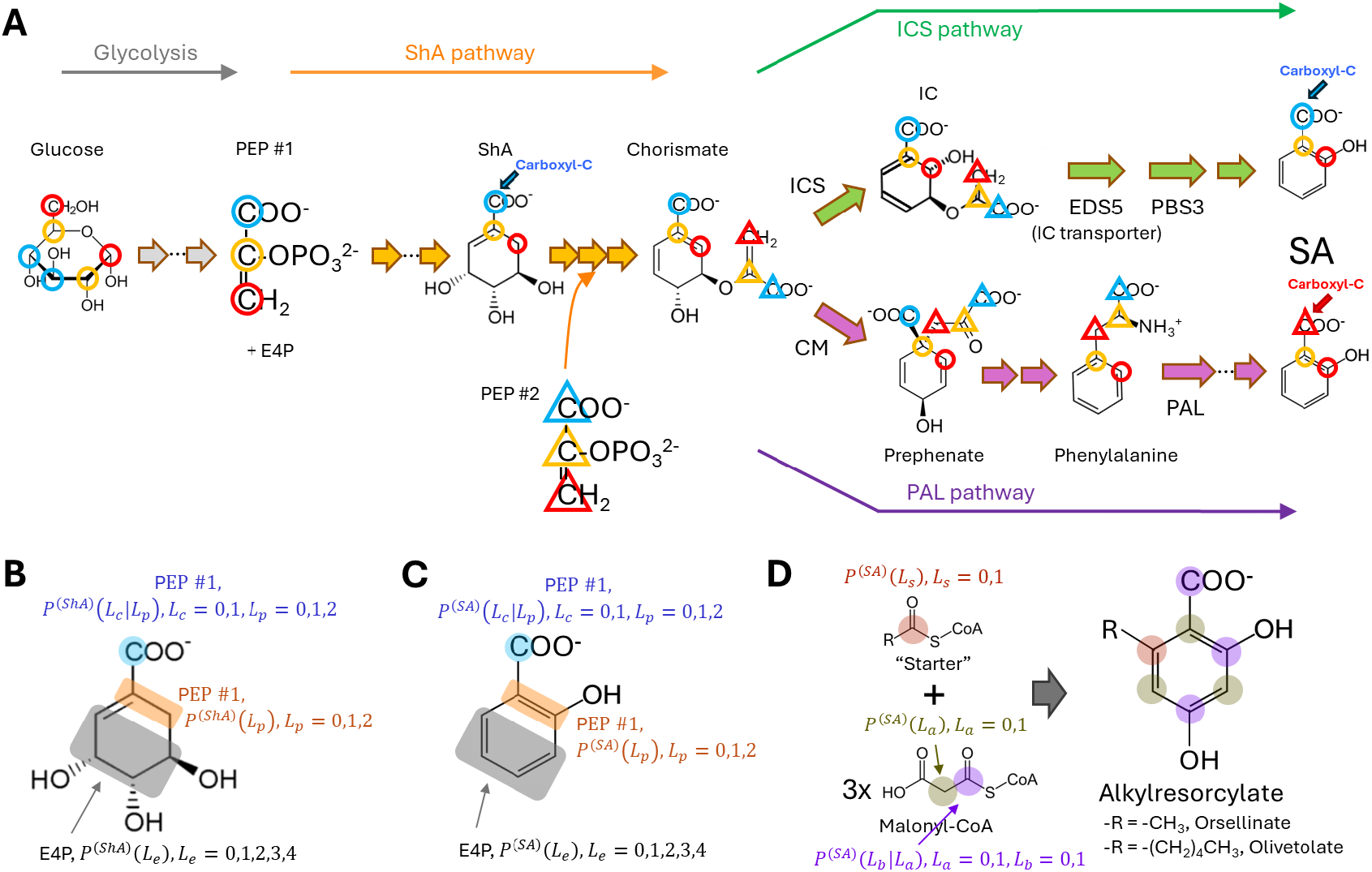
The source of the SA carboxyl carbon is distinct in different SA biosynthetic routes. (A) Flow of carbons from glucose to phosphoenolpyruvate (PEP) to SA through glycolysis, the shikimate (ShA) pathway, the isochorismate synthase (ICS) pathway, and the phenylalanine ammonia-lyase (PAL) pathway. The carbons of interest are 1-C (blue), 2-C (yellow), and 3-C (red) of PEP #1 (circle) and PEP #2 (triangle). They are tracked from glucose to SA. The other four carbons in ShA and SA (no circles or triangles) originate from the four carbons of erythrose-4-phosphate (E4P). The carboxyl carbons of ShA and SA, synthesized through the ICS and PAL pathways, are indicated by “Carboxyl-C”. Note that the carboxyl carbons of ShA and the ICS-pathway-synthesized SA originate from the 1-C of PEP #1 (blue circle) and that none of the bonds originated from PEP #1 are broken in them. In contrast, the carboxyl carbon of the PAL-pathway-synthesized SA originates from the 3-C of PEP #2 (red triangle), and the carboxyl carbon of the PAL-pathway-synthesized SA is randomized relative to its ring. (B, C) The probability variables of [^13^C]-labeled carbon states for ShA (B) and SA (C) used in the ICS-PAL model. The 3-, 4-, 5-, and 6-C (gray-rectangular-shaded) of the rings in ShA and SA originate from E4P, and the probability of having the number *L*_*e*_ of [^13^C]-labeled carbons among the four carbons in a single molecule is denoted as *P*^(*ShA*)^(*L*_*e*_) and *P*^(*SA*)^(*L*_*e*_), respectively. Similarly, the 1- and 2-C (orange-rectangular-shaded) of the rings in ShA and SA originate from PEP #1, and the probability to have the number *L*_*p*_ of [^13^C]-labeled carbons among the two carbons in a single molecule is denoted as *P*^(*ShA*)^(*L*_*p*_) and *P*^(*SA*)^(*L*_*p*_), respectively. *P*^(*ShA*)^(*L*_*p*_) = *P*^(*SA*)^(*L*_*p*_) was assumed in the model (the labeling kinetics assumption). The conditional probability of having the carboxyl carbon (blue-circle-shaded) of ShA and SA [^13^C]-labeled (*L*_*c*_ = 1) given the number *L*_*p*_ is denoted as *P*^(*ShA*)^(*L*_*c*_|*L*_*p*_) and *P*^(*SA*)^(*L*_*c*_|*L*_*p*_), respectively. By comparing *P*^(*ShA*)^(*L*_*c*_|*L*_*p*_) and *P*^(*SA*)^(*L*_*c*_|*L*_*p*_), the fraction of the SA carboxyl carbon that is conserved from the ShA carbon (the bond between the carboxyl carbon and the ring is never broken) can be obtained, and thus the proportion of the SA molecules in a sample that are synthesized through the ICS pathway can be estimated when all SA molecules are synthesized through the ICS or PAL pathways. (D) The probability variables of [^13^C]-labeled carbon states used in the PKS model for a hypothetical SA biosynthetic pathway involving a type III polyketide synthase (PKS). Via this pathway, the 6-C (salmon-shaded; the six ring carbons in alkylresorcylase are numbered from the one with the carboxyl carbon attached) of the alkylresorcylase ring originates from the carbonyl carbon of the “starter” acyl-CoA and the probability of having it [^13^C]-labeled is denoted as *P*^(*SA*)^(*L*_*s*_). The 1-, 3-, and 5-C (olive-shaded) of the alkylresorcylase ring originate from the methylene carbons of three malonyl-CoA molecules and the probability of having each of the three carbon atoms [^13^C]-labeled is denoted as *P*^(*SA*)^(*L*_*a*_). The carboxyl carbon and the 2- and 4-C (violet-shaded) of the ring originate from the carbonyl carbons in the thiolester of three malonyl-CoA molecules and the probability of having each of the three carbon atoms [^13^C]-labeled is denoted as the conditional probability given *L*_*a*_, *P*^(*SA*)^(*L*_*b*_|*L*_*a*_). As the LC-MS/MS data provide information on the bond between the carboxyl carbon and the ring, defining the labeling probability of the carboxyl carbon conditional to the 1-C of the ring gives more metabolism-guided constraints in the model.

Determining the relative contributions of the two SA biosynthetic pathways using loss-of-function genetic approaches is challenging. This is because the SA biosynthetic system typically involves feedback regulation from SA. For example, in Arabidopsis, *PAL4* in addition to *ICS1, EDS5*, and *PBS3* are transcriptionally induced by SA (e.g., Arabidopsis eFP browser [20] and Table S3 in [21]). In this case, the outcome of an *ics1* knockout may include reduction of SA synthesis via the PAL pathway due to reduced *PAL* gene expression, in addition to the reduction of SA produced by the ICS pathway. *Vice versa*, the outcome of a *pal* knockout may include reduction of SA synthesis via the ICS pathway. This positive feedback interpretation could explain two apparently contradictory observations in Arabidopsis: knocking out both of the *ICS* genes (*ics1 ics2*) resulted in a 96% reduction of SA accumulation induced by UV-C treatment [22] and knocking out all four *pal* genes resulted in a 50% reduction of SA accumulation induced by a pathogen [23] (Fig S1).

We developed a straightforward approach to directly quantify the relative contributions of the ICS and PAL pathways to SA biosynthesis, in which each assay requires only a small leaf. In this approach, ShA and SA were metabolically labeled by feeding [^13^C_6_]glucose to leaves. The [^13^C]-labeling patterns of the ring and carboxyl carbons in ShA and SA were measured by LC-MS/MS. By fitting a metabolism-guided statistical model to the LC-MS/MS data, the proportions of the conservation and randomization of the [^13^C]-labeling state of the carboxyl carbon in SA compared to that of ShA were determined. The carboxyl carbon labeling state is conserved in SA synthesized via the ICS pathway, while it is randomized in SA synthesized via the PAL pathway. Thus, the proportions of the conservation and randomization correspond to the proportions between the ICS and PAL pathways. Furthermore, this approach can detect non-ICS/PAL pathway(s) that generate a carbon labeling state different from possible combinations of the states expected for the ICS and PAL pathways. We initially surveyed 6 Brassicales species, 9 diverse non-Brassicales eudicot species, and a monocot species, rice. The core Brassicales clade, which includes the families Resedaceae, Cleomaceae, and Brassicaceae, mainly used the ICS pathway (ICS-main). Basal Brassicales plants, except moringa, had very low SA synthesis rates, suggesting that strong reduction of the SA synthesis rate preceded the innovation of the new ICS pathway. All the other eudicot and monocot plants mainly used the PAL pathway (PAL-main), suggesting that the most recent common ancestor (MRCA) of eudicots and monocots mainly used the PAL pathway. We expanded our survey to include two basal angiosperm and two magnoliid species and found that they primarily use a non-ICS/PAL pathway(s) for SA synthesis. Modeling of the carbon labeling state of SA suggests involvement of a type III polyketide synthase (PKS) in the non-ICS/PAL pathway. Thus, at least three very different pathways are used as the main SA biosynthetic pathways in flowering plants. Well-established biosynthetic pathways are usually conserved through evolution. With rapid changes in their biotic environments, flowering plants likely had to use an unconventional evolutionary mechanism, which is to repeatedly drastically change major biosynthetic pathways for the important immunity hormone SA.

## RESULTS

### Principles of our approach for determining the ICS and PAL pathway contribution proportions in SA biosynthesis

We initially assumed that the ICS and PAL pathways were the only possible SA biosynthetic pathways in plants based on the knowledge of the time [24]. Both the ICS and PAL pathways act downstream of the ShA pathway. The ten carbon atoms in chorismate, the product of the ShA pathway, originate from two molecules of phosphoenolpyruvate (PEP #1 and #2) and one molecule of erythrose-4-phosphate (E4P) (Fig 1A). When all the carbon atoms in SA made through the ICS and PAL pathways are tracked, the carboxyl carbon of SA originates from 1-C of PEP #1 (blue circled “C”) via the ICS pathway or from 3-C of PEP #2 (red triangled “C”) via the PAL pathway. Therefore, if the origin of the carboxyl carbon can be quantitatively tracked, the proportion of the ICS and PAL pathway contributions can be determined. The major fragmentation pattern of SA by MS/MS, which results in loss of the carboxyl group (https://mona.fiehnlab.ucdavis.edu/spectra/display/WA000474), would allow determination of the proportion of [^13^C]-labeled carbon at the carboxyl carbon of SA. We used glycolysis to [^13^C]-label PEP from [^13^C]-labeled glucose and then the ShA pathway to [^13^C]-label the carboxyl carbons of SA from [^13^C]-labeled PEP (Fig 1A).

To determine the origin of the carboxyl carbon, we investigated the conservation and randomization of the carboxyl carbon of SA from the carboxyl carbon of ShA via the ICS and PAL pathways, respectively. We statistically modeled the labeling states of the carboxyl carbon and the ring carbons to estimate the probabilities of the labeling state of the carboxyl carbon conditional to the PEP #1-originated ring carbon labeling state in each of ShA and SA (*P*^(*ShA*)^(*L*_*c*_|*L*_*p*_) and *P*^(*SA*)^(*L*_*c*_|*L*_*p*_) in Figs 1B and 1C, respectively). We assumed that the labeling kinetics from ShA to SA is much faster than that from glucose to ShA or that most newly synthesized SA molecules were synthesized after reaching a steady labeling condition (this assumption is hereafter referred to as the labeling kinetics assumption). Then the proportion of the ICS pathway contribution in the total synthesized SA, *μ*, can be calculated as:

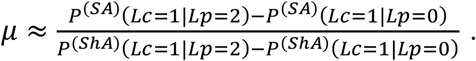

Since we only considered the ICS and PAL pathways, the proportion of the PAL pathway contribution is 1 − *μ*. Details of this metabolism-guided statistical model, the ICS-PAL model, and the R script used are provided in Text S1 and Data S1.

The advantages of this approach are four-fold. First, the contribution proportions of the ICS and PAL pathways are measured directly, so the approach is not affected by a potential complication due to the positive feedback from SA. Second, each assay only requires a small leaf (the size of an Arabidopsis adult leaf) for the plant material. Third, the *μ* value can be estimated for each leaf sample, so that large sample-to-sample labeling efficiency variation, which we observed in our preliminary study, is not an issue. This vastly reduced the required number of replicates for each plant species. Fourth, [^13^C_6_]glucose is much less expensive than position-specifically labeled glucose. All these practical advantages allowed us to survey many plant species, including exotic plant species from a conservatory.

### [^13^C]-labeling procedure

We designed the experimental procedure so that [^13^C_6_]glucose fed to the leaf samples was the major carbon source for the leaf tissue. First, excised leaves with their petioles in water were placed in the dark for ~20 hours to deplete the leaf starch storage. Second, the leaves were treated with UV-C (254 nm), which is known to induce SA synthesis in diverse plant species [25, 26]. Third, the leaves were moved to 50 mM [^13^C_6_]glucose solution and kept under a low light condition (PAR = 3.5 μmol/m^2^/s). The low light condition was used to keep stomata reasonably open for two purposes: transpiration for efficient uptake of the labeled glucose solution by the leaf sample; good respiration conditions to facilitate flux through glycolysis and efficient PEP labeling. In addition, the low light condition reduced the flux of unlabeled carbons from photosynthesis to the labeling metabolic pathways. After 6 to 30 hours of labeling, the leaves were collected for LC-MS/MS analysis. The LC-MS/MS data for all samples that resulted in reliable *μ* estimates are provided in Table S1. The precise UV-C treatment and labeling times and the fitted model parameter values for each sample are provided in Table S2.

### Arabidopsis mainly uses the ICS pathway, and tomato mostly uses the PAL pathway

We generated three types of plots that allow visual interpretations of whether the SA biosynthetic pathway used in a leaf sample is likely a combination of the ICS and PAL pathways and, if so, whether it is mainly the ICS or PAL pathway (“ICS-main” or “PAL-main”) and whether the data from the leaf sample satisfy the labeling kinetics assumption. The three plots are called the carboxyl carbon labeling profile, the ring carbon labeling profile, and the ring carbon labeling parameter plot. A carboxyl carbon labeling profile, such as Fig 2A, shows the proportion of [^13^C]-labeled carbon at the carboxyl carbon (*y*-axis) conditional to each number of the labeled ring carbons (0 to 6 in the *x*-axis) for ShA (black) and SA (pink). A ring carbon labeling profile, such as Fig 2B, shows the abundances relative to the abundance at *x* = 6 of ShA (black) or SA (pink) molecules with rings containing different numbers of labeled carbons. In the carboxyl carbon and ring carbon labeling profiles, the data are shown by lines so that the overall profile shapes across the number of the labeled ring carbons are easy to grasp, and the modeled values are shown by point characters (ShA, black square; SA, pink circle) so that the model fit can be easily visualized. In a ring carbon labeling parameter plot, such as Fig 2C, the model parameters for the proportions of different numbers of labeled carbons among the E4P-originated (*P*(*L*_*e*_ = 0,1,2,3,4) in Figs 1B and 1C; represented by “e0” to “e4” in gray in the plot) and the PEP #1-originated (*P*(*L*_*p*_ = 0,1,2) in Figs 1B and 1C; represented by “p0” to “p2” in red in the plot) ring carbons are compared between the models fit separately for ShA (*x*-axis) and SA (*y*-axis). The three plots for a single leaf sample in this order are horizontally aligned (e.g., Figs 2A to 2C), and the sample label (e.g., “#E2”) and the sample plant species/variety (e.g., Arabidopsis Col-0) are indicated in the left plot, such as those in Fig 2A. The color of the sample label shows which of the two versions of the ICS-PAL model was selected according to their AIC values: the common-ring model version (red; this version has 12 independent variables and assumes the same parameter value for each of e0 to e4 and p0 to p2 between ShA and SA) or the different-E4P-ring model version (blue; this version has 16 independent variables and assumes the same parameter value for each of p0 to p2 but different values for each of e0 to e4 between ShA and SA).

**Fig 2.**
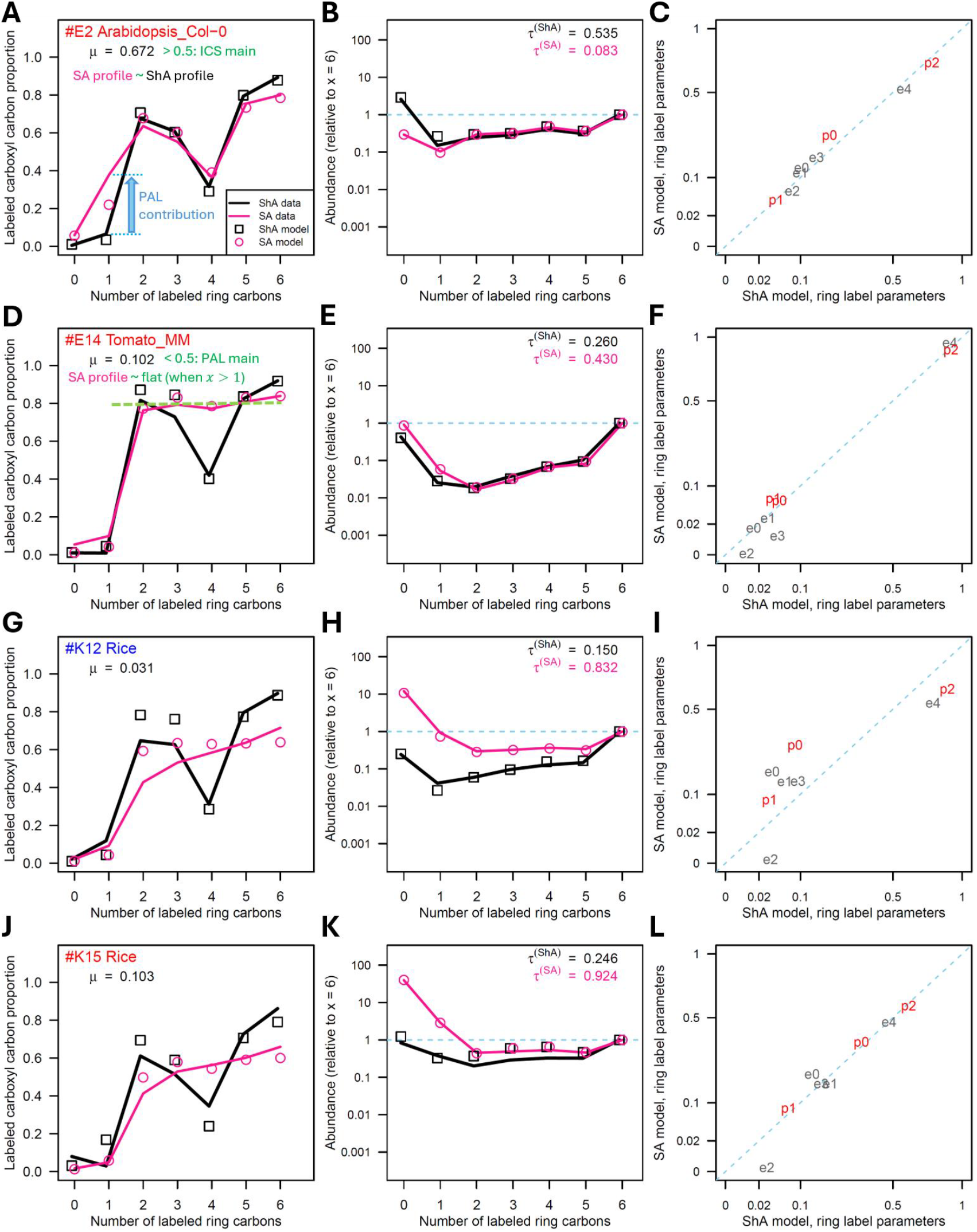
While Arabidopsis Col-0 is ICS-main, tomato and rice are PAL-main. Three types of plots, the carboxyl carbon labeling profile (A, D, G, J), the ring carbon labeling profile (B, E, H, K), and the ring carbon labeling parameter plot (C, F, I, L), for each of four samples from Arabidopsis Col-0 (A-C), tomato MoneyMaker (D-F), and rice (two samples, G-I and J-L) are shown. The sample labels and the plant species names are indicated on the top left of the carboxyl carbon labeling profiles. The color of the sample label letters indicates that either the common-ring model version (red; assumed *P*^(*ShA*)^(*L*_*p*_) = *P*^(*SA*)^(*L*_*p*_) and *P*^(*ShA*)^(*L*_*e*_) = *P*^(*SA*)^(*L*_*e*_), see Figs 1B and 1C) or the different-E4P-ring model version (blue; assumed *P*^(*ShA*)^(*L*_*p*_) = *P*^(*SA*)^(*L*_*p*_) only) of the ICS-PAL model was used. In the carboxyl carbon and ring carbon labeling profiles, the data are shown by lines, and the model estimates are shown by point characters for ShA (black) and SA (pink) as in the legend in A (*x*-axis integer values are slightly offset for better visualization). Visual features in the carboxyl carbon labeling profiles for the ICS-main (the proportion of the ICS pathway contribution, *μ* > 0.5) and PAL-main (*μ* < 0.5) cases are described in A and D. The blue arrow “PAL contribution” in A indicates that this large discrepancy between the ShA and SA data values at *x* = 1 demonstrates that there is a non-negligible PAL pathway contribution in this ICS-main sample. In the ring carbon labeling profiles, the *y*-axis is on a log scale, and *τ*^(*ShA*)^ (black) and *τ*^(*SA*)^ (pink) show the estimated proportion of preexisting and remaining ShA and SA, respectively. In the ring carbon labeling parameter plots, p0 to p2 (red) represent *P*^(*u*)^(L_p_), L_*p*_ = 0,1,2, and e0 to e4 (gray) represent *P*^(*u*)^(L_e_), L_*e*_ = 0,1,2,3,4, respectively, where *u* is “ShA” or “SA” (see Figs 1B and 1C). The parameter values shown on a square-root scale were estimated by fitting the ShA and SA parts of the ICS-PAL model separately (*x*- and *y*-axis, respectively). The corresponding three-type plots for 76 samples with reasonable model fits are given in Fig S2.

The carboxyl carbon labeling profiles of one Arabidopsis Col-0 (#E2; Fig 2A) leaf sample and one tomato cv. Money Maker (#E14; Fig 2D) leaf sample are shown to compare the ICS-main and PAL-main patterns. The profiles of the ShA data (black lines) were similar between Arabidopsis Col-0 and tomato: low at *x* = 0,1,4 and high at the other *x* values. It was low at *x* = 0,1 because these were mostly represented by the remaining ShA molecules that already existed before labeling (*τ*^(*ShA*)^ in Figs 2B and 2E is the model estimate of the proportion of the preexisting and remaining ShA in the sample). The reason it was low at *x* = 4 is due to the specific metabolic process. The ShA ring carbons originate from PEP #1 and E4P (Fig 1A). With [^13^C_6_]glucose, most labeled PEP and E4P molecules have all their carbons labeled (e4 ≫ e1, e2, e3 and p2 ≫ p1 in Figs 2C and 2F; Text S1, sections I.1 and II.3). Under this condition, the majority of the ShA molecules with *x* = 4 were made from no-carbon-labeled PEP #1 and all-carbon-labeled E4P, which had the no-labeled carboxyl carbon, and thus the labeled carboxyl carbon proportion was low at *x* = 4. The characteristics of the ShA carboxyl carbon labeling profile, low at *x* = 0,1,4, were conserved in all plant species we investigated when the ShA signals were sufficiently strong for model fitting.

The profiles of the SA data (pink lines) were very different between Arabidopsis Col-0 and tomato. The SA data profile was similar to the ShA data profile in Arabidopsis Col-0, indicating that the majority of the SA molecules had the carboxyl carbon conserved from the carboxyl carbon of ShA, which is the characteristic of the ICS pathway. Accordingly, the *μ* estimate for this Arabidopsis sample was 0.672 (> 0.5; ICS-main). With tomato, the SA data profile plateaued around *y* = 0.8 when *x* > 1 (yellow green horizontal dashed line in Fig 2D). This is the characteristic of the PAL pathway, indicating that the carboxyl carbon of ShA was replaced with a carbon from another source, i.e., the carboxyl carbons of most SA molecules originated from C-3 of PEP #2. Accordingly, the *μ* estimate for this tomato sample was 0.102 (< 0.5; PAL-main). The reason that the SA profiles were low at *x* = 0,1 is the same as that for the ShA profiles: these are mostly represented by the preexisting and remaining SA molecules (*τ*^(*SA*)^ in Figs 2B and 2E are the model estimates of the proportion of the preexisting and remaining SA in the samples).

Although the ShA and SA carboxyl carbon labeling profiles were similar in Arabidopsis Col-0, there was a distinct difference: the labeled carboxyl carbon proportion of SA was much higher than that of ShA at *x* = 1 (indicated by a blue “PAL contribution” arrow in Fig 2A). If all SA molecules were made through the ICS pathway, the SA proportion value should be very similar to the ShA proportion value. The theoretical maximum of the SA proportion value, which is calculated under the assumption that all preexisting ShA was metabolically inactive (see below for the metabolically inactive ShA pool), is (ShA proportion value) * (1 − *τ*^(*SA*)^)/(1 − *τ*^(*ShA*)^). This theoretical maximum is 0.13 for the #E2 sample while the observed SA proportion value was 0.38 (Figs 2A and 2B). This high SA proportion value at *x* = 1 can be explained if the PAL pathway is used partially. If the labeling kinetics of the part of the pathway from ShA to PEP #2 incorporation (the ShA kinase and the 5-enolpyruvylshikimate-3-phosphate synthase reaction steps) is faster than that of the part of the pathway from PEP #1 to ShA, the [^13^C_6_]glucose-labeled PEP #2 can be incorporated into chorismate before the chorismate ring carbons get labeled from [^13^C_6_]glucose. Naturally ~6% of the ring has a single ^13^C (since the natural abundance of ^13^C is 0.0107 [27]), and these rings with natural single ^13^C (*x* = 1) could have labeled carboxyl carbons from the labeled PEP #2-originated carbons via the PAL pathway. Thus, this high SA proportion value at *x* = 1 compared to the ShA proportion value indicates that a non-negligible proportion of the SA molecules was synthesized through the PAL pathway. Therefore, although Arabidopsis Col-0 mainly uses the ICS pathway, it uses the PAL pathway as well, although to a lesser extent. This fact was reflected in its *μ* estimate of 0.672 (Fig 2A), which is significantly lower than 1 (*p* = 0.0021, post hoc, two-sided).

If the labeling kinetics assumption is satisfied for both the PEP #1- and E4P-originated carbons, the labeling states of the newly labeled ring carbons should be very similar between ShA and SA via either the ICS or PAL pathways. In the ring carbon labeling profiles of ShA and SA for both Arabidopsis Col-0 and tomato, the profiles are very similar where *x* > 1 (Figs 2B and 2E), indicating that the labeling states of the newly labeled ring carbons were very similar, i.e., the labeling kinetics assumption was satisfied. Since the assumption was satisfied for both the PEP #1- and E4P-originated carbons, the common-ring model version was selected for these samples. The discrepancy between the ShA and SA ring carbon labeling profiles where *x* ≤ 1 is highly affected by the proportions of the preexisting and remaining ShA and SA, *τ*^(*ShA*)^ and *τ*^(*SA*)^. If ShA forms a single well-mixed pool, *τ*^(*ShA*)^ ≤ *τ*^(*SA*)^ (the ShA profile is lower than the SA profile where *x* ≤ 1): e.g., #E14 tomato may have such one well-mixed ShA pool (Fig 2E). If *τ*^(*ShA*)^ > *τ*^(*SA*)^, ShA has a metabolically inactive pool (i.e., very little labeling and very little conversion to SA of the pool of ShA during the labeling time) as well as a metabolically active pool (so that some SA molecules get labeled). Under this condition, the SA profile could be lower than ShA profile where *x* ≤ 1 (e.g., #E2 Arabidopsis Col-0, Fig 2B). The proportion of the inactive pool relative to total ShA amount is at minimum *τ*^(*ShA*)^ − *τ*^(*SA*)^.

The model-fitted values corresponding to the carboxyl carbon labeling profiles and the ring carbon labeling profiles by the ICS-PAL common-ring model version are included in Figs 2A, 2B, 2D, and 2E (black squares for ShA and pink circles for SA), showing that the model fits very well in these samples. The ICS-PAL model is a statistical model constrained by the metabolic origins of the carbons in ShA and SA, under the assumption of the linear combination of the ICS and PAL pathways (Figs 1A to 1C and Text S1). Note that these two types of profiles for the ShA and SA data represent 26 independent observations (i.e., 7 and 6 for the carboxyl carbon and ring carbon labeling profiles, respectively, for each of ShA and SA), and that the common-ring model version has 12 independent variables: i.e., the model has 14 degrees of freedom. Thus, good model fits in the profiles of these samples strongly suggest that the underlying metabolic processes are indeed the ICS and/or PAL pathways and that their *μ* estimates are accurate.

The ring carbon labeling parameter plot visually examines the labeling kinetics assumption with a higher resolution than the ring carbon labeling profile. The models for ShA and SA were separately fit to the ShA and SA data. Then the ring carbon labeling parameter values were compared between the ShA and SA models. If the data support the assumption for both the PEP #1- and E4P-originated carbons, the parameter values are very similar and the points, p0 to p2 in red and e0 to e4 in gray, are generally on the diagonal line of *y* = *x* (blue dashed line). Both the Arabidopsis Col-0 and tomato sample data well support the assumption (Figs 2C and 2F), and these plots justify use of the ICS-PAL common-ring model version.

The summary of the *μ* estimates across multiple biological replicates for Arabidopsis Col-0 and tomato is included in Fig 3B (the numbers of biological replicates, 12 and 11, respectively, are shown under “n” on the right flank of the plot). The means (black circle) and their 95% confidence intervals (error bars) are *μ* = 0.712±0.030 for Arabidopsis Col-0 and *μ* = 0.045±0.038 for tomato, demonstrating that the trends observed in the examples in Figs 2A to 2F are typical for these plants: i.e., Arabidopsis Col-0 and tomato are ICS-main and PAL-main, respectively.

**Fig 3.**
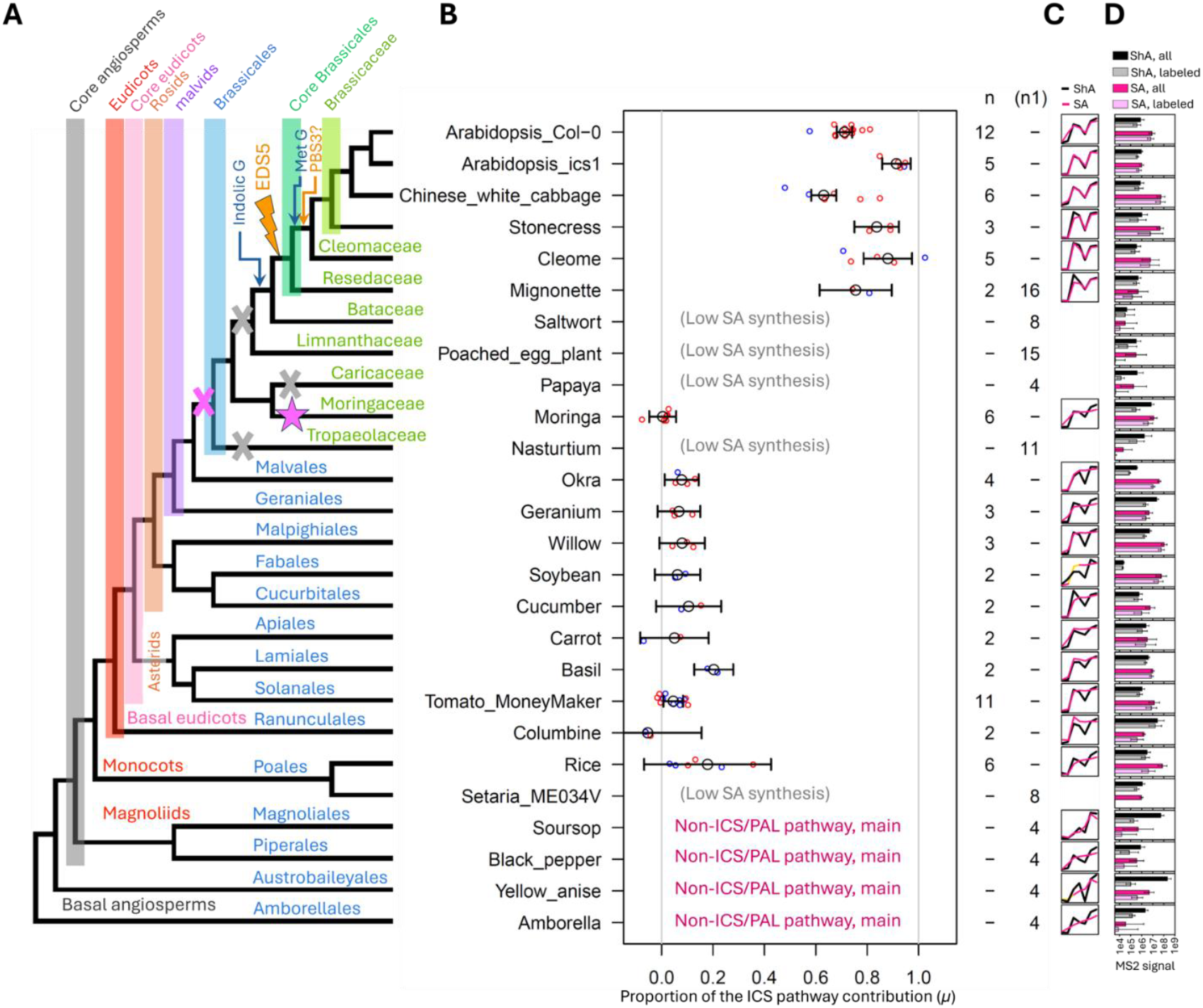
Non-Brassicales Eudicots are PAL-main. (A) The phylogenetic relationships among the plants analyzed are shown relative to the Arabidopsis lineage (top lines of the tree and vertical color bars for the clades). In addition to the taxonomic orders (blue letters), the taxonomic families within the order Brassicales (green letters) and higher-level clade names, basal angiosperms, magnoliids, monocots, basal eudicots, and asterids, are indicated. In the order Brassicales, we do not know whether the SA biosynthesis rate became very low in three independent events (three gray “X”s) or whether the MRCA of the order acquired a very low SA biosynthesis rate (pink “X”) and the Moringaceae linage regained the PAL pathway (pink star). The points on the phylogeny when indolic glucosinolates (“Indolic G” in blue), *EDS5* (orange lightning with “EDS5”), methionine-derived glucosinolates (“Met G” in blue), and *PBS3* (“PBS3?” in orange, showing the likely timing) were innovated [16, 41] are also indicated. (B) The proportion of the ICS pathway contribution to SA synthesis (*μ*) is shown for the indicated plants. *μ* = 1 and 0 represent 100% ICS and 100% PAL pathway contributions, respectively. Since the *μ* value is calculated as the coefficient of the linear combination between the 100% ICS and 100% PAL pathway contribution cases, with some measurement errors the *μ* estimate could be slightly outside the range of [0,1] when the actual *μ* value is close to the boundary values. For each plant, the mean (black circle) and its 95% confidence interval (black error bar) across biological replicates are shown. The *μ* estimate of each sample is shown by a small circle for the common-ring model version (red) or the different-E4P-ring model version (blue) selected for the sample. “(Low SA synthesis)” for four basal Brassicales plants and one monocot plant indicates that low SA labeling and consequential missing measurements in the LC-MS/MS analysis prevented us from estimating the *μ* values. “Non-ICS/PAL pathway, main” indicates that the basal angiosperm and magnoliids plants mainly use a non-ICS/PAL pathway and cannot be modeled by the ICS-PAL model. The number of well-modeled samples for each plant are shown on the right of the plot under “n”: the *μ* estimates of these samples are used in 1-way ANOVA in the plot. (C) The carboxyl carbon labeling profiles of the ShA (black) and SA (pink) data of a representative sample of each of the plants with the *μ* estimates or the non-ICS/PAL main plants are shown (the representative samples used are from the top: #E2, E3, F4, L9, L18, H17, H21, G9, J4, J5, F21, F24, G5, G13, E14, I8, K15, N21, N5, N8, and N1; Fig S2 for larger profiles with model estimates). The SA data profile compared to the ShA data profile tells which of the three pathways, the ICS, PAL or non-ICS/PAL pathways, is mainly used (See text). (D) The amounts of total and labeled ShA and SA in the samples for each plant (mean and SD (error bar); color legends are on the top). The signal values from MS/MS were used as approximate amounts. The ShA or SA signals for two or more labeled ring carbons were used as the labeled amounts. The *x*-axis is log scaled, and each tick shows one 10-fold (one log_10_) difference (see the tick labels at the bottom). For some plants, the samples that were not used for the ICS-PAL modeling were included (their numbers are under “(n1)” on the right end of B) in addition to the samples well modeled (the numbers under “n”). The *μ* estimates of the individual samples are given in Table S2.

### Rice mainly uses the PAL pathway

Fitting the ICS-PAL model to rice data was often challenging, probably because of a high basal SA level [28]. A high preexisting and remaining SA, which corresponds to a very high *τ*^(*SA*)^, slows the labeling kinetics from ShA to SA, i.e., the data violates the labeling kinetics assumption. For example, the ring carbon labeling profile of the #K12 sample showed higher SA values than ShA values where 1 < *x* < 6 (Fig 2H). Accordingly, the ICS-PAL different-E4P-ring model version was selected as shown by the blue color of the sample label in Fig 2G. This model allows the ring labeling parameter values for the E4P-originated ring carbons to be different between ShA and SA: the model assumes a situation where the E4P-originated ring carbon labeling kinetics may be fast, but the PEP #1-originated ring carbon labeling kinetics are slow compared to the labeling kinetics from ShA to SA, which still satisfies the labeling kinetics assumption. However, the ring carbon labeling parameter plot of #K12 shows that the PEP #1-originated ring carbon labeling parameter estimates (p0, p1, and p2 in red) are not very similar between ShA and SA (Fig 2I). Thus, the data violates the labeling kinetics assumption. Understandably, the model fit in the carboxyl carbon labeling profile is not great (Fig 2G) although the fit is good in the ring carbon labeling profile (Fig 2H). In contrast, with the #K15 sample, the labeling kinetics assumption is well-supported (Figs 2K and 2L), and the model fit is decent (Figs 2J and 2K). Thus, we could have a higher confidence in the *μ* estimate for #K15 than that for #K12. In any case, in rice, the SA carboxyl carbon labeling profiles more closely resemble the PAL-main type, and the *μ* estimates for both samples are close to 0. Thus, we concluded that the monocot rice mainly uses the PAL pathway for SA synthesis.

The mean and its 95% confidence interval of the *μ* estimates across six biological replicates for rice were *μ* = 0.179±0.247 (Fig 3B). The main reason the 95% confidence interval for rice is much larger than those for Arabidopsis Col-0 and tomato is that the *μ* value of each leaf sample used in 1-way ANOVA in Fig 3B was weighted according to its standard error estimated by its ICS-PAL model (Table S2). Since the model fits for rice samples were not great in general as seen in the above examples, the large standard errors of *μ* with the rice samples were reflected in the large confidence interval.

### Surveys of many flowering plants for their main SA biosynthetic pathways were conducted

Taking advantage of our approach, we conducted focused and broad surveys of flowering plants for their main SA biosynthetic pathways (Fig 3). Fig 3A shows a phylogenetic tree of the plant species studied, in respect to the Arabidopsis Col-0 lineage, which is the uppermost lineage in the figure. Hereafter, we use the terms of the divergence and an ancestor of a lineage specifically for the divergence from and an ancestor on the Arabidopsis lineage, unless otherwise specified. One survey was focused on the order Brassicales (blue vertical bar in Fig 3A), and eight families in the order were represented by at least one species per family. Another survey included diverse eudicots (red vertical bar): one basal eudicot, three asterid, three fabid, and two malvid orders were represented by one species per order, in addition to the Brassicales plants. The third survey covered the major flowering plant groups other than the eudicot group: one monocot, two magnoliid, and two basal angiosperm orders. The plants used in the study, their binomial nomenclatures, their taxonomic descriptions, and their sources are listed in Table S3.

### The SA biosynthetic pathways switched from PAL-main to ICS-main within the order Brassicales

We surveyed multiple plant species and a mutant in the order Brassicales since this taxonomic order was known to contain plant species with the functional ICS pathway [16]. Fig 4 contains the three types of plots for selected plant samples. The corresponding plots for all modeled samples are provided in Fig S2. Fig 4 shows that the data from the plant samples were all well-modeled.

**Fig 4.**
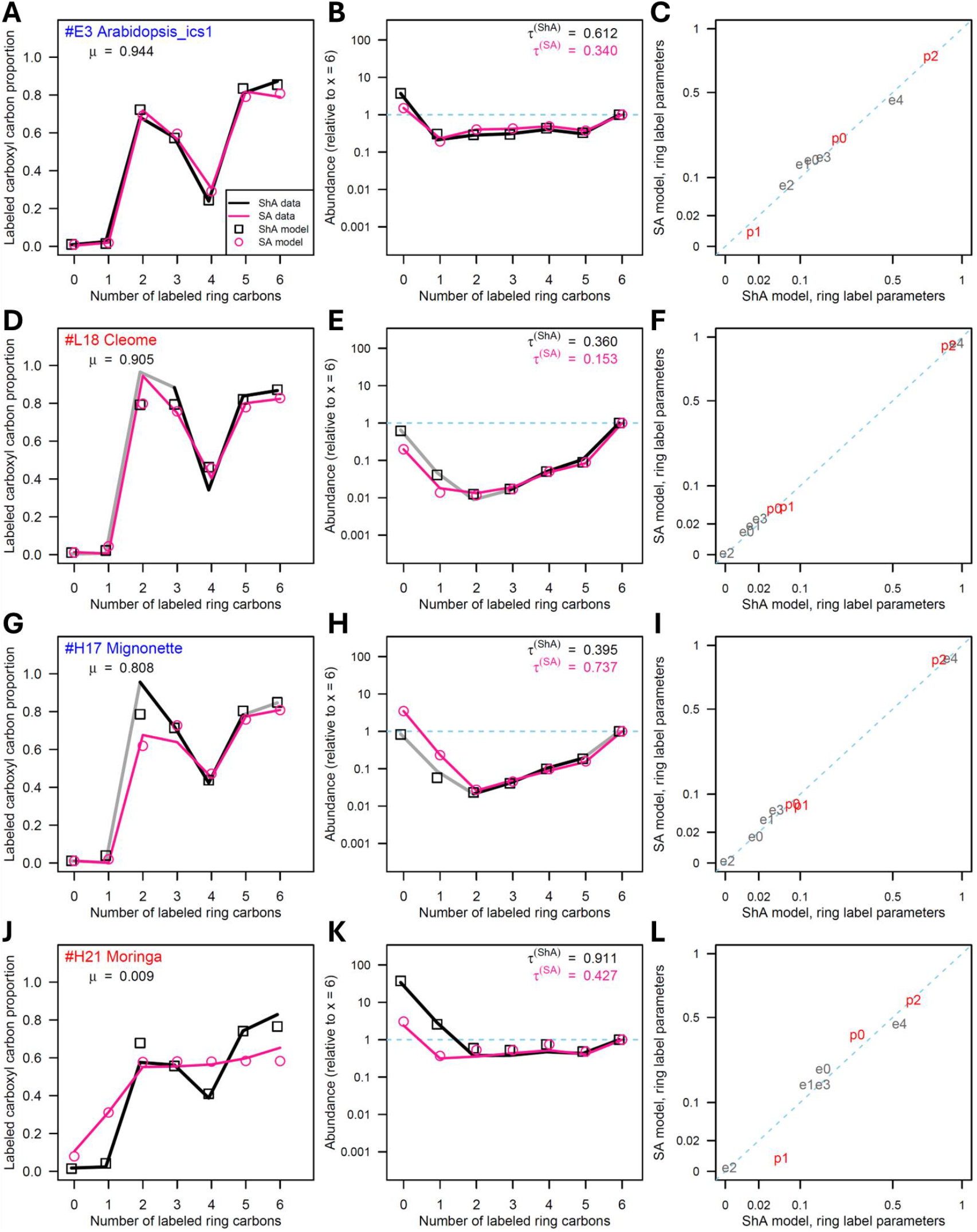
The main pathway switched from the PAL pathway to the ICS pathway within the order Brassicales. The carboxyl carbon labeling profile (A, D, G, J), the ring carbon labeling profile (B, E, H, K), and the ring carbon labeling parameter plot (C, F, I, L) for each of four samples from Arabidopsis *ics1* (A-C), cleome (D-F), mignonette (G-I), and moringa (J-L) are shown. See Fig 2 for details of the three types of plots.

We analyzed an Arabidopsis *ics1* loss-of-function mutant as an example of a major ICS pathway deficiency in Arabidopsis (Figs 4A to 4C). *ICS1* is the major, pathogen-inducible gene of the two Arabidopsis *ICS* genes. Arabidopsis *ics1* accumulated SA at about 10% of Arabidopsis Col-0 (wild type) (Fig 3D), confirming a previous report [29]. Despite a major deficiency in the ICS pathway, the proportion of the ICS pathway contribution was higher in *ics1* than in Col-0 as their *μ* estimates were 0.944 and 0.672, respectively (Fig 4A and 2A). The other ICS gene *ICS2*, which is lowly, constitutively expressed, is intact in the *ics1* mutant, which explains the ICS pathway contribution in the mutant. The *μ* difference between *ics1* and Col-0 was highly significant across multiple biological replicates (*p* = 2.5 × 10^−8^, post hoc, two-sided; *μ* = 0.914±0.055 and *μ* = 0.712±0.030, respectively; Fig 3B). This higher ICS pathway contribution of *ics1* was also visually evident in the carboxyl carbon labeling profiles: the SA profile more closely tracks the ShA profile in *ics1* than in Col-0 (Figs 2A and 4A). In particular, the increased proportion in SA compared to ShA at *x* = 1, which was evident in Col-0, was lost in *ics1*. Generally, no obvious SA proportion increase at *x* = 1 does not necessarily mean very little or no PAL pathway contribution (i.e., the inverse of a true statement is not necessarily true). For example, the profiles of tomato, a PAL-main plant, do not show much proportion increase at *x* = 1 (Fig 2D), likely because the labeling kinetics from PEP #1 to ShA are relatively fast in tomato. However, in the case of *ics1* compared to Col-0, it is unlikely that the *ics1* mutation strongly affects the labeling kinetics before chorismate in the ShA pathway. Thus, it is likely that no obvious proportion increase at *x* = 1 in *ics1* indeed indicates very little contribution from the PAL pathway. This observation of the decreased proportion of the PAL pathway contribution in *ics1* compared to Col-0 is consistent with the positive feedback interpretation: when the amount of SA synthesized through the ICS pathway was reduced by the *ics1* mutation, the decreased positive feedback caused the amount of SA synthesized through the PAL pathway to be even more reduced by proportion (Fig S1). The 90% reduction in the accumulated SA in *ics1* is an additive effect of the decreased SA synthesis through both the ICS and PAL pathways. Chinese white cabbage and stonecress, as well as Arabidopsis, belong to the family Brassicaceae. Stonecress represents the most basal extant genus in the family [30]. All these Brassicaceae species were ICS-main (Fig 3B).

Cleome (family Cleomaceae), mignonette (family Resedaceae), and moringa (family Moringaceae) belong to the order Brassicales but do not belong to the family Brassicaceae (Fig 3A). Cleome and mignonette were ICS-main, and moringa was PAL-main (Fig 3B). These observations are of high confidence as the ICS-PAL modeling worked well for these species (Figs 4D to 4L and Fig S2). Since the divergence time of moringa was earlier than those of mignonette, cleome, and the Brassicaceae species (Fig 3A), these observations with Brassicales species strongly suggest that the main SA biosynthetic pathway switched from the PAL pathway to the ICS pathway within the order Brassicales.

To narrow the time interval when the switch of the pathways occurred in the evolution of the order Brassicales, we attempted to collect data from species belonging to other basal Brassicales families, namely nasturtium (family Tropaeolaceae), papaya (family Caricaceae), poached egg plant (family Limnanthaceae), and saltwort (family Bataceae) (Fig 3A). However, the levels of labeled SA, i.e., the amounts of newly synthesized SA or the approximate SA synthesis rate averaged during the labeling time, in these plants were extremely low (1/1000 or lower compared to Arabidopsis Col-0, Fig 3D), and we were not able to model the pathway contribution using the data from these plants. Note that our ICS-PAL model approach requires sufficiently accurate data for most of the data points shown in both the carboxyl carbon and the ring carbon labeling profiles. Since the levels of labeled ShA were not extremely low, except for papaya, labeling of compounds in the ShA pathway in general was not an issue: the newly synthesized SA amounts were truly low in these plants at least under our experimental conditions. The newly synthesized SA amount in mignonette was also relatively low (Fig 3D), and we were able to estimate the pathway contributions for only two of the 18 samples we prepared from mignonette (the numbers under “n” and “(n1)” on the right flank of Fig 3B).

### The MRCA of eudicots and monocots was likely PAL-main

We included more diverse eudicots in the analysis to investigate the main SA biosynthetic pathways in eudicots in general. Although we analyzed a very diverse panel of eudicots (Fig 3A), we found that they were all PAL-main (Fig 3B). Except for basil (order Lamiales), the *μ* estimates were not significantly different from 0 after multiple tests correction. Thus, these non-Brassicales eudicots almost exclusively use the PAL pathway. This contrasts with the ICS-main Brassicales plants, such as Arabidopsis Col-0, that use the ICS pathway mainly but use the PAL pathway as well. Okra in the order Malvales, which is the sister order of Brassicales (Fig 3A), is PAL-main. This observation is also consistent with the notion that the main pathway switched from the PAL to the ICS pathway within the order Brassicales. Since diverse eudicots, except Brassicales plants, were PAL-main, it is most likely that the MRCA of all eudicots was PAL-main. In addition, since the monocot, rice, is also PAL-main (Fig 3B), the MRCA of eudicots and monocots was likely PAL-main as well.

### Basal angiosperms and magnoliids mainly use a non-ICS/PAL pathway(s)

To broadly cover extant flowering plants, we included two species of basal angiosperms, Amborella and yellow anise, and two species of magnoliids, black pepper and soursop, in our analysis (Fig 3A). SA biosynthesis in these plants appeared to be very different. First, the carboxyl carbon labeling profiles of these plants did not have SA data profile patterns that could be explained by combinations of the ICS and PAL pathways (pink lines in Figs 5A, 5D, 5G, and 5J). They had a pattern in which the labeled carboxyl carbon proportion gradually increases as the number of labeled ring carbons increases. In some cases, it peaked at *x* = 5 (Figs 5D and 5J). Second, in many cases, the SA data values were lower than the ShA data values where 1 < *x* < 6 in the ring carbon labeling profiles (Figs 5H and 5K). This trend in the ring carbon labeling profiles can be explained by two mutually non-exclusive possibilities: (1) there were two or more metabolically active ShA pools with different ring carbon labeling kinetics in the sample, and a ShA pool(s) with labeling kinetics faster than the weighted average of all the metabolically active ShA pools was mainly used for SA synthesis through the ICS or PAL pathways, while the actual data represent the weighted average of all; (2) SA was substantially synthesized via a non-ICS/PAL pathway(s). Although some of these plants have very large metabolically inactive ShA pools (ShA values where *x* ≤ 1 in Figs 5E and 5K; *τ*^(*ShA*)^ in Figs 5F and 5L; plants in the genus *Illicium*, which contains yellow anise, are known to accumulate high levels of ShA [31]), it is unlikely these plants have multiple metabolically active ShA pools with diverse labeling kinetics. It is more likely that these plants use a non-ICS/PAL pathway(s). Third, if the ICS and/or PAL pathways are assumed, then the ring carbon labeling parameters were very different between ShA and SA (Figs 5C, 5F, 5I, and 5L). In many cases for SA, the e1 value is as high as the e4 value, and/or the p1 value is substantial compared to the p2 value (Figs 5C, 5F, and 5L). These are highly unlikely if the ring of SA was mainly synthesized through the ShA pathway (Text S1, section II.3 for e1 and e4; Fig 1A for p1 and p2). For these reasons, we conclude that these basal angiosperm and magnoliid species mainly use a non-ICS/PAL pathway(s) for SA biosynthesis (“non-ICS/PAL main”). Since we are not aware of pathways downstream of the ShA pathway that can synthesize SA with a carboxyl carbon that originates from a source other than 1-C of PEP #1 and 3-C of PEP #2 (which can be modeled by the ICS-PAL model) and since the ring of SA was unlikely to be made mainly via the ShA pathway, the main pathway is most likely a non-ShA pathway.

**Fig 5.**
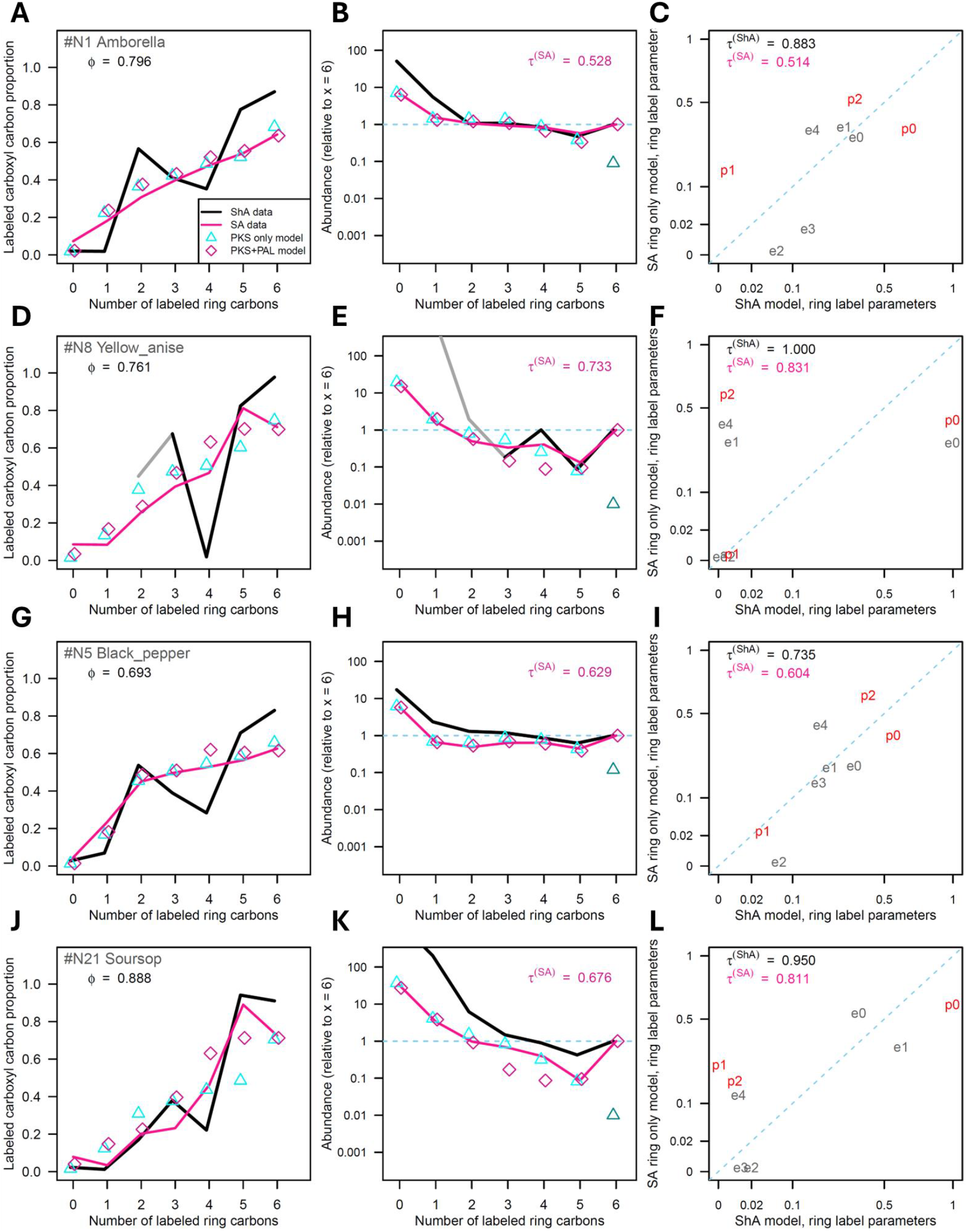
Basal angiosperms and magnoliids mainly use a non-ICS/PAL pathway. The carboxyl carbon labeling profile (A, D, G, J), the ring carbon labeling profile (B, E, H, K), and the ring carbon labeling parameter plot (C, F, I, L) for each of four samples from Amborella (A-C), yellow anise (D-F), black pepper (G-I) and soursop (J-L) are shown. See Fig 2 for details of the three types of plots, except that: in the carboxyl and ring carbon labeling profiles in this figure that have SA estimates from the PKS model (cyan triangle) and the PKS-PAL model (purple diamond) instead of estimates from the ICS-PAL model; the *ϕ* values are the estimated proportion of the PKS pathway contribution in the PKS-PAL model; *τ*^(*SA*)^ in the ring carbon labeling profiles are the preexisting and remaining SA estimates by the PKS-PAL model. *τ*^(*ShA*)^ and *τ*^(*SA*)^ in the ring carbon labeling parameter plots are estimates from the ICS-PAL model fit to ShA and SA data separately. In F, *τ*^(*ShA*)^ is extremely large, and the ShA values (*x*-axis) are unreliable.

The only non-ShA metabolic pathways that we are aware of can synthesize the benzene ring *de novo* in plants are those involving type III polyketide synthases (PKSs), such as chalcone synthase [32]. Among the classes of compounds synthesized via type III PKSs, alkylresorcylates, such as orsellinate in oleander [33] and olivetolate in cannabis [34], attracted our attention for their structural similarity to SA (Fig 1D). In this pathway, a tetraketide intermediate is synthesized by a type III PKS from one molecule of “starter” acyl-CoA, which is specific to each alkylresorcylate (e.g., acetyl-CoA for orsellinate), and three molecules of malonoyl-CoA, and the intermediate is cyclized via C2-C7 aldol condensation to form the alkylresorcylate [32]. We made a PKS model with an assumption that SA was made from an alkylresorcylate by substitutions of 4-OH and the side chain with hydrogens. In the PKS model, the SA carbon labeling state was modeled by the carbon labeling states of the starter acyl-CoA and malonyl-CoA (4 independent carbon labeling state parameters in Fig 1D and one parameter for the preexisting and remaining SA proportion, *τ*^(*SA*)^, were fit to 13 independent observations; Text S2 for details). Unlike the ICS-PAL model, the PKS model was able to model the increasing trend of the SA carboxyl carbon labeling profiles (cyan triangles in Figs 5A, 5D, 5G, and 5J) although the model was not able to generate the profile peaks at *x* = 5, observed in some samples (shortcoming #1; Figs 5D and 5J). The model fits the ring carbon labeling profiles well (cyan triangles in Figs 5B, 5E, 5H, and 5K), except for *x* = 6 (shortcoming #2; turquoise triangle).

We thought that a minor contribution of the PAL pathway in addition to the major contribution of the PKS pathway would be a reasonable assumption because the MRCA of eudicots and monocots was likely PAL-main. The SA only data (13 independent observations) did not have many degrees of freedom left to fit the parameters of the full PAL model, in addition to the PKS model parameters and the proportion parameter of the PKS pathway contribution. Note that the full PAL model has 7 parameters since *P*^(*SA*)^(*L*_*c*_ = 1|*L*_*p*_ = 2) = *P*^(*SA*)^(*L*_*c*_ = 1|*L*_*p*_ = 1) = *P*^(*SA*)^(*L*_*c*_ = 1|*L*_*p*_ = 0) for 100% PAL pathway contribution. To avoid this situation with very low degrees of freedom, we decided to use the parameter values of the #E14 tomato ICS-PAL model (Figs 2D to 2F) as the base and made *P*^(*SA*)^(*L*_*c*_ = 1|*L*_*p*_ = 2) and the proportion of the PKS pathway contribution, *ϕ*, the only additional parameters in this PKS-PAL model (total 7 parameters). The details of the PKS-PAL model and the R script used are provided in Text S2 and Data S2. The corresponding plots and the fitted model parameter values for all modeled samples are provided in Fig S3 and Table S4.

The PKS-PAL model fits well to the carboxyl carbon and ring carbon labeling profiles in general (purple diamonds in Figs 5A, 5B, 5D, 5E, 5G, 5H, 5J, 5K) although the fits to the carboxyl carbon profile peaks at *x* = 5 in Figs 5D and 5J (shortcoming #1) are still somewhat limited. If we fit other PAL model parameters for these samples, instead of using those from the ICS-PAL model for #E14 tomato, these might be further improved. Note that the proportion estimates of the PKS pathway contributions, *ϕ*, are larger than 0.5 (Figs 5A, 5D, 5G, and 5J): the PKS pathway is the main pathway. With this general success in fitting of the PKS-PAL model to the basal angiosperm and magnoliid plant sample data, we conclude that the non-ICS/PAL main pathway mainly used in these plants is likely a pathway involving a type III PKS and that there is likely a minor contribution from the PAL pathway.

## DISCUSSION

We developed a straightforward approach to quantify the proportion of the ICS pathway contribution to SA synthesis in plant leaves when SA is synthesized via either the ICS or PAL pathways. Applying this approach across diverse plant species, we concluded: (1) The MRCA of eudicots and monocots was likely PAL-main; (2) A switch from PAL-main to ICS-main occurred within the order Brassicales; (3) Basal angiosperms and magnoliids mainly use a non-ICS/PAL pathway, which possibly involves a type III PKS.

### The main SA biosynthetic pathways for flowering plants changed drastically more than once during their evolution

Biosynthetic pathways are typically conserved once established and maintained by selection on their end products, because pathway-level function depends on multiple enzymatic steps that may be difficult to favor individually during stepwise evolution [35]. Here, we observed the main biosynthetic pathways for the crucial plant immunity hormone SA drastically changed twice during the evolution of flowering plants: (1) from a non-ICS/PAL pathway to the PAL pathway around the time of the monocot divergence (approximately 140-150 Mya [36]); (2) from the PAL pathway to the ICS pathway within the order Brassicales (approximately 70 to 80 Mya, see below). These changes occurred in a relatively short span of time compared to the likely origin of the green plants (Viridiplantae) more than 1500 Mya [37]. The pathway changes were drastic. Although both the ICS and PAL pathways are downstream of the ShA pathway and share the same ring carbon origins in the final product of SA, the pathways bifurcate downstream of chorismate and have distinct origins of the carboxyl carbon in SA. The non-ICS/PAL pathway is most likely a non-ShA pathway, and both the ring carbons and the carboxyl carbon of SA have origins distinct from the ICS and PAL pathways. There must be an evolutionary reason(s) that such drastic biosynthetic changes occurred more than once in a relatively short time span - i.e., it was not a very rare event that occurred just once by chance. We think that strong pressures from rapidly evolving biotic environments were the driver of such drastic changes in the immunity hormone biosynthesis. For example, a pathogen effector targeting one of the pathway enzymes may have arisen and made the existing main SA biosynthetic pathway ineffective.

### A type III PKS pathway is the best candidate for the non-ICS/PAL pathway in basal angiosperms and magnoliids

The gradually increasing SA carboxyl carbon labeling profiles with basal angiosperms and magnoliids (pink lines in Figs 5A, 5D, 5G, and 5J) cannot be explained by a combination of the ICS and PAL pathway. When we modeled SA labeling assuming an alkylresorcylate synthesized via a type III PKS as an intermediate for SA synthesis, the PKS model was able to explain the increasing SA carboxyl carbon labeling profiles (cyan triangles in Figs 5A, 5D, 5G, and 5J). By assuming a minor contribution of the PAL pathway, the PKS-PAL model was also well fit to the ring carbon labeling profiles (purple diamonds in Figs 5B, 5E, 5H, and 5K). In addition, the PKS-PAL model was able to generate a carboxyl carbon labeling profile with a peak at *x* = 5 although the fitted peak proportion was not as high as that in the data (purple diamonds in Figs. 5D and 5J). Thus, we think that the main SA biosynthetic pathway in basal angiosperms and magnoliids is likely a type III PKS pathway.

Type III PKS gene candidates for SA biosynthesis can be identified by homology to the chalcone synthase gene [38]. If there is a basal angiosperm or magnoliid species that is amenable to transient gene knock-down, such as virus-induced gene silencing, these candidate genes could be silenced, and the type III PKS gene for SA biosynthesis could be identified if its silencing substantially reduced UV-induced SA synthesis. Once the type III PKS gene was identified, the starter acyl-CoA for the enzyme could be identified *in vitro* [32]. Once the starter acyl-CoA was identified, the alkylresorcylate product would be known, which would narrow down the search of the two downstream ring substitution reactions.

### Rice may partially use a type III PKS pathway

Although we speculated that the MRCA of the monocots and the eudicots was likely PAL-main based on the assumption that the ICS and PAL pathways are the only possible pathways, the discovery of the non-ICS/PAL pathway in basal angiosperms and magnoliids challenges this speculation. When the SA data carboxyl carbon labeling profiles of rice and black pepper (pink lines in Figs 2G, 2J, and 5G) are compared, they share an evident trend: the proportion of the labeled carboxyl carbon gradually increases even where *x* > 1, which cannot be explained by the ICS-PAL model but is a feature of the PKS model. This shared trend suggests that SA synthesis in rice could be a mixture of the PAL and non-ICS/PAL pathways. If this is the case, switching to truly PAL-main may have occurred after the divergence of monocots. Since the carboxyl carbon labeling profiles of the basal eudicot columbine do not show evident increase where *x* > 1 (Fig S2), the MRCA of all extant eudicots was probably truly PAL-main.

If rice indeed uses the PKS pathway substantially, the experimental approach discussed for basal angiosperms and magnoliids above can be employed in rice. Since rice is an established experimental system for genetic manipulation, using rice might be the shortest path to identify an example type III PKS for SA biosynthesis.

### Switching of main SA biosynthetic pathways may have occurred more than twice in flowering plants

Switching of the main SA biosynthetic pathways occurred more than once in flowering plants. In one case, it occurred within a single taxonomic order of Brassicales. There are 43 other extant eudicot orders and 11 extant monocot orders [17]. Thus, we think that there is a good chance to discover more switching events if the depth of the plant survey is substantially increased. It should be noted that we only tested leaf tissues under very limited experimental conditions. The proportions of pathway contributions could be different in different tissues and/or under different conditions even within the plant species we have already tested because different tissues and/or different conditions have likely been under different biotic pressures.

We have some ideas about which clades of plants might have undergone switching events. Among all non-Brassicales eudicots surveyed, only basil (order Lamiales) had a *μ* value significantly higher than 0 (Fig 3B; *p* = 1.4 × 10^−6^, post hoc, two-sided). This observation in basil might indicate a precursor of a switching event. Deeply surveying the highly diverse order Lamiales may result in discovery of another switching event.

In Brassicales, it appears that strong reduction of the SA synthesis rate (the amount of the labeled SA) preceded the innovation of the Arabidopsis-type ICS pathway (Figs 3B and 3D). If this is a common case, surveying clades that contain plants with very low SA synthesis rates may lead to discovery of other switching events. We found that Setaria ME034V had a very low SA synthesis rate while rice had a high rate (Fig 3D). The family Poaceae, which includes these species, may be a good clade for a deep survey. We dropped *Nicotiana benthamiana* from our study at a very early stage because it had a very low SA synthesis rate after UV-treatment. It has been reported that *N. benthamiana* does not accumulate as much SA as *N. tabacum* does [39]. Thus, the genus *Nicotiana* may be another good clade for a survey. If selecting clades for a survey based on members with very low SA synthesis rates in the clades is indeed a good strategy, screening plants broadly for very low SA synthesis rates first would allow targeted deeper surveys of other clades likely to include pathway switch events.

### Evolution of SA biosynthesis in Brassicales may have been shaped by changing biotic environments

We speculate that evolution of basal Brassicales plants (which we define as the Brassicales lineages that diverged before emergence of the core Brassicales) and extensive diversification of the core Brassicales were strongly shaped by their biotic interactions with pathogens and insects. The following are possible scenarios of the Brassicales evolution we propose. The dates for Brassicales evolution below are according to [40].

We discovered that very low SA synthesis rates were common among extant basal Brassicales. Among the five basal Brassicales families we surveyed (Tropaeolaceae, Moringaceae, Caricaceae, Limnanthaceae, and Bataceae), only Moringaceae was able to synthesize a high level of SA mainly via the PAL pathway (Figs 3B, 3D, and 4J to 4L). With their phylogenetic relationships (Fig 3A), currently we cannot determine which of the two possible cases occurred: the MRCA of all Brassicales (~103 Mya) had already lost the high SA synthesis rate (pink “X” in Fig 3A), and somehow the Moringaceae lineage regained the high rate (pink star in Fig 3A); the MRCA of all Brassicales had a high SA synthesis rate, Moringaceae inherited it, and the other lineages lost the high SA synthesis rate in at least three independent events (three gray “X”s in Fig 3A). Structural and expression analysis of the genes involved in SA synthesis via the PAL pathway and in its regulation in extant basal Brassicales plants will reveal when and how the high SA synthesis rate was lost, which will enable us to determine which of the two cases in fact occurred.

In either case, a very low SA synthesis rate must have somehow benefited the ancestors of these basal Brassicales. There are two possible cases that are not mutually exclusive. First, an environment that would benefit from a very low SA synthesis rate is one where insects (and/or necrotrophic pathogens) are much more problematic to the plant than biotrophic pathogens. In such an environment, a very low SA synthesis rate would be favored because SA signaling suppresses JA signaling, and JA induces immune responses that are effective against insects and necrotrophic pathogens [11]. With a much lower SA synthesis rate, plants could have induced faster and stronger immune responses against insects. Second, a major pathogen(s) of those days in the area may have evolved an effector(s) that targets the PAL pathway. If so, having the PAL pathway as the major SA synthetic pathway would have become ineffective for inducing SA-mediated immune responses, and consequently the plant may have lost a PAL pathway function for an energy/resource efficiency reason.

Note that since the immune signaling subnetworks back up one another to some extent, plants have some level of remaining immunity against biotrophic pathogens even without SA, and thus, such plants are not completely defenseless against biotrophic pathogens [7]. The ancestors of basal Brassicales were likely in the Americas, as that is where extant basal Brassicales are present these days, with rare exceptions like Moringaceae in the Old World [40]. Probably at least some parts of the Americas have retained similar environments since those days (approximately 80-100 Mya) so that the basal Brassicales plants with very low SA synthesis rates were able to survive to date.

After the divergence of the Limnanthaceae lineage, indolic glucosinolates, which form a powerful chemical defense against insects, were innovated approximately 80 Mya (“Indolic G” arrow in Fig 3A) [41]. We think that this event shifted the balance sheet of SA signaling: having a higher SA synthesis rate and stronger inducible immunity against biotrophic pathogens was no longer a disadvantage as plants could afford to have weaker JA signaling – plants can have strong immunity against both insects and biotrophic pathogens. This possible SA-signaling-favored period lasted until approximately 70 Mya when the Pierina butterfly evolved to have indolic glucosinolate detoxifying enzyme genes and began rapid diversification [41]. Interestingly, the Arabidopsis-type ICS pathway for SA synthesis was innovated in an ancestor of the core Brassicales (blue-green vertical bar in Fig 3A), which includes Resedaceae, Cleomaceae, and Brassicaceae, approximately in this 10-million-year potentially-SA-signaling-favored period. The innovation was likely initiated by neofunctionalization of the *EDS5* IC transporter gene after tandem duplication of its ancestral gene (“EDS5” lightning symbol in Fig 3A) [16]. If a pathogen(s) with a PAL-pathway-inhibiting effector gene(s) was still presnt at that time (see above), it makes sense that a new SA biosynthetic pathway was innovated instead of restoring the impaired PAL pathway. Strong immunity against insects and regained stronger immunity against biotrophic pathogens probably contributed to the rapid radiation of the core Brassicales into different environments with different pathogens and insects, in all continents except Antarctica [40]. Moringaceae, a rare example of a basal Brassicales family native to the Old World, has a high SA synthesis rate, suggesting that a high SA synthesis rate and consequential strong immunity against biotrophic pathogens were crucial for surviving in the Old World to date.

Although mignonette (Resedaceae) can synthesize SA via the Arabidopsis-type ICS pathway (Figs 3B and 4G to 4I), its SA synthesis rate was low compared to cleome (Cleomaceae) and the Brassicaceae species (Fig 3D). It would be interesting to compare the SA synthesis rates of plant species between the sister clades of the core Brassicales: the clade containing the GRFT clade, which includes Resedaceae, and the Brassicaceae-Cleomaceae-Capparaceae (BCC) clade. If the SA synthesis rate is consistently higher in the BCC clade, further strengthening of SA signaling might have been enabled by innovation of another class of glucosinolates, methionine-derived glucosinolates, in an ancestor of the BCC clade (“Met G” arrow in Fig 3A) [41]: just like the innovation of indolic glucosinolates may have allowed the innovation of the Arabidopsis-type ICS pathway in an ancestor common to both clades. While the BCC clade plants have orthologs of the IC-glutamate conjugating enzyme gene *PBS3*, the transcriptomes of mignonette, saltwort, and poached egg plant lacked *PBS3* orthologs [16]. Thus, this possible SA synthesis rate increase in a BCC clade ancestor appears to have coincided with the innovation of PBS3, which would explain the large increase of the SA synthesis rate through the ICS pathway in this clade. Note that ICS1 and EDS5 are absolutely required for SA synthesis through the reconstituted ICS pathway in *N. benthamiana* while inclusion of PBS3 is not required but strongly enhances SA synthesis (Fig 2B of [42]). In any case, methionine-derived glucosinolates, the high SA synthesis rate, and the consequential very strong immunity against both insects and biotrophic pathogens may have contributed to the success of the BCC clade (~4500 extant species [40]).

### SA biosynthetic pathways could be even more diverse than the three discussed here

We have been discussing three pathways, the ICS, PAL, and non-ICS/PAL pathways because these are distinguished by our approach. However, lower-level diversities within the three pathways, which would not be detected by our approach, are also possible. For example, the pathways downstream of *t*-cinnamate, which is the product of PAL, in the PAL pathway may be diverse. Recently, downstream of *t*-cinnamate, the benzyl-benzoate (BB) pathway was found to be the major pathway in several flowering plants, including rice and *N. benthamiana* [43–45]. Before these studies, it was thought that 2-hydroxylation of benzoate was the likely last step of SA synthesis in the PAL pathway [46]. In the BB pathway, benzoate was made into the benzyl ester (which is BB) first, then the benzoate part of the benzyl ester was 2-hydroxylated, and finally the benzyl group was removed by hydrolysis of the ester to form SA. Why are this esterification and its reversal included in the BB pathway? – it appears to be a metabolic detour and a waste of energy. It is plausible that a simpler pathway without benzyl esterification could have been used earlier and that the simpler pathway may have been targeted by a pathogen effector(s) and compromised, which may have generated the selection pressure for the innovation of the BB pathway. In fact, two of the enzyme genes that compose the BB pathway, *OSD2* and *OSD4*, are missing in non-seed land plants, such as a fern and a lycophyte [44], while such non-seed land plants are known to accumulate SA [47, 48]. Thus, non-seed land plants may have a different pathway downstream of *t*-cinnamate in the PAL pathway or mainly use a non-ICS/PAL pathway.

The Arabidopsis lineage once lost much of its SA synthesis ability via the PAL pathway, resulting in the very low SA synthesis rates observed in poached egg plant and saltwort (Fig 3D). How is it possible for Arabidopsis Col-0 to synthesize ~30% of total SA via the PAL pathway (Fig 3B), which is an orders of magnitude higher amount (Fig 3D)? This is remarkable when Arabidopsis Col-0 is missing *OSD3* and *OSD4* [44]: i.e., Arabidopsis does not have the complete BB pathway. Since Arabidopsis Col-0 still has the other enzyme genes for the BB pathway [44], it is possible that this lineage evolved alternatives to the *OSD3* and *OSD4* genes, which encode a hydroxylase of a benzene ring and an ester hydrolase, respectively. With some levels of substrate promiscuity, it is plausible that some other genes encoding enzymes of these common classes may have initially functioned to some extent and have improved through selection. Rice knockout mutants of *osd3* or *osd4* can still accumulate low but detectable levels of SA while an *osd2* mutant accumulates much lower levels of SA (Figs 1i, 1l, and 1o of [44]): i.e., rice has low efficiency alternative enzymes for OSD3 and OSD4. Probably many other plants also have such low efficiency alternative enzymes. If this notion of low efficiency alternative enzymes for OSD3 and OSD4 was true in an ancestor of Arabidopsis as well, the scenario of having started with similar enzymes with substrate promiscuity is likely. It should be noted that mignonette (Resedaceae) had the Arabidopsis-type ICS pathway, which synthesized ~75% of SA, and the SA synthesis rate was still lower compared to the plant species that diverged later, such as cleome and Brassicaceae species (Fig 3D). It is likely that the MRCA of mignonette and Arabidopsis was like mignonette: the SA synthesis rate was low and ~25% of SA was synthesized via the PAL pathway. If this is the case, the efficacy of the PAL pathway in addition to that of the ICS pathway must have improved through selection between the divergence times of Resedaceae and Cleomaceae.

The reason we have been specifying the ICS pathway in Brassicales as “Arabidopsis-type” is because we could imagine different types of ICS pathway. For example, since chorismate is present in the cytosol as well as the plastid [49], it could be possible to evolve an ICS pathway solely in the cytosol if the *ICS* gene is duplicated and the protein encoded by one of the duplicated genes is evolved to be targeted to the cytosol. This might occur if a pathogen effector targeted the plastidic ICS. In such a scenario, a plastidic IC transporter, like EDS5, would not be required. Thus, if another lineage that evolved an ICS pathway independently is found, it may have a different type of ICS pathway. If it was this cytosolic type, it could be identified by cytosol-targeting of an ICS.

### How can we genetically test the SA biosynthetic pathways under strong positive feedback from SA?

We have pointed out the fundamental issue associated with pathway-perturbation studies of systems with feedback regulation. If SA is synthesized by two pathways and both are regulated by positive feedback from the final product of SA, genetically disabling one pathway decreases the output of the second pathway as well. Consequently, the contribution of the second pathway is underestimated or undetectable (Fig S1). We observed that the proportion of the PAL pathway contribution was decreased by a major deficiency in the ICS pathway in Arabidopsis (Col-0 and *ics1* in Figs 2A, 2D, and 3B), and we postulated that this was because of decreased positive feedback to the PAL pathway. How could this notion be experimentally tested? If the objective is to prove the existence of a second pathway, an artificial application of a functional analog of SA, such as benzothiadiazole (BTH) [21], would be a solution. BTH would provide the effect of SA positive feedback, but newly synthesized SA could be distinguished from BTH analytically. For example, if an Arabidopsis *ics1 ics2* plant, which accumulates only ~4% of SA compared to wild type after UV treatment [22], accumulates significantly more SA upon BTH treatment (in addition to UV treatment), it would prove the existence of a non-ICS pathway. In combination with our approach, it could be determined whether the increased amount of SA produced with BTH treatment is indeed synthesized through the PAL pathway. Furthermore, if the non-ICS pathway is indeed the PAL pathway, comparison of the SA levels after BTH treatment in the *ics1 ics2* double mutant and the *ics1 ics2 osd2* triple mutant would reveal whether the PAL pathway uses the modified BB pathway, in which alternative enzymes for OSD3 and OSD4 are used (see the immediately previous section).

In the unlikely situation that the inducer had to be SA instead of BTH, metabolically labeling newly synthesized SA using [^13^C_6_]glucose, as in our approach, could distinguish newly synthesized SA from SA used for induction, making this type of study feasible.

We have also speculated that a non-ICS/PAL pathway may have a partial contribution to SA synthesis in rice based on the shape of the carboxyl carbon labeling profiles (Figs 2G, 2J, and S1). Since SA accumulation is very low in the rice *osd2* mutant [44], the non-ICS/PAL pathway might be positively regulated by SA. The existence of a non-ICS/PAL pathway could be tested by treating the rice *osd2* mutant with BTH.

The approach we proposed here, BTH-treatment of a mutant with a defect in one pathway, aims to detect the existence of a second pathway. To determine the contribution proportion of each pathway, the contributions need to be measured in intact plant systems, such as by using our approach, when the systems are regulated by feedback.

### Do some plants, such as rice, have a large SA pool that is inactive for immune signaling?

Rice is known to have a very high basal SA amount and not much SA increase after pathogen challenge [28]. We speculate that the total SA amount in rice may include a substantial pool of signaling-inactive SA for two reasons. First, many of the enzyme genes involved in the BB pathway are highly pathogen-inducible [44]. When an ancestor of rice evolved the inducible transcriptional regulatory system and the rice lineage has maintained it, it does not make sense if the actual signal output is the total SA amount, which is not inducible. Second, although the *OSD1*-overexpression lines showed only a two-fold increase in the total SA amount, the lines exhibited a clear increase in immunity [44]. It is not easy to build a response system that reliably detects a two-fold difference in the signal in a noisy biological environment. These two points could be explained if we assume that a large portion of the preexisting SA amount is signaling-inactive and that the newly synthesized SA amount is inducible and mainly drives the SA signaling response. The newly synthesized SA amount could be obscured by the large amount of preexisting SA in the total SA measured. To test this hypothesis, the newly synthesized SA amount needs to be measured separate from the total SA amount. We developed a simple and efficient SA-metabolic-labeling method, which would suffice for this purpose (e.g., Fig 3D; only an LC-MS, instead of an LC-MS/MS, is needed for this application). One advantage of this method is that by measuring the newly synthesized ShA, the flux of the labeling carbons into the ShA pathway can be assured. Thus, our labeling method will help test this hypothesis of the newly synthesized SA as the major SA signaling driver. We found that the newly synthesized SA amounts in rice and Arabidopsis Col-0 were comparable after a UV-C treatment (Fig 3D), and thus we think that the newly synthesized SA amount in rice is likely inducible. We observed a very high ratio of total SA vs. newly synthesized SA in Setaria ME034V as well (newly synthesized SA was undetectable; Fig 3D). The large signaling-inactive SA pool might be a widespread trait in the family Poaceae. Furthermore, if indeed some plants have a substantial signaling-inactive SA pool, the amount of newly synthesized SA could be a better measure of the SA immune signaling function in general than the amount of total SA.

### Can a very low SA synthesis rate be the cause of a high Agrobacterium-mediated transformation efficiency?

We noticed a correlation between plants with very low SA synthesis rates and plants known to be highly (transiently or stably) transformable by Agrobacterium, such as *N. benthamiana* [50] and Setaria ME034V [51]. We previously reported that strongly compromising plant immune signaling highly increased the efficiency of transient transformation by Agrobacterium in Arabidopsis [52]. Although the effect of an *ics1* (*sid2*) mutation alone on transformation efficiency was limited (Fig S3 in [16]), the *ics1* mutant still has quite a high SA synthesis rate on the scale of interest here (Fig 3D). This correlation suggests that strongly reducing the SA synthesis rate could substantially increase the transformation efficiency by Agrobacterium. It is interesting to see whether any of the basal Brassicales plants with very low SA synthesis rates (Fig 3D) have high transient transformation efficiency by Agrobacterium. If this potential causality is generally true, it would have strong application implications. For example, in a crop species, SA biosynthetic pathway mutations generated by CRISPR technology in a transformable variety can be introgressed into elite varieties. More than one pathway may need to be targeted to obtain a very low SA synthesis rate. Once such elite variety mutant lines with very low SA synthesis rate were established, they could be transformed easily for GM or genome editing purposes. With just one backcross to the original elite varieties to remove the SA biosynthesis mutations (and the CRISPR-related transgenes in the case of genome editing), GM or genome-edited lines with elite variety backgrounds would be ready for propagation.

### Concluding remarks

Using [^13^C]-metabolic labeling, LC-MS/MS, and the metabolism-guided statistical modeling of ShA and SA in a plant leaf, we were able to determine the proportions of the ICS and PAL pathway contributions or whether a non-ICS/PAL pathway is mainly used. Using this approach, we observed that major SA biosynthetic pathways in flowering plants drastically changed at least twice, which strongly implies that plants had to use an unconventional evolutionary mechanism to adapt to their rapidly evolving biotic environments. We predict that flowering plants are likely to have more diversity in SA biosynthesis routes and that our approach will help discover such events by expanding the plant species surveyed.

## METHODS

### Plant materials

We purchased seeds or other plant materials from online stores, used plants common in plant research labs, or obtained materials from the University of Minnesota College of Biological Sciences Conservatory. The details of the plant sources are given in Table S3. Seeds of some plants were subjected to stratification before they were sown, and the stratification information is also given in Table S3. Most plants were germinated/grown at 22°C with ~60%RH and a 12-hour light (PAR = 110 μmol/m^2^/s)/12-hour dark cycle in a home-made plant growth chamber designed for Arabidopsis growth [53] before use (Fig S4A). Monocot seedlings were grown at 25°C with ~50%RH and a 12-hour light/12-hour dark cycle for 14 days after sowing. Leaves of basal angiosperms and magnoliids were obtained from the conservatory.

### Labeling with [^13^C]-labeled glucose

A leaf used in a labeling experiment was harvested with a ~1-cm petiole. If the petiole was too short, part of the leaf was cut off to make the petiole length ~1 cm. If a leaf was too large (> 10 cm^2^), it was cut into a leaflet with a ~1-cm main vein extending from it as if it were a petiole (e.g., papaya in Figs S3B and S3C). If leaves were too small multiple leaves were used (e.g., moringa in Figs S3B).

First, the petiole of each leaf was placed into 2 ml water in a well of a 24-well microtiter plate. The plate with leaves was placed in a dark box (a cardboard box lined with aluminum foil), and the box was placed in the growth chamber at 22°C with ~60%RH for ~20 hours. Second, the plate was placed under a handheld 254-nm UV-C lamp (Analytik Jena US, UVP UVG-54) 30-cm below the lamp, for up to 10 minutes. Leaves of some plant species were highly sensitive to UV-C, so they were treated for a shorter time or not UV-C treated at all. The UV-C treatment time for each leaf sample is given in Tables S2 and S4. Third, the leaves were moved to 50 mM [^13^C_6_]glucose (Cambridge Isotope Laboratories, CLM-1396-1) solution (typically 250 μl in a well of a 96-well microtiter plate) and kept under a low light condition on the top shelf of the home-made growth chamber (PAR = 3.5 μmol/m^2^/s, indirect light from the lower shelves; measured with an Apogee DLI-500; Fig S4A) for 8 to 30 hours (the labeling time is given in Tables S2 and S4). Each leaf sample was harvested into a 2-ml pre-weighed screw-capped tube containing two ball bearings, weighed, and flash-frozen with liquid N_2_. The frozen leaf sample in the tube was pulverized into powder using a paint shaker for three minutes twice, and 70% isopropanol was added to the tube (0.5 to 1 ml/0.1 g FW). The tube was vortexed for 10 seconds three times and placed on ice. The tube was centrifuged at 15,000 x g for 30 min at 4°C. The supernatant was collected in a 1.5-ml low-binding tube and kept at −80°C until LC-MS/MS analysis.

### Liquid chromatography-mass spectrometry (LC-MS & MS/MS) analysis

Labeled plant extracts were thawed on ice and concentrated under vacuum with centrifugation (SpeedVac, ThermoFisher) for 1 hour to remove isopropanol. The samples were then subjected to centrifugation to remove particulates in a refrigerated microcentrifuge (16,000 g, 10 minutes, and 4°C), 45 µl of supernatant was transferred to glass inserts in autosampler vials prior to acidification by addition of 5 µl of 5% (v/v) formic acid in water. Samples were analyzed by LC-MS using an Ultimate 3000 LC (with autosampler sample vial block maintained at 4°C) coupled to a Q-Exactive quadrupole-Orbitrap mass spectrometer equipped with a heated electrospray ionization (HESI) source (ThermoFisher Scientific, Waltham MA). Chromatographic separation was accomplished using a sample injection volume of 0.5 μl, and an HSS T3 C18 column (2.1 x 100 mm, 1.8 µm; Waters, Milford MA), with column temperature 40°C and flow rate 0.40 ml/min. The solvent system consisted of solvent A (0.1% formic acid in water) and solvent B (0.1% formic acid in acetonitrile). The 14-min LC gradient was: initial 0.5% B, 2 min 0.5% B, 4 min 15% B, 5 min 25% B, 11 min 35% B, 12 min 98% B, 13 min 98% B, 14 min 0.5% B. For MS1 analysis, data were collected in profile mode at 70k resolution, from *m/z* 100 – 500, and in negative ionization mode with the following HESI settings: desolvation temperature 350°C, spray voltage 3800 V, auxiliary gas flow rate 20, sheath gas flow rate 50, sweep gas flow rate 1, S-Lens RF level 50, and auxiliary gas heater temperature 300°C. MS/MS was performed in profile mode, at 17.5k resolution, and selecting targeted *m/z* values with ±0.5 Da isolation window centered on each isotopologue of labeled SA and ShA using the same HESI settings as for MS1. Elution windows and normalized collision energy (NCE) values were as follows: for ShA: 0-2 min., NCE 10; SA: 6.5-8.5 min, NCE 50). To maximize signal, each significant isotopologue of labeled SA and ShA were progressively isolated for HCD fragmentation over multiple repeated injections for each entire elution window. Data were acquired using Xcalibur™ software version 2.1 (ThermoFisher Scientific) and were centroided and converted from Thermo’s proprietary format (.raw) to an open-source format (.mzML) using MSConvert from the ProteoWizard package [54]. The MS1 data were used to perform a linear regression between each isotopologue and the ^12^C-monoisotopic masses for SA and ShA over their respective elution windows. The slope values for each isotopologue regression provided the relative abundance ratio of that isotopologue intensity to that of the ^12^C-monoisotopic peak yielding the entire isotopologue distribution for each compound relative to its monoisotopic peak intensity. For the MS2 data, the isotope distributions of specific product ions (e.g., [M–H–CO_2_]^−^) in each MS/MS run were used to extract positional labeling information for SA and ShA for each significant isotopologue peak of the molecular ion.

### Estimation of the μ value from LC-MS/MS data

Details of the theoretical basis and implementation of the metabolism-guided ICS-PAL statistical model are provided in Text S1, and the R script using the minpack.lm package is provided as Data S1. For each sample, the *μ* mean estimate and its standard error were calculated by the R script. The sample’s *μ* estimate was deemed valid if it satisfied all the following four conditions: (i) the *μ* standard error is no larger than 0.8; (ii) the *μ* estimate is between −0.1 and 1.1 (not including the boundary values); the number of observations with weights higher than 1/80 is at minimum 17, (i.e., there are not too many bad observations); (iv) the number of observations with absolute values of the residuals that are equal to or larger than log4 and with weights equal to or larger than 0.4 is no larger than 1, (i.e., if the residual is large, it should be associated with an observation with a low weight).

### PKS model and PKS-PAL model

Details of the theoretical basis and implementation of the metabolism-guided PKS and PKS-PAL statistical models are provided in Text S2, and the R script using the minpack.lm package is provided as Data S2.

### Analysis of the μ estimates across plant species

For Fig 3B, 1-way ANOVA with the plants as the fixed effect was applied to the *μ* mean estimates weighted with 1/(standard error of *μ*)^2^ (Data S1 is an R script including generating Figs 3B, 3C, and 3D). The residual distribution of the ANOVA model justified the use of the ANOVA model (Fig S5).

## AUTHOR CONTRIBUTIONS

FK conceived the project, inspired by early, then-unpublished results by KH and KT [16]. FK, EAL, JDC, and ADH designed the research. FK and EAL performed experiments. FK and EAL analyzed data. FK wrote the manuscript with assistance from the other authors.

## ACKNOWLEDGEMENTS

We thank the University of Minnesota, College of Biological Sciences Conservatory and Botanical Collections for basal angiosperm and magnoliids plant materials, The 2Blades Group for monocot plant materials, and Kaylie Richard for critical reading of the manuscript.

## SUPPLEMENTAL INFORMATION

**Fig S1.**
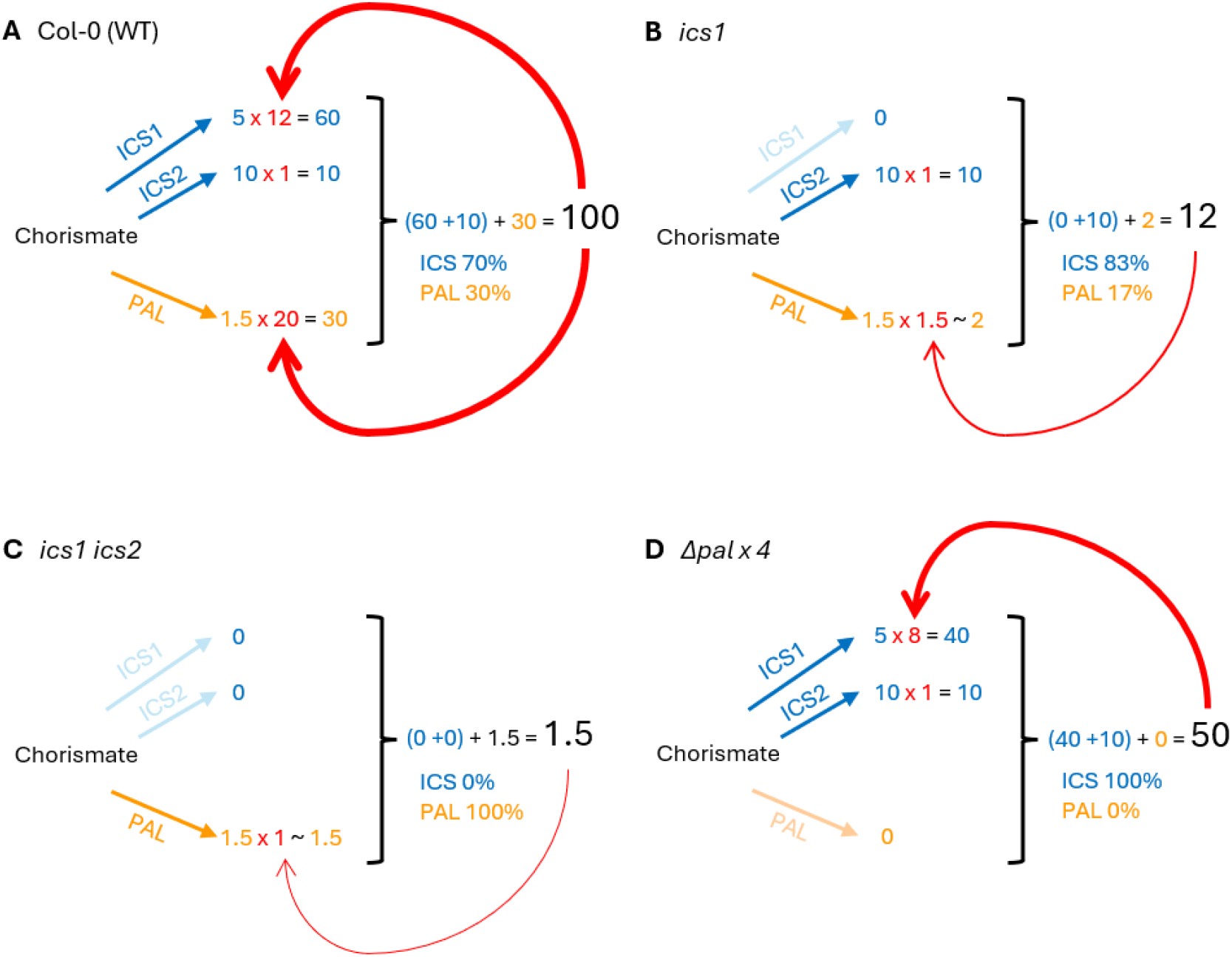
Schematic diagrams of positive feedback on SA synthesis from SA in (A) Col-0 (WT), (B) *ics1*, (C) *ics1 ics2*, and (D) all four *pal* deletions (*Δpal x 4*) of Arabidopsis. The ICS pathway is divided into the ICS1 and ICS2 pathways (blue arrows) since while the *ICS1* mRNA level is SA inducible [20, 21], the *ICS2* mRNA level is constitutive [22]. The PAL pathway is made as a single collective pathway (orange arrow). The values for the basal activities (blue and yellow values at the beginning of multiplication for the ICS and PAL pathways) and the fold values for induction (red values preceded by the multiplication sign, “x”) in SA synthesis by different pathways were arbitrarily chosen for the purpose of explaining the principle of the positive feedback interpretation of the SA biosynthesis pathways, except that the induction fold value for the ICS2 pathway was assumed to be 1 as the *ICS2* mRNA level is constitutive. Red arrows indicate the positive feedback from SA, and their widths represent the strength of the feedback shown by the induction fold values, which is positively correlated to the total SA synthesized amount value (in a black large font, most right in each panel) for each pathway. Note that the feedback strength is not assumed to be linearly correlated. The positive feedback interpretation can qualitatively explain the following observations in these genotypes: (1) The induced SA accumulation level is approximately 10% in *ics1* compared to WT [22]; (2) The induced SA accumulation level in *ics1 ics2* is even lower than *ics1* but not zero [22]; (3) The induced SA accumulation level in *Δpal x 4* is approximately 50% of WT [23]; (4) The contribution of the ICS pathway is approximately 70% in WT (Figs 2A and 3B); (5) The contribution of the ICS pathway is higher in *ics1* than in WT (Figs 4A and 3B).

**Fig S2.** Three types of plots, the carboxyl carbon labeling profile, the ring carbon labeling profile, and the ring carbon labeling parameter plots, for 76 samples, to whose data the ICS-PAL model fit well. See Fig 2 for details of the plots.

**Fig S3.** Three types of plots, the carboxyl carbon labeling profile, the ring carbon labeling profile, and the ring carbon labeling parameter plots, for 8 samples, to whose SA data the PKS-PAL model fit reasonably well, and 3 additional Amborella samples (#N7, N13, and N19) to show their ShA data profiles (most SA data points were unreliable or below detection). See Fig 2 for details of the plots.

**Fig S4.**
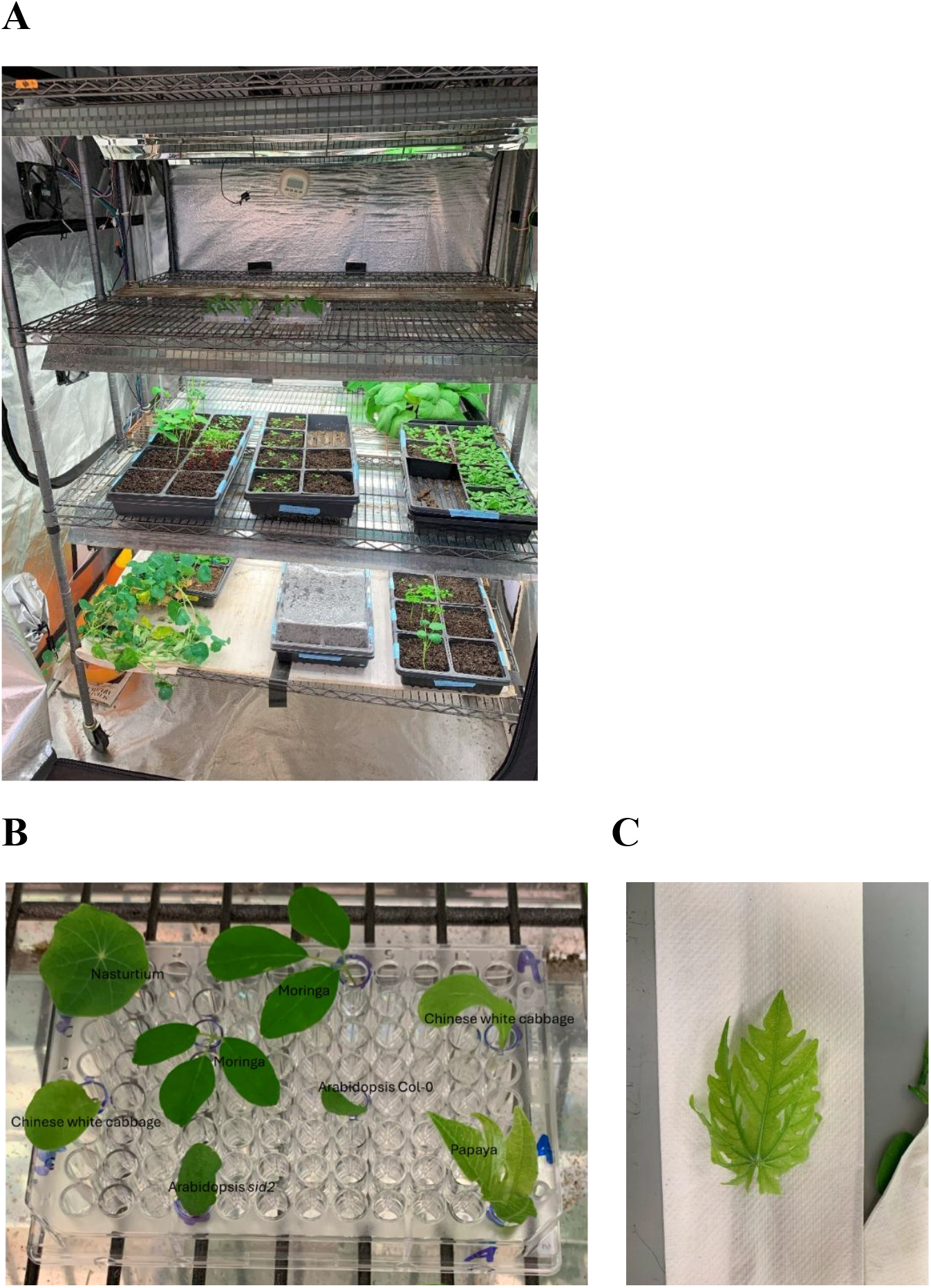
Photos of experimental procedures. (A) A homemade growth chamber. The top shelf had all the lights off and was used for the low light condition during [^13^C]-labeling of leaf samples. The middle and bottom shelves were used to grow plants for the experiments. The shorter edge of the black plant growth flat is ~30 cm. (B) An example of the [^13^C]-labeling experiment (samples #F1 to #F8). This is to show the typical leaf/leaflet sizes used in the experiments. The 96-well plate is 12.5 cm x 8 cm. The papaya leaflet (#F8) in the bottom right corner was prepared from a leaf in C. (C) The papaya leaf, from which the leaflet (#F8) in B was prepared. The width of the paper towel under the leaf is 16 cm.

**Fig S5.**
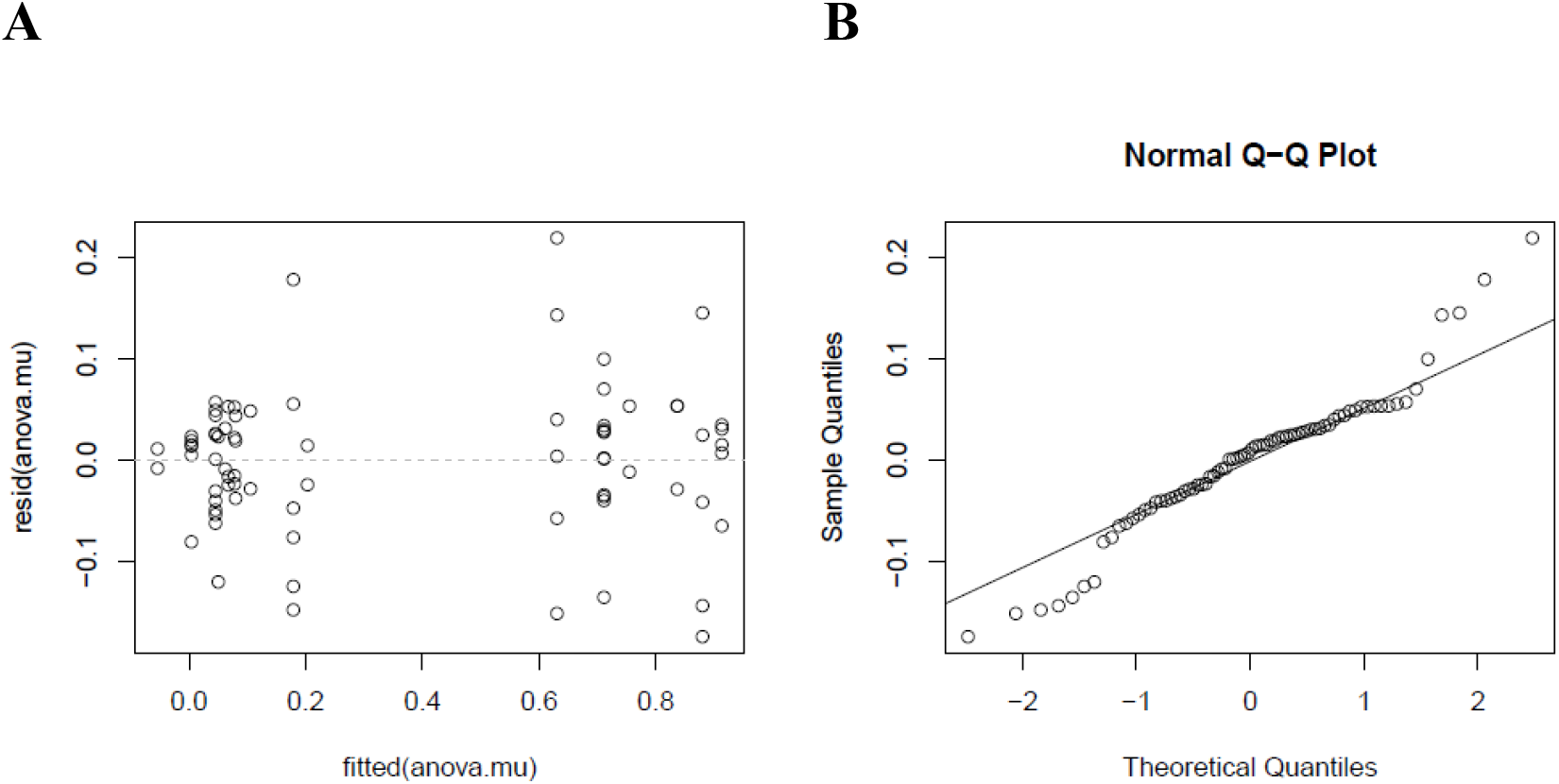
Diagnostic plots for the 1-way ANOVA model used in Fig 3. (A) A residual vs. fitted plot shows that the residual distribution is approximately symmetric regarding 0 for each sample and not strongly affected by the fitted values. (B) Q-Q plot of the residual distribution against the standard normal distribution shows that most residuals are approximately normally distributed.

Table S1. The MS1 and MS/MS data for the samples in Figs 1 to 4, S1, S2, and S4 and Table S2. The columns are: (1) Sample label; (2) Compound (Cmpd), ShA or SA; (3) MS1 or MS/MS (MS2); (4) Isotopologue relative mass number (the one with all-[^12^C] is set to 0); (5) Relative mass number of the ring fragment in MS/MS (NA for MS1); (6) Signal value. The signal values are either the MS1 ratio values vs. isotopologue 0 by a linear regression method for MS1 or the MS/MS signals by an area-under-curve method for MS/MS, for SA and ShA in all the samples.

Table S2. The list of 76 samples with their experimental conditions and the ICS-PAL model parameter estimates. The columns are: (1) Sample label; (2) plant species; (3) UV-C treatment time; (4) [^13^C]-labeling time in hours (h), “all” means all glucose used was [^13^C_6_]glucose, i.e., not isotope-diluted; (5) version of the ICS-PAL model used, “common_ring” or “diff_E4P_ring”; (6) AIC of the model version used; (7) *μ* estimate; (8) standard error of the *μ* estimate; (9) total ShA MS2 signal; (10) total SA MS2 signal; (11) labeled ShA MS2 signal (two or more ring carbons labeled); (12) labeled SA MS2 signal (two or more ring carbons labeled); (13)-(33) the ICS-PAL model parameter estimates, *P*(*L*_*p*_ = 2), *P*(*L*_*p*_ = 1), *P*(*L*_*p*_ = 0), *P*^(*ShA*)^(*L*_*e*_ = 4), *P*^(*ShA*)^(*L*_*e*_ = 3), *P*^(*ShA*)^(*L*_*e*_ = 2), *P*^(*ShA*)^(*L*_*e*_ = 1), *P*^(*ShA*)^(*L*_*e*_ = 0), *P*^(*ShA*)^(*L*_*c*_ = 1|*L*_*p*_ = 2), *P*^(*ShA*)^(*L*_*c*_ = 1|*L*_*p*_ = 1), *P*^(*ShA*)^(*L*_*c*_ = 1|*L*_*p*_ = 0), *τ*^(*ShA*)^, *P*^(*SA*)^(*L*_*e*_ = 4), *P*^(*SA*)^(*L*_*e*_ = 3), *P*^(*SA*)^(*L*_*e*_ = 2), *P*^(*SA*)^(*L*_*e*_ = 1), *P*^(*SA*)^(*L*_*e*_ = 0), *P*^(*SA*)^(*L*_*c*_ = 1|*L*_*p*_ = 2), *P*^(*SA*)^(*L*_*c*_ = 1|*L*_*p*_ = 1), *P*^(*SA*)^(*L*_*c*_ = 1|*L*_*p*_ = 0), *τ*^(*SA*)^. Note that *P*^(*ShA*)^(*L*_*p*_) = *P*^(*SA*)^(*L*_*p*_) in both model versions, and they are shown as *P*(*L*_*p*_). Also, *P*^(*ShA*)^(*L*_*e*_) = *P*^(*SA*)^(*L*_*e*_) in the common-ring model version, so only *P*^(*ShA*)^(*L*_*e*_) is shown.

Table S3. The sources of the plant materials. The columns are: (1) Common plant species name used in the paper; (2) Species binomial nomenclature; (3) additional information about accession, variety, or genotype of the plant; (4) taxonomic family name; (5) taxonomic order name; (6) Source of the plant material; (7) Web site for the internet store or additional information; (8) Stratification conditions when stratification was used before germination of the seed; (9) Growth stage of the plants used for the experiment.

Table S4. The list of 8 samples with their experimental conditions and the PKS-PAL model parameter estimates. The columns are: (1) Sample label; (2) plant species; (3) UV-C treatment time; (4) [^13^C]-labeling time in hours (h), “all” means all glucose used was [^13^C_6_]glucose, i.e., not isotope-diluted; (5) the PKS-PAL model was used; (6)-(12) the PKS-PAL model parameter estimates, *P*(*L*_*a*_ = 1), *P*(*L*_*b*_ = 1|*L*_*a*_ = 1), *P*(*L*_*b*_ = 1|*L*_*a*_ = 0), *P*(*L*_*s*_ = 1), *τ*^(*SA*)^, *P*(*L*_*c*_ = 1|*L*_*p*_), *ϕ*; (13) the standard error of *ϕ*. Note that *P*(*L*_*c*_ = 1|*L*_*p*_) is the same for all *L*_*p*_ values (*L*_*p*_ = 0,1,2) for the PAL pathway.

Text S1. Theoretical basis of the ICS-PAL model and its implementation

Text S2. Theoretical basis of the PKS-PAL model and its implementation

Data S1. The R script for the ICS-PAL model. The script requires Table S1 in the same working directory. It fits the ICS-PAL model to the data for 76 samples (Table S2) and generates Figs 3B, 3C, 3D, S1, and S4 and Table S2. Note that Figs 2 and 4 are excerpts of Fig S1. In addition, it generates “MS1_MS2_regression_v13_c4_sqrt_fin.pdf”, which shows the performance level of the preprocessing of the MS2 data (see Text S1). It also generates two R Data files, “processed.data_v13_c4_sqrt_fin.RData”, which is the preprocessed MS2 data, and “ICS_PAL_model_v13_c5_sqrt_fin_model.fit.RData”, which is the ICS-PAL model fitting results. Both R Data files are required in Data S2.

Data S2. The R script for the PKS-PAL model. The script requires “processed.data_v13_c4_sqrt_fin.RData” and “ICS_PAL_model_v13_c5_sqrt_fin_model.fit.RData” in the same directory. It generates Fig S2 and Table S4 in addition to the R Data file, “PKS_PAL_model_v2_sqrt_fin.RData”, which is the PKS-PAL model fitting results.

Text S1. The theoretical basis of the ICS-PAL model and implementation of the model

## I. Theoretical basis

### I.1 The strategy

The carbons in a single molecule of shikimate (ShA) are derived from one molecule of phosphoenolpyruvate (PEP; we call this PEP molecule PEP #1) and one molecule of erythrose-4-phosphate (E4P). Then the carbons from another PEP molecule (PEP #2) are added to form the 10 carbons of chorismate, which is the last common compound between the isochorismate synthase (ICS) and the phenylalanine ammonia lyase (PAL) pathways. In the context of SA biosynthesis, the PAL pathway starts with chorismate mutase (CM). In chorismate, the carboxyl carbon and the 1-C and 2-C of the benzene ring are derived from 1-C, 2-C, and 3-C of PEP #1, respectively (Fig 1A; blue, yellow, and red circled carbons, respectively).

PEP#1 and PEP#2 are labelled using [^13^C_6_]glucose via glycolysis. (E4P is also labeled from [^13^C_6_]glucose, presumably mainly through Fructose-6-phosphate and a transketolase reaction under the labeling conditions, in which photosynthesis is generally suppressed; see below) Then 1,6-C positions, 2,5-C positions, and 3,4-C positions of glucose result in the 3-C, 2-C and 1-C positions of PEP, respectively (Fig 1A). Note that if glycolysis is the only metabolic process from glucose to PEP, the C-C bond between 3-C and 2-C of glucose and the C-C bond between 4-C and 5-C of glucose are conserved as the C-C bond between 1-C and 2-C of PEP.

If a molecule of salicylate (SA) is synthesized via the ICS pathway, the C-C bond between 1-C and 2-C of PEP#1 is never broken during synthesis of the SA molecule. This C-C bond corresponds to the C-C bond between the carboxyl carbon and the 1-C of the ring in SA (Fig 1A). That is, the carboxyl carbon of SA is conserved from that of ShA.

If a molecule of SA is synthesized via the PAL pathway, the C-C bond between 1-C and 2-C of PEP#1 is broken and the carboxyl carbon of SA is derived from the 3-C of PEP#2 (Fig 1A). That is, the carboxyl carbon of SA is randomized compared to that of ShA.

Thus, the fraction of SA molecules in which the C-C bond between the carboxyl carbon and the 1-C of the ring of SA was conserved from PEP#1 or from Shikimate (ShA) represents the fraction of SA synthesized via the ICS pathway. We denote this fraction as *μ*. Our goal is to estimate the *μ* value. To reach this goal, we statistically modeled the labeling proportions of all 7 carbons in SA and ShA with the following probability variables: *P*^(*SA*)^(*L*_*e*_), *L*_*e*_ = 0,1,2,3,4, the probability of having *L*_*e*_ among the four E4P-originated ring carbons (3- to 6-Cs of the SA ring) [^13^C]-labeled; *P*^(*SA*)^(*L*_*p*_), *L*_*p*_ = 0,1,2, the probability of having *L*_*p*_ of the PEP #1-derived ring carbons (1- and 2-Cs of the SA ring) [^13^C]-labeled; *P*^(*SA*)^(*L*_*c*_|*L*_*p*_), *L*_*c*_ = 0,1, *L*_*p*_ = 0,1,2, the probability of having *L*_*c*_ of the SA carboxyl carbon [^13^C]-labeled, conditional to the *L*_*p*_ value; *P*^(*ShA*)^(*L*_*e*_), *L*_*e*_ = 0,1,2,3,4, the probability of having *L*_*e*_ among the four E4P-originated ring carbons (3- to 6-Cs of the ShA ring) [^13^C]-labeled; *P*^(*ShA*)^(*L*_*p*_), *L*_*p*_ = 0,1,2, the probability of having *L*_*p*_ of the PEP #1-derived ring carbons (1- and 2-Cs of the ShA ring) [^13^C]-labeled; *P*^(*ShA*)^(*L*_*c*_|*L*_*p*_), *L*_*c*_ = 0,1, *L*_*p*_ = 0,1,2, the probability of having *L*_*c*_ of the ShA carboxyl carbon [^13^C]-labeled, conditional to the *L*_*p*_ value. For SA and ShA, 14 probability variables are defined for each compound.

If we can assume that either most labeled SA molecules were synthesized after a steady-state labeling was achieved or that the labeling kinetics from ShA to SA are substantially faster than those from glucose to ShA … (1), we can approximate the labeling states of the rings in SA and ShA as being the same, i.e.,

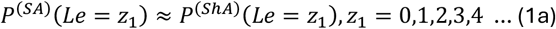

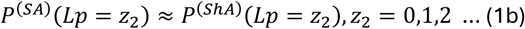

We call assumption (1b) the labeling kinetics assumption of the ICS-PAL model.

With assumption (1),

i. If SA is synthesized through the ICS pathway only, the proportions among the labeling states of the PEP#1-derived carbons are approximately the same between SA and ShA:

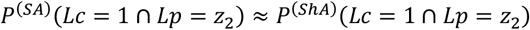

which is the equivalent of:

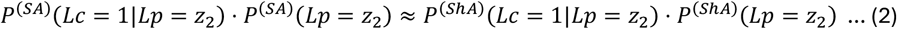
ii. If SA is synthesized through the PAL pathway only, the proportions of the labeled carboxyl carbon conditional to *Lp* are approximately the same for SA across *Lp*:

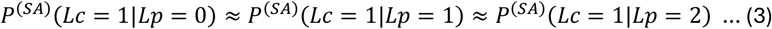

We denote

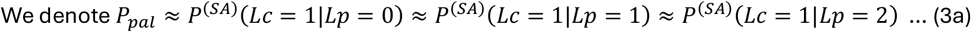

Considering the ICS only case (2), and the PAL only case (3a), the proportion of the labeled SA carboxyl carbon can be expressed as a linear combination between them with the linear coefficient *μ*. Note that ICS is 100% when *μ* = 1, and PAL is 100% when *μ* = 0. When *Lp* = 0,

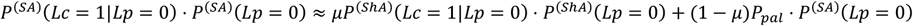

By dividing both sides by *P*^(*SA*)^(*Lp* = 0),

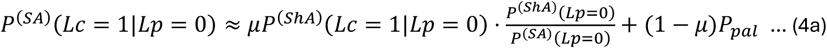

Similarly, when *Lp* = 1,

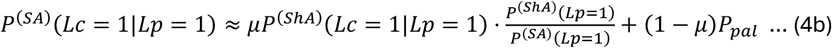

Similarly, when *Lp* = 2,

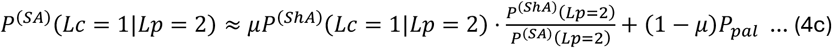

In practice, *P*^(*SA*)^(*Lp* = 1), *P*^(*ShA*)^(*Lp* = 1) are close to 0, and consequently the estimates of *P*^(*SA*)^(*Lc* = 1|*Lp* = 1) · *P*^(*SA*)^(*Lp* = 1), *P*^(*ShA*)^(*Lc* = 1|*Lp* = 1) · *P*^(*ShA*)^(*Lp* = 1) would not have good accuracy. Thus, we ignore (4b).

We eliminate *P*_*pal*_. By subtracting (4a) from (4c),

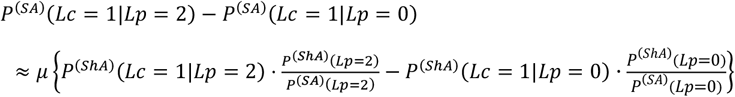

Thus,

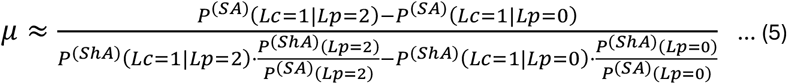

We can obtain the approximate *μ* value if we know eight probability estimates, *P*^(*SA*)^(*Lc* = 1|*Lp* = 2), *P*^(*SA*)^(*Lc* = 1|*Lp* = 0), *P*^(*ShA*)^(*Lc* = 1|*Lp* = 2), *P*^(*ShA*)^(*Lc* = 1|*Lp* = 0),

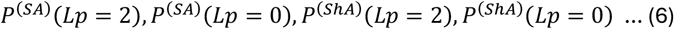

With assumption (1b), (5) can be approximated to:

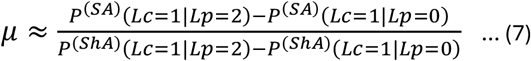

Thus, we can obtain the approximate *μ* value if we know four conditional proportion value estimates,

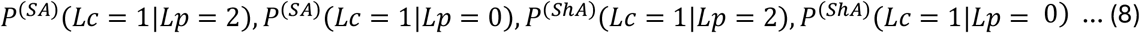

We used assumption (1b) but not assumption (1a), so (1a) is not required for (7) and (8).

### I.2 Statistical model of the labeling state of SA

For this purpose, we include the proportion of SA molecules that pre-existed and remain in the measured sample vs. the total SA molecules in the sample, *τ*^(*SA*)^. Then we have the following 15 (= 14 + 1) variables: *P*^(*SA*)^(*Le* = 0), *P*^(*SA*)^(*Le* = 1), *P*^(*SA*)^(*Le* = 2), *P*^(*SA*)^(*Le* = 3), *P*^(*SA*)^(*Le* = 4), *P*^(*SA*)^(*Lp* = 0), *P*^(*SA*)^(*Lp* = 1), *P*^(*SA*)^(*Lp* = 2), *P*^(*SA*)^(*Lc* = 0|*Lp* = 0), *P*^(*SA*)^(*Lc* = 1|*Lp* = 0), *P*^(*SA*)^(*Lc* = 0|*Lp* = 1), *P*^(*SA*)^(*Lc* = 1|*Lp* = 1), *P*^(*SA*)^(*Lc* = 0|*Lp* = 2), *P*^(*SA*)^(*Lc* = 1|*Lp* = 2), *τ*^(*SA*)^. However, note that the proportion variable values are constrained as follows:

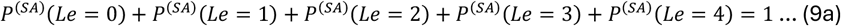

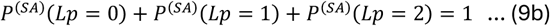

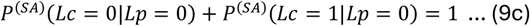

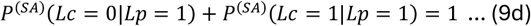

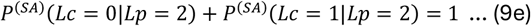

Thus, these represent 10 independent variables.

We denote the proportions of the MS/MS variants of labeled SA (meaning 1 − *τ*^(*SA*)^ of the SA molecules in the sample) as 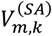, where *m* is the mass variant number of SA and *k* is the mass variant number of the phenol fragment. *m* = 0,1, …,7, *k* = *m, m* − 1,0 ≤ *k* ≤ 6. Then every 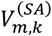 can be expressed using the probability variables:

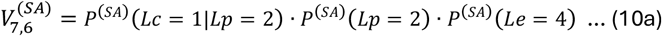

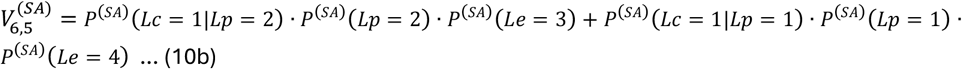

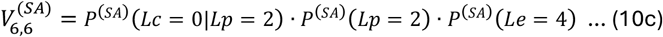

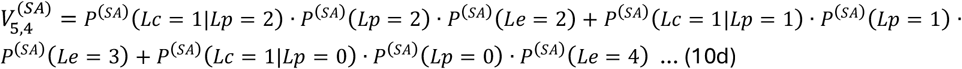

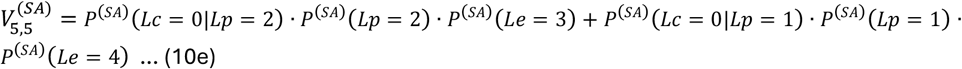

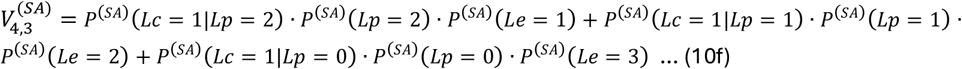

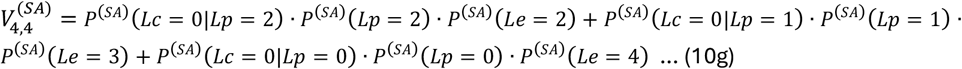

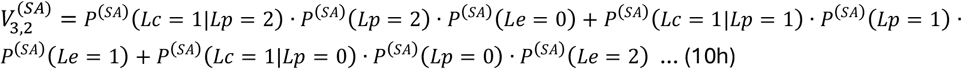

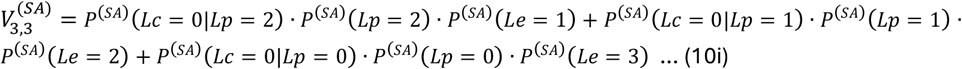

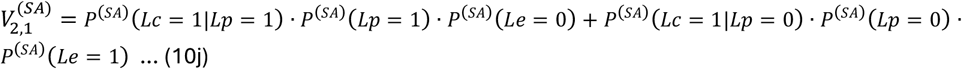

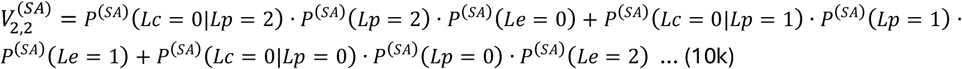

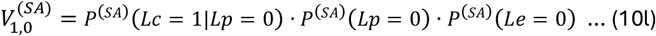

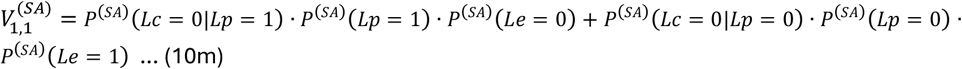

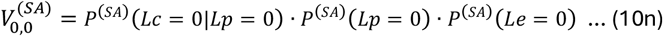

However, in a real assay, a fraction, *τ*^(*SA*)^, of SA detected in the assay was pre-existing before the labeling procedure. Thus, the actual MS/MS measurement 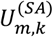 is:

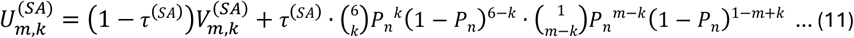

where *P*_*n*_ = 0.0107 is the natural abundance of ^13^C. We use the binomial distribution for the labeling state of the naturally existing SA (the second term in (11)) because we assume that the labeling state of the naturally existing SA is well equilibrated.

By fitting the nonlinear model (10.) and (11) to the actual MS/MS measurements 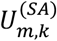, the values for *P*^(*SA*)^(*Lc* = 1|*Lp* = 2) and *P*^(*SA*)^(*Lc* = 1|*Lp* = 0) can be estimated (13 independent observations for 10 independent variables).

### I.3. Statistical model of the labeling state of ShA

By fitting a nonlinear model the same way as (10.) and (11), except for replacing the top-right index “SA” with “ShA”, to the actual MS/MS measurements for ShA, 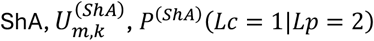 and P^(S*hA*)^(*Lc* = 1|*Lp* = 0) can be estimated (13 independent observations for 10 independent variables). We name the equations corresponding to (10.) and (11) for SA as (12.) and (13) for ShA.

### I.4. An issue with the pre-existing and remaining SA molecules

There is one case that violates the conditions for the *μ* approximation (5). If SA is synthesized through the PAL pathway, it is possible that pre-existing unlabeled ShA molecules can be labeled with labeled PEP#2 and measured as “labeled” SA molecules. Such “labeled” SA molecules are not accounted for as being synthesized from labeled ShA molecules and violate the linear combination assumption (5) … (14). What fraction of the labeled SA molecules are synthesized in this way depends on the labeling kinetics before the incorporation of PEP #2 (including the E4P-originated carbons) and the labeling kinetics after that (the blue “PAL contribution” arrow in Fig 2A and the related text). However, we cannot derive this labeling kinetics information from only the ShA and SA data. While we can be sure that #E2 sample (Fig 2A) had a non-negligible PAL contribution, we cannot precisely estimate how much this violation of the linear combination assumption affects the *μ* estimate.

## II. Practical considerations

### II.1. The numbers of observations and independent variables in the model

First the models for SA and ShA were fit separately, and each model fitting had 13 independent observations for 9 independent variables. The variable number was reduced by one (i.e., 9 instead of 10) by assuming *P*^(*u*)^(L_c_ = 1|*L*_*p*_ = 1) = { P^(*u*)^(L_*c*_ = 1|*L*_*p*_ = 0) + P^(*u*)^(L_*c*_ = 1|*L*_*p*_ = 2)}/2, which should be a tolerable approximation because *P*^(*u*)^(L_c_ = 1|*L*_*p*_ = 0) < *P*^(*u*)^(L_c_ = 1|*L*_*p*_ = 1) < *P*^(*u*)^(L_c_ = 1|*L*_*p*_ = 2) and in most cases *P*^(*u*)^(L_p_ = 1) ≪ P^(*u*)^(L_*p*_ = 0), *P*^(*u*)^(L_p_ = 2). Each model had 4 degrees of freedom.

Then the estimates from these models were used as the starting values for the models fitting ShA and SA together (2 * 13 = 26 independent observations). The common-ring model version used both assumptions (1a) and (1b) and had 12 independent variables. The variable numbers were reduced by two by using fixed values for *P*^(*u*)^(L_c_ = 1|*L*_*p*_ = 1), just like the models for SA and ShA separately above. This resulted in 14 degrees of freedom. The different-E4P-ring model version used only assumption (1b) and had 16 independent variables (4 more variables) with 10 degrees of freedom.

Having reasonably high degrees of freedom in model fitting is important as some of the MS/MS signal values were very low or missing (we filled the missing values with half of the minimum MS/MS signal detected in an experiment). These signal values would be associated with relatively high errors, and we assigned low weights to such values. See “III.1. Preprocessing of the MS and MS/MS data” below.

### II.2. Checking whether assumptions (1a) and (1b) are satisfied in a biological sample

Whether assumptions (1a) and (1b) are satisfied in a particular biological sample can be checked *post hoc* by plotting the *P*^(*SA*)^(*Le* = *z*_1_), P^(SA)^(*Lp* = *z*_2_) values (estimated by fitting all (10.) and (11), i.e., using all MS/MS data for SA) against *P*^(*ShA*)^(*Le* = *z*_1_), P^(Sh*A*)^(*Lp* = *z*_2_) values (estimated by fitting (12.) and (13)) (*z*_1_ = 0,1,2,3,4; *z*_2_ = 0,1,2). If the plotted points are close to *y* = *x*, we can conclude that assumptions (1a) and/or (1b) are satisfied. We call this type of plot the ring carbon labeling parameter plot, e.g., Fig 2C. Fig S1 includes such plots for all samples with good fits of the ICS-PAL model, showing that assumption (1b) is generally satisfied and in many cases, assumption (1a) is also satisfied.

If condition (14) is a substantial issue, the deviation between *P*^(*SA*)^(*Le* = *z*_1_), P^(SA)^(*Lp* = *z*_2_) values and *P*^(*ShA*)^(*Le* = *z*_1_), P^(Sh*A*)^(*Lp* = *z*_2_) values would be mainly driven by the values when *Le* = 0 and/or *Lp* = 0 (since *P* are proportions, they would affect the other values in a predictive manner).

Other types of discrepancies between *P*^(*SA*)^(*Le* = *z*_1_), P^(SA)^(*Lp* = *z*_2_) and P^(*ShA*)^(*Le* = *z*_1_), P^(Sh*A*)^(*Lp* = *z*_2_) could indicate one, two, or all of the following three situations.

i. MS/MS measurements have errors that are too large. This could easily occur when the labeling efficiency of SA and/or ShA is low and/or the amount of SA and/or ShA produced by the plant is low. These can be roughly estimated by the strengths of the MS2 signals for each sample (Fig 3D).
ii. Although all SA molecules are synthesized from the ICS and/or PAL pathways, assumption (1b) is not satisfied, i.e., the labeling kinetics of the steps after ShA are not substantially faster than those of the steps up to ShA. This could be the case when the number of pre-existing and remaining SA molecules is much greater than the number of newly labeled SA molecules, i.e., a sample has a very large *τ*^(*SA*)^ value. This is discussed in the main text related to Figs 2G to 2L.
iii. A large proportion of SA is synthesized by a pathway(s) other than the ICS or PAL pathways (i.e., SA is synthesized mainly via a non-ShA pathway). Basal angiosperm and magnoliids samples were such cases (Fig 5 and the related text). We assumed an SA biosynthetic pathway involving a type III polyketide synthase (PKS). Details of the PKS model are described in Text S2.

### II.3. The labeling state of the E4P-originated ring carbons via the ShA pathway

E4P is most likely synthesized from glucose through glycolysis and the pentose phosphate pathways involving the transketolases (TKs) and transaldolase (TA) under our labeling conditions (Fig TS1.1) [55]. It is likely that E4P was mainly synthesized through glycolysis and the left TK reaction. Either through glycolysis or the pentose phosphate pathway, none of the carbon-carbon bonds in E4P are broken from those in glucose: the bonds of 3-C, 4-C, 5-C, and 6-C in glucose are conserved as the bonds of 1-C, 2-C, 3-C, and 4-C of E4P (cyan, brown, purple, and green circled carbons in Fig TS1.1). The only possible way to disrupt these critical C-C bonds of E4P is that 3-C labeled fructose 6-P #2 is used as a substrate in the left TK reaction. However, the concentration of fructose 6-P #2 would be much lower than that of fructose 6-P #1. Also, for one molecule of fructose 6-P #2, one molecule of all-carbon-labeled E4P is already produced by the TA reaction. Thus, the number of the all-carbon-labeled E4P molecules would be much higher than the number of the single-carbon-labeled E4P made from fructose 6-P #2. Therefore, during labeling with [^13^C_6_]glucose, the labeling states of the majority of E4P are either [^12^C_4_] (no carbon labeled) or [^13^C_4_] (all carbons labeled), which correspond to “e0” and “e4” labeling parameters, respectively, in the ring carbon labeling parameter plot, such as Fig 2C: i.e., if any of the four E4P-originated ring carbons of ShA or SA are labeled, most likely all four carbons are labeled. Thus, among the ring carbon labeling parameters for the E4P-originated carbons, it is very unlikely that any of the labeling parameters “e1”, “e2”, and “e3” have higher values than “e4” if ShA or SA are mainly synthesized through the ShA pathway.

**Fig TS1.1.**
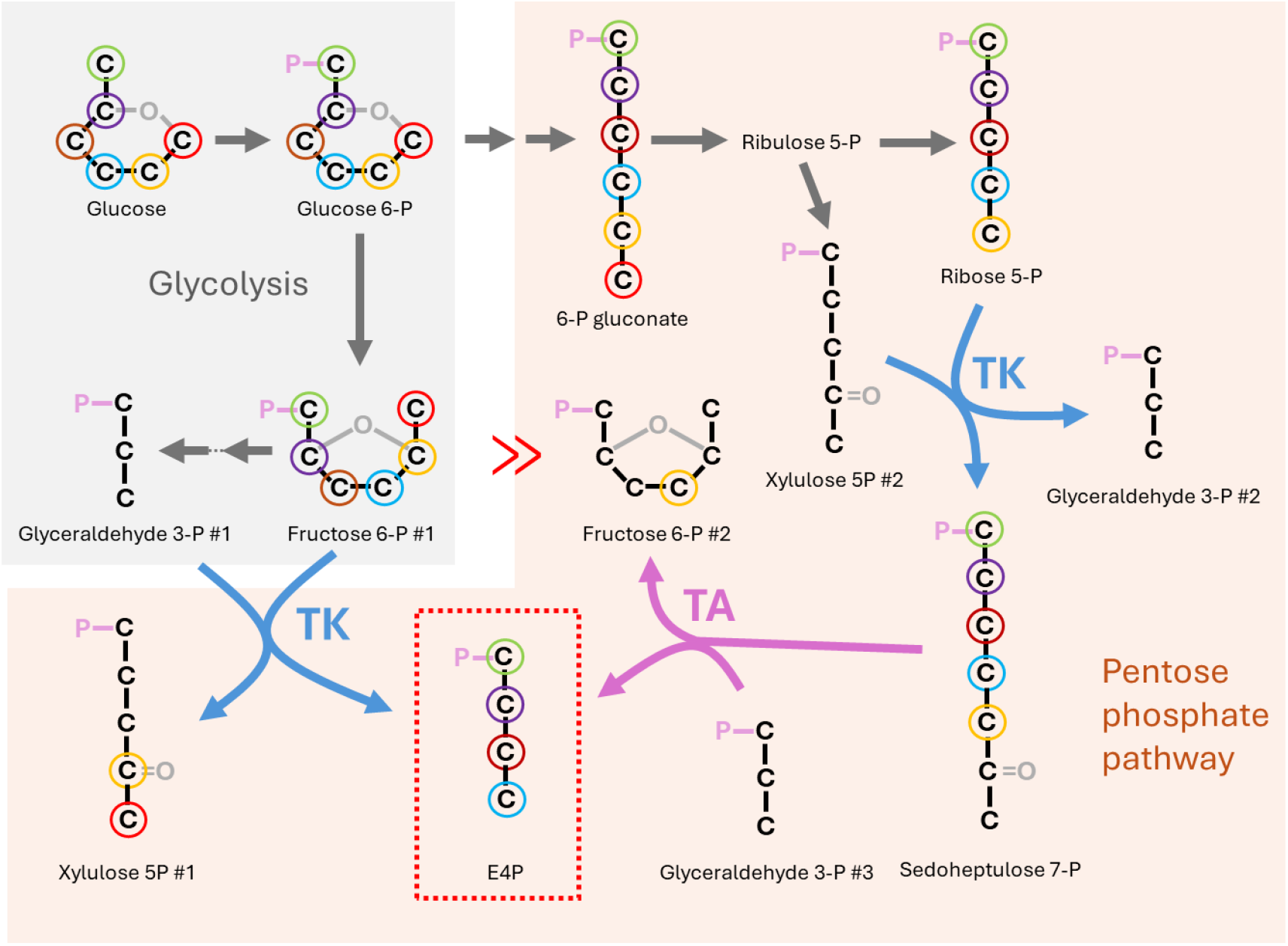
The metabolic processes for synthesis of labeled E4P (red dotted box) from [^13^C_6_]glucose. Glycolysis (gray shaded) and Pentose phosphate pathway (peach shaded) are involved. TK, transketolase and TA, transaldolase. The same metabolites that appear in the figure for different reactions are numbered as #1, #2, …

In fact, in all samples whose data were well-modeled by the ICS-PAL model (i.e., the eudicot and monocot samples), clearly, e4 > e1, e2, e3 in both ShA and SA. On the other hand, many samples from basal angiosperms and magnoliids showed e4 < e1 in SA, strongly suggesting that, under our experimental conditions, the main SA biosynthetic pathway in at least some of these plants is not the ShA pathway.

## III. Implementation

### III.1. Preprocessing of the MS and MS/MS data

We preprocessed the MS1 and MS/MS data for ShA and SA before the measurements were applied to the ICS-PAL or PKS-PAL model fitting. First, we filled missing MS/MS signal values with half of the minimum MS/MS signal value of the experiment. Then according to the signal value of the other MS/MS fragment of the same isotopologue, the weight for the filled in value was assigned: if the other signal value is larger than 5 times the minimum value, weight = 1/9; if the other signal value is equal to or smaller, weight = 1/36; if both MS/MS values of the isotopologue are missing, weight = 1/100 for both. For an MS/MS signal value that was not missing: if the value is equal to or larger than 2.5 times the minimum value, weight = 1; if the value is smaller, the signal value / 2.5 times the minimum value was squared and used as the weight. These weights were used in the ICS-PAL or PKS-PAL model fitting.

Second, the signal values for each isotopologue of ShA or SA were determined as follows. Initially we did not realize that the MS data of the ShA and SA isotopologues often had signals contaminated with signals from some isotopologues of other compounds of the same masses as the LC did not sufficiently separate these compounds from ShA or SA. This situation occurred because many compounds were labeled for their multiple isotoplogues due to the metabolic labeling by [^13^C_6_]glucose, from which labeled carbons flux into many metabolic pathways and because some ShA or SA isotopologue signals were very weak but we need most of them for modeling. Empirically we learned that the MS/MS data usually gave more accurate signal values than the MS1 data most likely because the specific fragmentation patterns of ShA and SA filtered out contaminating signals from other compounds. On the other hand, the MS/MS data for each isotopologue of ShA or SA were collected from an independent LC-MS run, and it may have increased error from small differences in the injection amount, the LC separation, the ionization, and the fragmentation processes. To evaluate the accuracy of the MS1 signal values, we fit a log-log linear regression model with the slope of 1 to the isotopologue signal values calculated from the MS/MS signals (the sum of the MS/MS fragment signals for each isotopologue) vs. the isotopologue signal values from the MS1 measurements. Note that the slope of 1 in the log-log scale means that the response values (the isotopologue signal values calculated from the MS/MS measurements) are proportional to the explanatory variable values (the MS1 isotopologue values). The log-log scale was used because the signal values spread across orders of magnitude. Since we did not measure MS/MS signals for isotopologues 0 and 7, the data used for linear regression were for isotopologues 1 to 6 (6 data pairs). If two or more data pairs out of six were highly off from this linear regression model, we called the data accuracy as too low, and the data were not used. If only one data pair was highly off, the linear regression with the slope of 1 was refit to the other data pairs. For the signal values for isotopologues 1 to 6, the signal values from the MS/MS measurements were used. For the signal values for isotopologues 0 and 7, the values predicted by the linear regression model for the MS1 signal values were used. The plot to visualize the linear regression for each of ShA and SA for each sample in Table S1 can be obtained by executing Data 1: the file name, “MS1_MS2_regression_v13_c4_sqrt_fin.pdf” (Fig TS1.2 for some examples).

In retrospect, it would have been better to measure the MS/MS values for isotopologues 0 and 7 as well (although it would be just one fragment for each) so that the predictions based on the linear regression model would not have been needed.

**Fig TS1.2.**
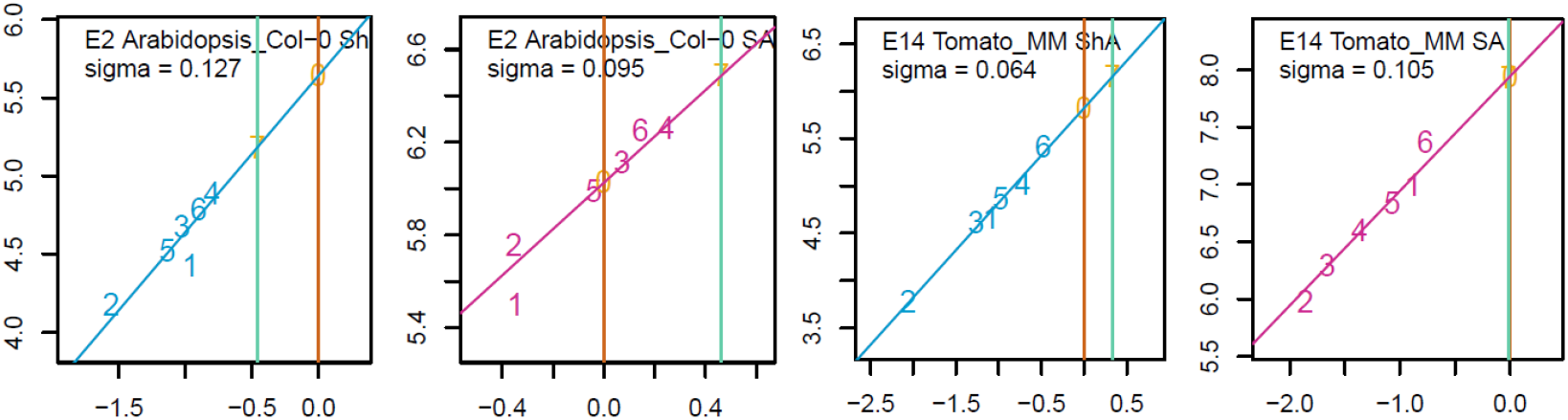
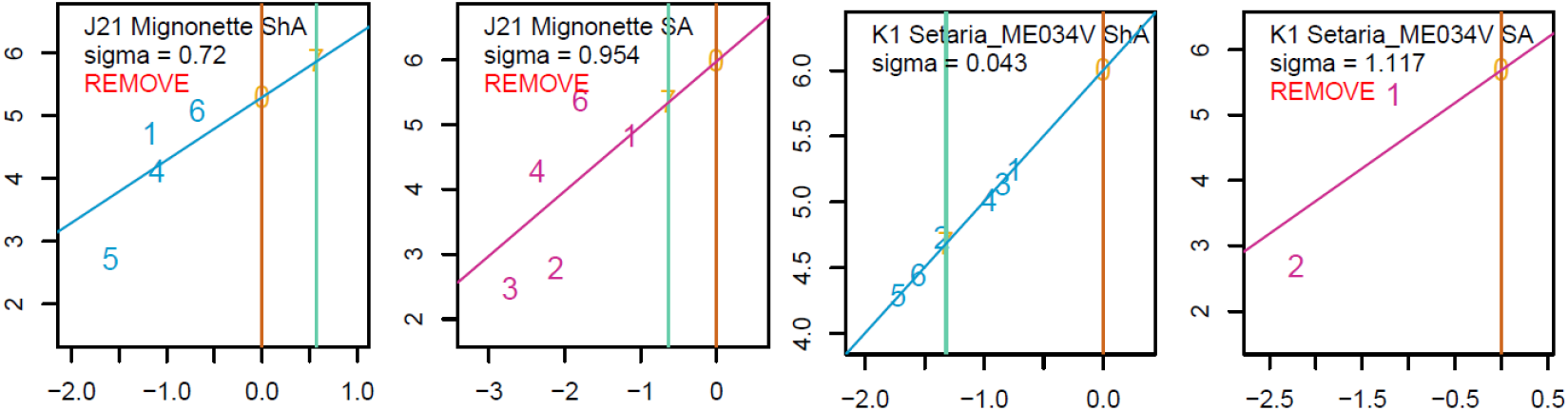
Examples of the MS/MS vs MS1 signal plots. Each plot of ShA and SA (side by side) for four samples, #E2, E13, J21 and K1. Both MS/MS (*y*-axis) and MS1 signal (*x*-axis) values were log-transformed. The number characters 0 to 7 in the plot represent the isotopologues. The values for isotoplopogues 1 to 6 (sky-blue and maroon for ShA and SA, respectively) were used for linear regression with the slope of 1 (assuming that MS/MS and MS1 signals were proportional). The MS signals of isotopologues 0 and 7, where only MS signal values are available, are shown by dark-gold and aquamarine vertical lines, respectively. The red “REMOVE” indicates that the fit of the linear regression in the plot was bad (a high sigma value, which is the residual standard error of the linear regression model), and the corresponding data were not used for the following ICS-PAL model fitting.

Preprocessing of the data was done by the function data.QC (line 138) in Data 1.

### III.2. Key functions in the R script in Data S1

(2.1) Wm_Vmk_v12 (line 14-): a function to calculate the MS/MS values 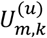, i.e., implementation of (10.) and (11) for *u* = “SA” or (12.) and (13) for *u* = “ShA”, according to m, the numbers of ^13^C out of 7 carbons in the SA or ShA molecule and k, the numbers of ^13^C out of 6 carbons in the corresponding ring fragment. The proportion variables for *P*^(*u*)^(Le), P^(*u*)^(Lp) were reparameterized for constraining the variable value ranges (see section 2.1.1). The function includes the *τ*^(*u*)^ variable as tau.

#### (2.1.1) Reparameterization of the proportion variables

For *P*^(*u*)^(Le = *z*_1_), *z*_1_ = 0,1,2,3,4, to constrain *P*^(*u*)^(Le = *z*_1_) to be

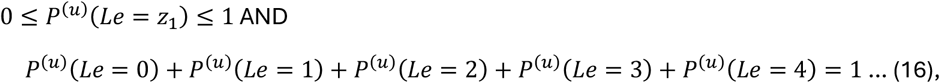

other than one variable value (*P*^(*u*)^(Le = 4) was chosen), the variables were reparameterized as the proportion of the remaining:

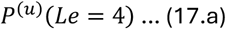

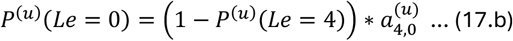

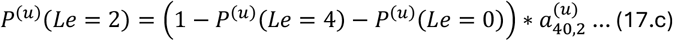

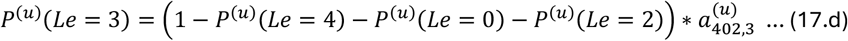

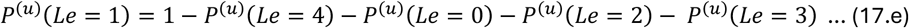

i.e., P^(*u*)^(Le = *z*_1_), *z*_1_ = 0,1,2,3,4 (4 independent variables) were reparameterized with 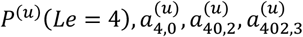 and constraining 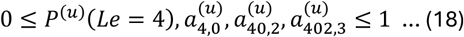 is the equivalent of (16).

The order of reparameterization *P*^(*u*)^(Le), L*e* = 4,0,2,3 was determined based on the initial speculation of their decreasing order. This is because the accuracy of the coarse starting value search (section 2.2) would be better using the decreasing order. The initial speculation was not necessarily correct, but we did not change the order because fundamentally the sets of variables before and after the reparameterization are equivalent.

Similarly, *P*^(*u*)^(Lp = *z*_2_), *z*_2_ = 0,1,2 was reparametrized as:

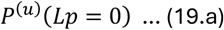

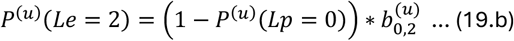

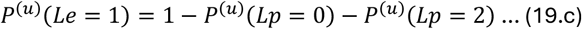

Two reparameterized variables *P*^(*u*)^(Lp = 0), *b*_0,2_ were constrained as

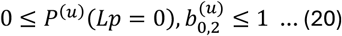

(2.2) mu.s.est.f6 (line 108-): a function for the ICS-PAL statistical model to be fit to the data including both SA and ShA. All proportion variables for SA and ShA are individually specified, i.e., the function does not automatically assume assumptions (1a) and (1b). This function calls the Wm_Vmk_v12 function for each of SA and ShA, which is specified by the Cmpd variable (“SA” or “ShA”).

#### (2.2.1) Reparameterization for the *μ* variable

To fit *μ* directly, in this function, *P*^(*SA*)^(*Lc* = 1|*Lp* = 2) was reparameterized as: from (5),

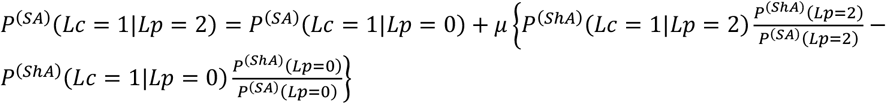

(2.3) para.search (line 301-): a function for a coarse, broad parameter value search for SA or ShA. It calls `Wm_Vmk_v12`. The parameter values were evaluated by the residual of the square-root-transformed data values and model. This search is used to determine the starting values of the parameters in `nls` model fitting. It returns the best parameter value set among those searched and the distributions of the parameter values by quartile among the top 50 parameter value sets.

### III.3. A high level organization of the MAIN part of the R script

For each leaf sample:

(3.1) `Run data.QC`. for the MS1, MS/MS data of ShA and SA separately to yield preprocessed data stored in the list `processed.sample.data`., whose top-level index is the sample label. This list is saved as “processed.data_v13_c4_sqrt_fin.RData”, which is required to execute Data 2 script.

For the ICS-PAL model fitting, the square root-transformed response values and the square root-transformed model were used because the square root-transformation made the residuals of the model distribute properly in general, compared with untransformed and log-transformed. For example, `nls.r2a` in line 707-for the common-ring model version.

(3.2) Fit `nls.r1a` model (line 592-) with 9 independent variables to the data of ShA and SA separately (13 independent observations). The model calls the `Wm_Vmk_v12` function. The start value estimation for the model is performed by the `para.search` function. The results of the model fitting are stored in `list.comps`, including the cases in which the model fitting did not work for ShA and/or SA. If fitting to the ShA or SA data failed, further model fitting for the leaf sample was omitted.

(3.3) Fit the `nls.r2a` model (line 707-) to the ShA and SA data together with both assumptions (1a) and (1b) (the common-ring model version; `comp.name` labeled as `“ShA_SA”`). Then fit the `nls.r2p` model (line 784–) to the ShA and SA data together with assumption (1b) only (the different-E4P-ring model version; comp.name labeled as “ShA_p_SA”). These models call the mu.s.est.f6 function. The estimates from the models in (3.2) were used as the start values for the `nls.r2a` model (for the *P*(*L*_*p*_), *P*(*L*_*e*_) start values common between ShA and SA in the model, the ShA model estimates from (3.2) were used). The estimates from the `nls.r2a` model were used as the start values in the `nls.r2p` model (for the *P*^(*u*)^(L_p_), P^(*u*)^(L_e_) start values specific to ShA and SA in the model, the common *P*(*L*_*p*_), *P*(*L*_*e*_) estimates from the `nls.r2a` model were used).

(3.4) The sample’s *μ* estimate was deemed valid if it satisfied all of the following four conditions: (i) the *μ* standard error is no larger than 0.8; (ii) −0.1 < *μ* < 1.1 ([-0.1, 1.1] was set for the possible *μ* estimate in model fitting; thus, this criterion excludes the boundary values for the fitted *μ* value); (iii) the number of observations with weights higher than 1/80 is at minimum 17, (i.e., there are not too many bad SA observations); (iv) the number of observations with the absolute values of the residuals that are equal to or larger than log4 and with weights equal to or larger than 0.4 is no larger than 1, (i.e., if the residual is large, it should be associated with an observation with a low weight).

Text S2. The theoretical basis of the PKS-PAL model and implementation of the model

### Tracking the [^13^C]-labeling state of the carbons in the PKS pathway only model

We assumed that the carboxyl carbon and the ring carbons of the alkylresorcylate are conserved in SA when SA is synthesized through this hypothetical pathway involving a type III PKS. Note that with the PKS model, we model the labeling state of SA only.

The seven carbons considered originate from the salmon-shaded carbon ([^13^C]-labeled probability, *P*^(*SA*)^(*L*_*s*_)) from the starter acyl-CoA molecule and the olive-shaded carbon ([^13^C]-labeled probability, *P*^(*SA*)^(*L*_*a*_)) and the violet-shaded carbons ([^13^C]-labeled probability, *P*^(*SA*)^(*L*_*b*_)) in three molecules of malonyl-CoA (Fig 1D). The olive- and violet-shaded carbons in a single malonyl-CoA molecule are mostly (100% if glycolysis were the only metabolic pathway from glucose) derived from the 2-C and 1-C of a single PEP molecule and, therefore, have a strong labeling linkage when we use [^13^C6]glucose for labeling. Thus, we instead use the conditional probability *P*^(*SA*)^(*L*_*b*_|*L*_*a*_), thus *P*^(*SA*)^(*L*_*a*_ ∩ L_b_) = P^(*SA*)^(*L*_*b*_|*L*_*a*_)*P*^(*SA*)^(*L*_*a*_) … (1) because the labeling linkage *P*^(*SA*)^(*L*_*b*_|*L*_*a*_) is metabolically constraining and informative regarding the C-C bond between the carboxyl carbon and 1-C of the ring (we always refer to the ring carbon directly connected to the carboxyl carbon as 1-C of the ring to avoid confusion when comparing the models although this carbon position numbering is not conventional as the alkylresorcylate could have a large side chain). For example, *P*^(*SA*)^(*L*_*b*_ = 1|*L*_*a*_ = 1) ≫ *P*^(*SA*)^(*L*_*b*_ = 1|*L*_*a*_ = 0) … (2) is expected.

### The PKS only statistical model

We model the labeling state of SA with *P*^(*SA*)^(*L*_*s*_), *L*_*s*_ = 0,1 (1 independent variable), *P*^(*SA*)^(*L*_*a*_), *L*_*a*_ = 0,1 (1 independent variable), *P*^(*SA*)^(*L*_*b*_|*L*_*a*_), *L*_*a*_ = 0,1, *L*_*b*_ = 0,1 (2 independent variables), and the proportion of the pre-existing and remaining SA molecules in the sample, *τ*^(*SA*)^; total 5 independent variables. Since we do not consider any other pathways, all MS/MS data of SA (13 independent observations) are used in the model fitting.

Since two carbons originate from a single malonyl-CoA molecule at the (5-C and 4-C) or (3-C and 2-C) of the ring are not distinguishable in the MS/MS data of SA, to increase the readability of the equations below, for each of the two carbon positions, we introduce *P*^(*SA*)^(*L*_*m*_) defined as follows:

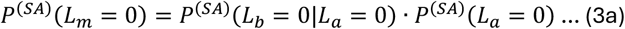

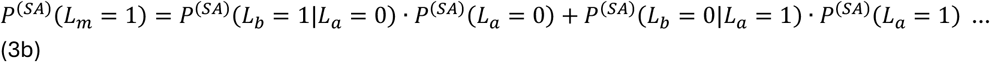

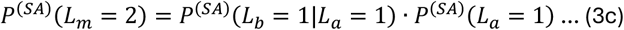

Furthermore, the four ring carbons, 2- to 5-C, are not distinguishable in the MS/MS data of SA. So, the labeling state of these four ring carbons, *P*^(*SA*)^(*L*_25*C*_), L_25*C*_ = 0,1,2,3,4 can be defined as:

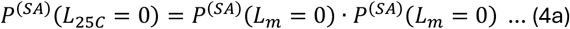

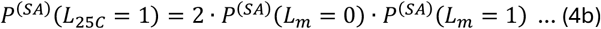

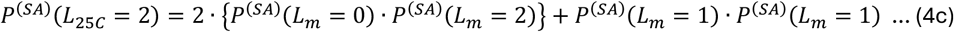

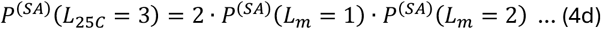

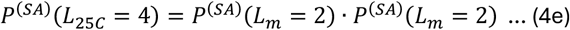

We denote the proportions of the MS/MS variants of labeled SA (meaning 1 − *τ*^(*SA*)^ of the SA molecules in the sample) as 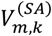, where *m* is the mass variant number of SA and *k* is the mass variant number of the phenol fragment. *m* = 0,1, …,7, *k* = *m, m* − 1,0 ≤ *k* ≤ 6. Then every 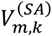 can be expressed using the probability variables:

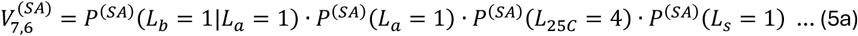

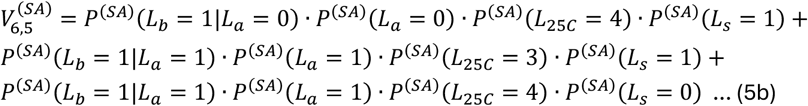

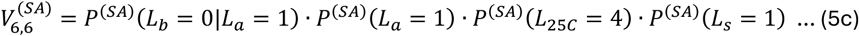

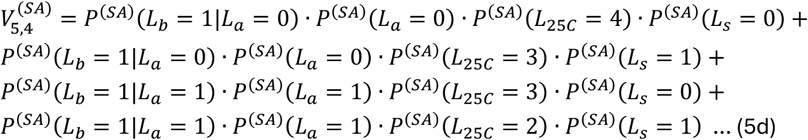

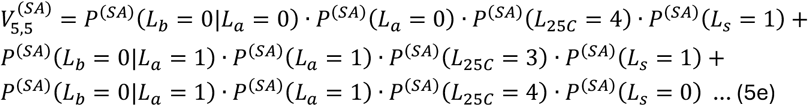

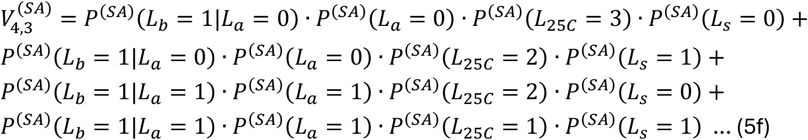

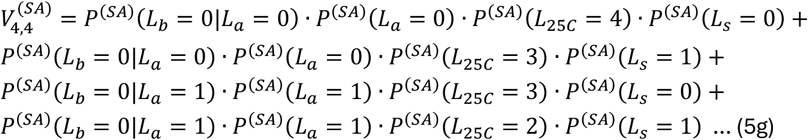

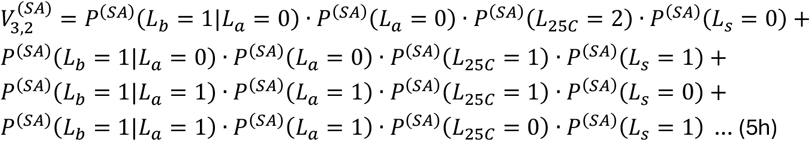

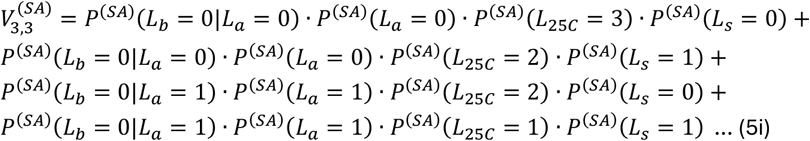

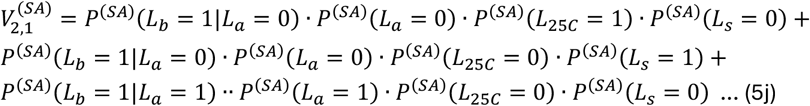

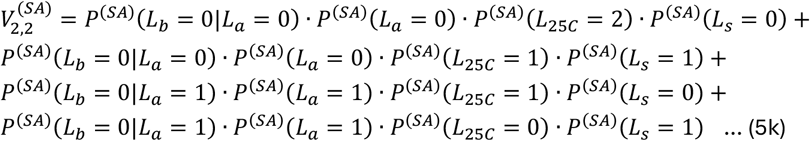

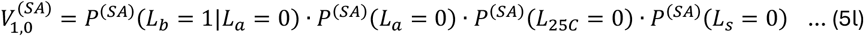

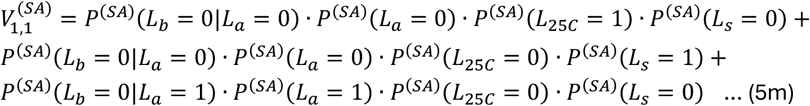

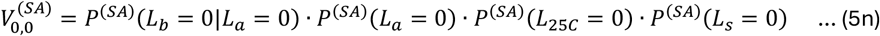

In a real assay, however, a fraction, *τ*^(*SA*)^, of SA detected in the assay was pre-existing before the labeling procedure. Thus, the actual MS/MS measurement 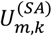 is:

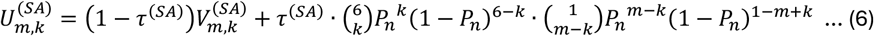

where *P*_*n*_ = 0.0107 is the natural abundance of ^13^C. We use the binomial distribution for the labeling state of the naturally existing SA (the second term in (6)) because we assume that the labeling state of the naturally existing SA is well equilibrated.

By fitting the nonlinear model (3.), (4.), (5.), and (6) to the actual MS/MS measurements 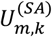, both in square-root-scale, it will become evident whether this PKS model can explain the data well.

### An additional variable whose requirement suggests a minor contribution from the PAL pathway

We realized that the PKS model did not well-explain the proportion of the ring with six [^13^C]-labeled carbons against the other ring isotopolgues, i.e., the data value corresponding to 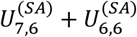, which is the SA data point at *x* = 6 in the ring-carbon labeling profiles in Figs 5 and S2 (“shortcoming #2” in the main text). We initially made the PKS only model to ignore this data point when the model is fit.

Ignoring this data point was achieved by introducing another floating variable, *d*. We denote the data proportion for the ring with six [^13^C]-labeled carbons among all the ring isotopologues, 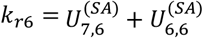, and also denote the modeled values corresponding to 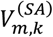 as 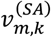. Then we calculated 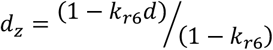. We substituted (5.) with 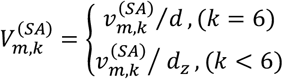. The consequence of this introduction of *d* is that the model can be freely fit to the 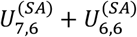 data value by fitting *d*, i.e., ignoring the 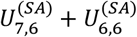 data value. This is the reason the color of the corresponding modeled value points in the ring carbon labeling profiles in Figs 5 and S2 were made into turquoise triangles (the *y*-axis value is the model fitted value for *k*_*r*6_*d*), while that of the other modeled value points were made cyan. Note that the ratio between 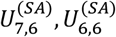 is not affected by variable *d*, and therefore, this ratio seen at *x* = 6 in the carboxyl carbon labeling profiles in Figs 5 and S2 can be directly explained by the PKS only model (5.) and (6).

In addition to shortcoming #2, the PKS only model could not address shortcoming #1, either. We thought that these shortcomings might be addressed by assuming a minor contribution from another SA biosynthesis pathway(s). Since non-Brassicales eudicots and a monocot, rice, are PAL-main, we speculated a minor contribution from the PAL pathway. We developed the PKS-PAL model as follows to test whether a minor contribution from the PAL pathway could address the shortcomings of the PKS only model.

### The PKS-PAL model

To address two shortcomings of the PKS pathway only model, we assumed a minor contribution of the PAL pathway (the PKS-PAL model). Since the number of independent observations with only SA is limited, we decided to use the PAL model parameter values from the model for sample #E14 tomato MoneyMaker (Figs 2D-2F and Table S2) as the base and had only the parameter, *P*^(*SA*)^(*L*_*c*_ = 1|*L*_*p*_ = 0) = *P*^(*SA*)^(*L*_*c*_ = 1|*L*_*p*_ = 1) = *P*^(*SA*)^(*L*_*c*_ = 1|*L*_*p*_ = 2) (these three parameter values are the same under the 100% PAL assumption, i.e., complete randomization of the carboxyl carbon) and the parameter *ϕ* for the proportion of the PKS pathway contribution in the total SA, in addition to the PKS model parameters, float for model fitting. Note that the labeled SA proportion, 1 − *τ*^(*SA*)^, includes SA molecules labeled through both the PKS and PAL pathways and that the parameter *d* value in the PKS model was set to 1 (i.e., removing the *d* parameter).

### Implementation of the PKS-PAL model (the R script in Data S2)

#### 1. Key functions

(1.1) `Wm_Vmk_pks` (line 17-): a function for the PKS only model to calculate the MS/MS values 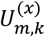, i.e., implementation of (3.), (4.), (5.), and (6) according to m, the number of ^13^C out of 7 carbons in the SA molecule, and `k`, the number of ^13^C out of 6 carbons in the corresponding ring fragment. The function parameters include *P*^(*SA*)^(*L*_*s*_ = 1), *P*^(*SA*)^(*L*_*a*_ = 1), *P*^(*SA*)^(*L*_*b*_ = 1|*L*_*a*_ = 1), *P*^(*SA*)^(*L*_*b*_ = 1|*L*_*a*_ = 0), *τ*^(*SA*)^, *d* as `P_Ls1, P_LMa1, P_LMb1a1, P_LMb1a0, tau_pks, d`. The data 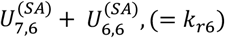 value needs to be given as k1e6.

(1.2) `para.search.pks` (line 212-): a function for a coarse, broad parameter value search for SA with the PKS only model. It calls the `Wm_Vmk_pks` function. The parameter values were evaluated by the residual of the square-root-transformed data values and model. This search was used to determine the starting values of the parameters in `nls` model fitting. It returns the best parameter value set among those searched and the distributions of the parameter values by quartile among the top 1/1e4 parameter value sets.

(1.3) `Wm_Vmk_pks_pal` (line 263-): a function for the PKS-PAL model to calculate the MS/MS values 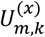. For the PKS part, function (1.1) was used. For the PAL part, the function `Wm_Vmk_v12` (Text S1, section III.2, function (2.1)) was used. In addition to the PKS model parameters, `P_Ls1, P_LMa1, P_LMb1a1, P_LMb1a0, tau_pks`, (but no *d* parameter), the PAL model parameter *P*^(*SA*)^(*L*_*c*_ = 1|*L*_*p*_ = 0) = *P*^(*SA*)^(*L*_*c*_ = 1|*L*_*p*_ = 1) = *P*^(*SA*)^(*L*_*c*_ = 1|*L*_*p*_ = 2) as `Qcajk_fr_1Qcajk_fr_1` and the PKS proportion parameter *ϕ* as phi were included. The PAL parameter values (fixed values) for sample #14 were (Table S2):

`Qem_4 Qem_N40 Qem_N42 Qem_N43 Qak_0 Qak_f2 0.89883395 0.15013187 0.00100000 0.70764314 0.06060478 0.97421657`

(1.4) `para.search.pks.pal` (line 498-): a function for a coarse, broad parameter value search for SA with the PKS-PAL model. It calls the `Wm_Vmk_pks_pal` function. The parameter values were evaluated by the residual of the square-root-transformed data values and model. This search was used to determine the starting values of the parameters in `nls` model fitting. It returns the best parameter value set among those searched and the distributions of the parameter values by quartile among the top 1/1e4 parameter value sets.

#### 2. A high level organization of the MAIN part of the R script

For each leaf sample:

(1.1) The preprocessed SA MS/MS data were used. See Text S1, section III.1 for the preprocessing algorithm used.

(2.2) The PKS-PAL model was fit to the SA MS/MS data in the following order: (i) select the start parameter values using `para.search.pks`; (ii) fit the PKS only model `Wm_Vmk_pks` in `nls.r1b1` (line 853-); (iii) fit the PKS-PAL model `Wm_Vmk_pks_pal` with only `Qcajk_fr_1` and `phi` floating and the PKS model values fixed to the fitted values from (ii) in nls.r1c1 (line 905-); fit all parameters of the PKS-PAL model `Wm_Vmk_pks_pal` with the full parameters using the values from (iii) as the starting values in `nls.r1c2` (line 932-). The fitted values of (iv) are used (Table S2).

