## Supplementary material for "Biosynthesis of a major plant immunity hormone, salicylate, underwent multiple drastic changes during evolution of flowering plants": Fig S2

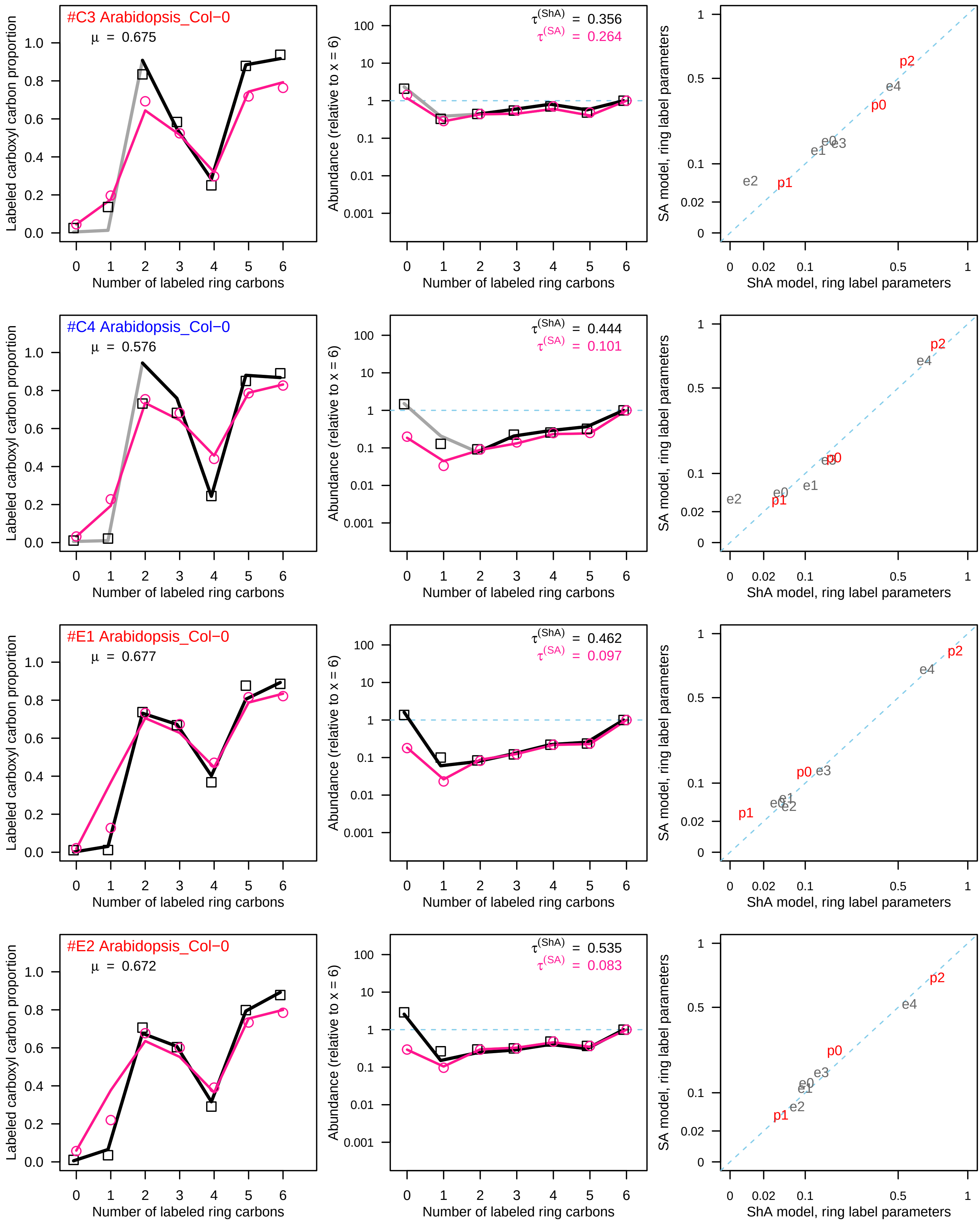
#C3 Arabidopsis_Col−0
μ = 0.675
τ(ShA) = 0.356
τ(SA) = 0.264
#C4 Arabidopsis_Col−0
μ = 0.576
τ(ShA) = 0.444
τ(SA) = 0.101
#E1 Arabidopsis_Col−0
μ = 0.677
τ(ShA) = 0.462
τ(SA) = 0.097
#E2 Arabidopsis_Col−0
μ = 0.672
τ(ShA) = 0.535
τ(SA) = 0.083
Labeled carboxyl carbon proportion
Abundance (relative to x = 6)
SA model, ring label parameters
Number of labeled ring carbons
ShA model, ring label parameters
e0
e1
e2
e3
e4
p0
p1
p2

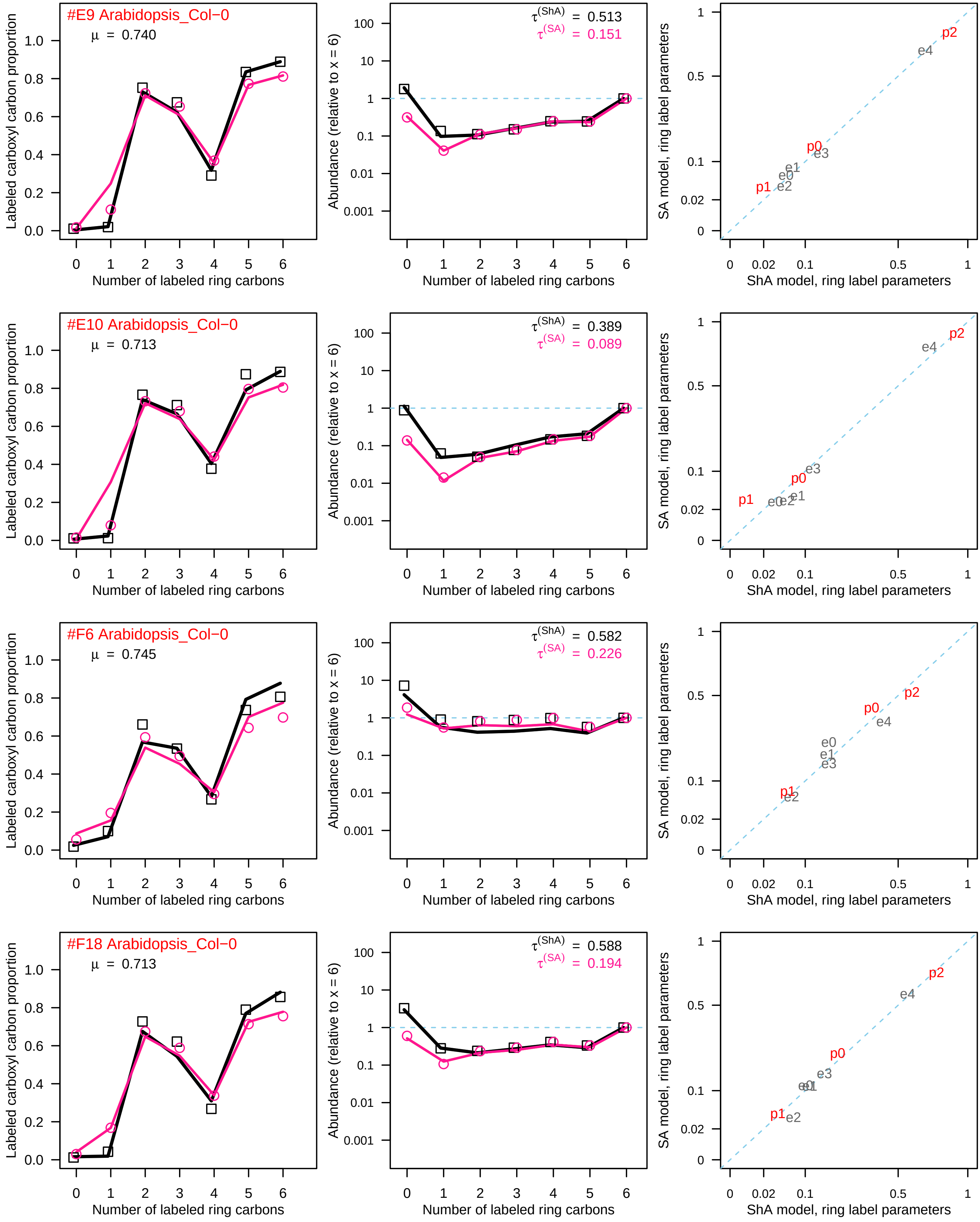

#E9 Arabidopsis_Col−0
μ = 0.740
τ(ShA) = 0.513
τ(SA) = 0.151
#E10 Arabidopsis_Col−0
μ = 0.713
τ(ShA) = 0.389
τ(SA) = 0.089
#F6 Arabidopsis_Col−0
μ = 0.745
τ(ShA) = 0.582
τ(SA) = 0.226
#F18 Arabidopsis_Col−0
μ = 0.713
τ(ShA) = 0.588
τ(SA) = 0.194
Labeled carboxyl carbon proportion
Abundance (relative to x = 6)
SA model, ring label parameters
Number of labeled ring carbons
ShA model, ring label parameters
p0
p1
p2
e0
e1
e2
e3
e4

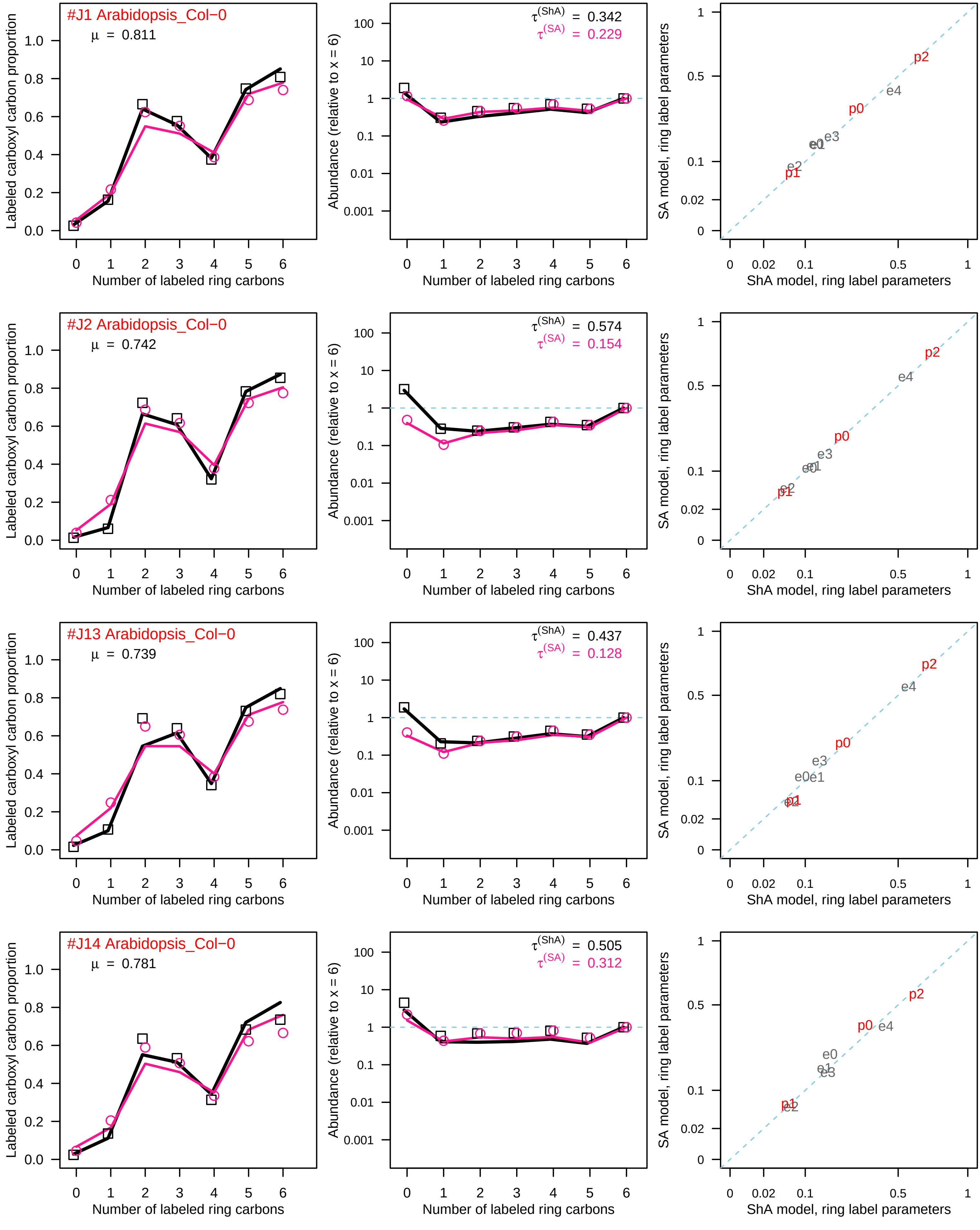
#J1 Arabidopsis_Col−0
μ = 0.811
τ(ShA) = 0.342
τ(SA) = 0.229
#J2 Arabidopsis_Col−0
μ = 0.742
τ(ShA) = 0.574
τ(SA) = 0.154
#J13 Arabidopsis_Col−0
μ = 0.739
τ(ShA) = 0.437
τ(SA) = 0.128
#J14 Arabidopsis_Col−0
μ = 0.781
τ(ShA) = 0.505
τ(SA) = 0.312
Labeled carboxyl carbon proportion
Abundance (relative to x = 6)
Number of labeled ring carbons
SA model, ring label parameters
ShA model, ring label parameters
p0
p1
p2
e0
e1
e2
e3
e4

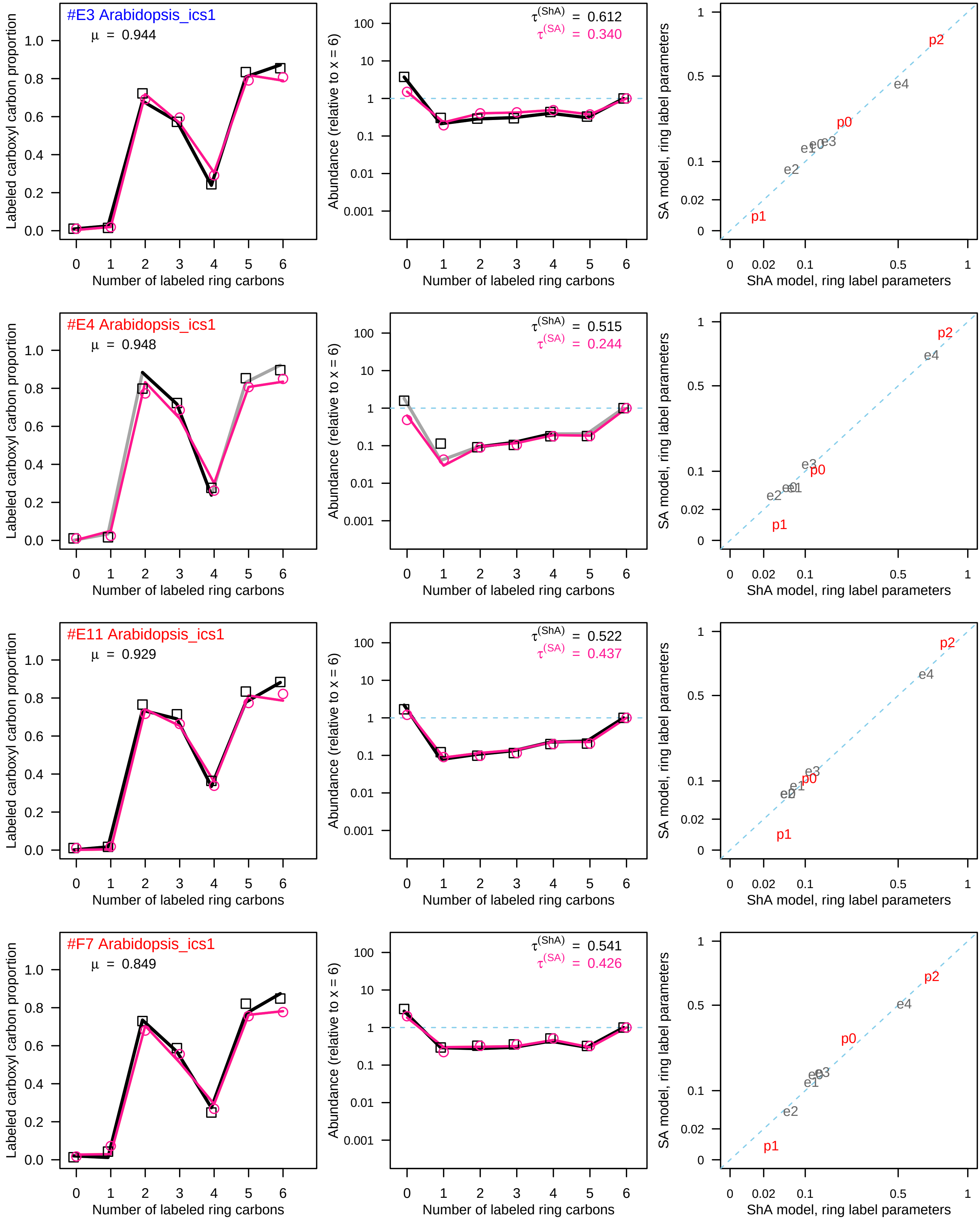
#E3 Arabidopsis_ics1
μ = 0.944
τ(ShA) = 0.612
τ(SA) = 0.340
#E4 Arabidopsis_ics1
μ = 0.948
τ(ShA) = 0.515
τ(SA) = 0.244
#E11 Arabidopsis_ics1
μ = 0.929
τ(ShA) = 0.522
τ(SA) = 0.437
#F7 Arabidopsis_ics1
μ = 0.849
τ(ShA) = 0.541
τ(SA) = 0.426
Labeled carboxyl carbon proportion
Abundance (relative to x = 6)
SA model, ring label parameters
Number of labeled ring carbons
ShA model, ring label parameters
p0
p1
p2
e0
e1
e2
e3
e4

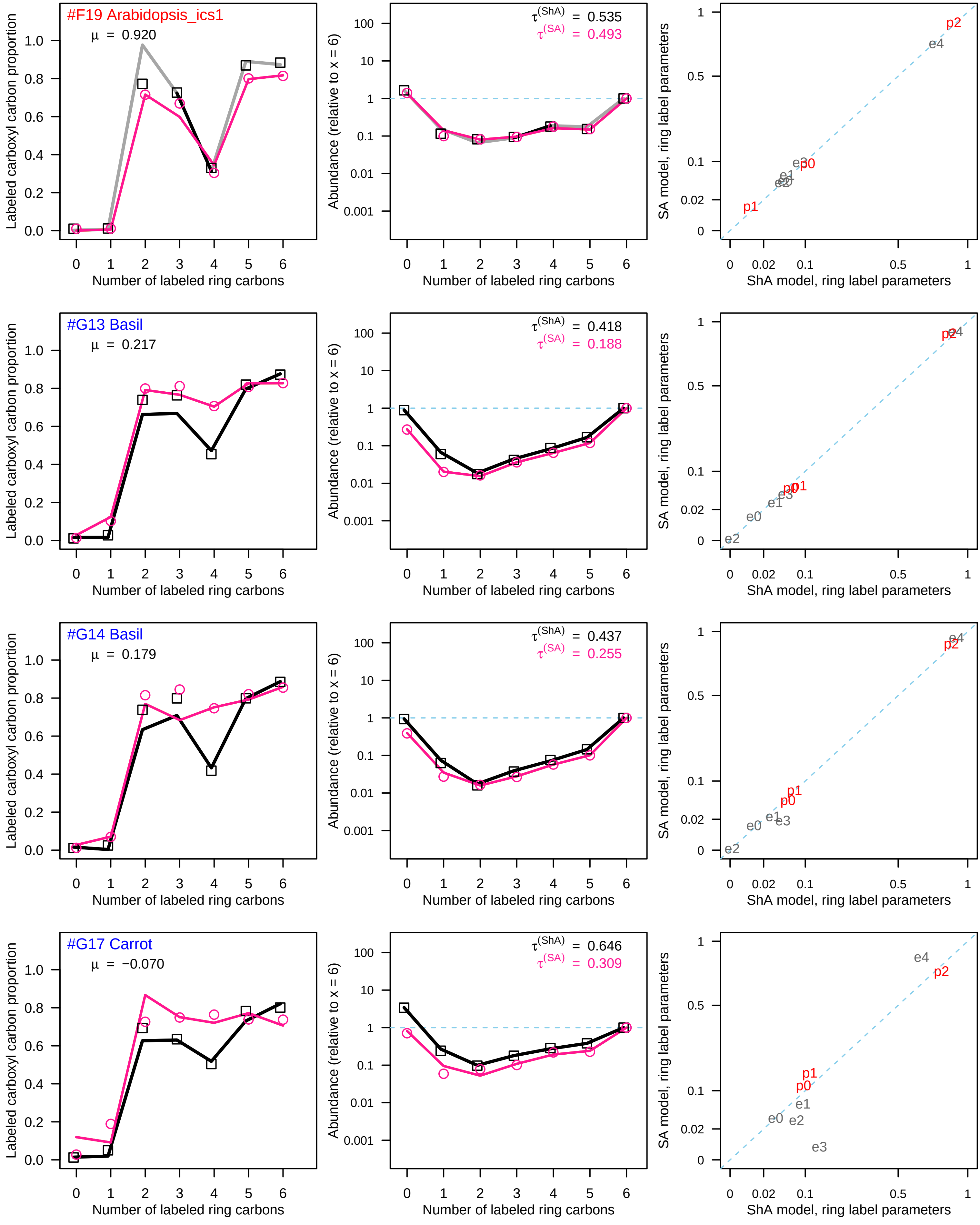
#F19 Arabidopsis_ics1
μ = 0.920
τ(ShA) = 0.535
τ(SA) = 0.493
#G13 Basil
μ = 0.217
τ(ShA) = 0.418
τ(SA) = 0.188
#G14 Basil
μ = 0.179
τ(ShA) = 0.437
τ(SA) = 0.255
#G17 Carrot
μ = −0.070
τ(ShA) = 0.646
τ(SA) = 0.309
Labeled carboxyl carbon proportion
Abundance (relative to x = 6)
Number of labeled ring carbons
SA model, ring label parameters
ShA model, ring label parameters
p0
p1
p2
e0
e1
e2
e3
e4

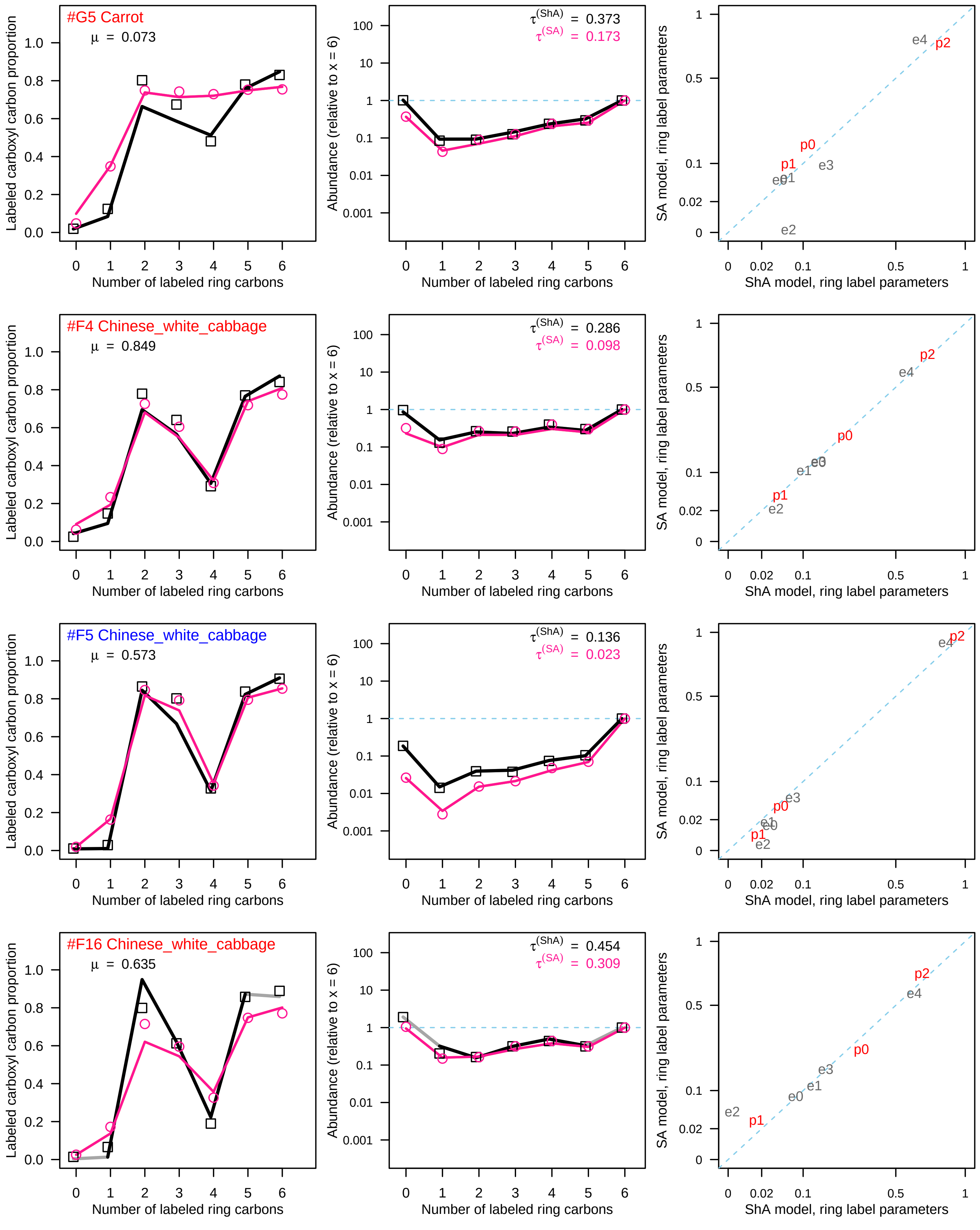
#G5 Carrot
μ = 0.073
τ(ShA) = 0.373
τ(SA) = 0.173
#F4 Chinese_white_cabbage
μ = 0.849
τ(ShA) = 0.286
τ(SA) = 0.098
#F5 Chinese_white_cabbage
μ = 0.573
τ(ShA) = 0.136
τ(SA) = 0.023
#F16 Chinese_white_cabbage
μ = 0.635
τ(ShA) = 0.454
τ(SA) = 0.309
Labeled carboxyl carbon proportion
Abundance (relative to x = 6)
SA model, ring label parameters
Number of labeled ring carbons
ShA model, ring label parameters
p0
p1
p2
e0
e1
e2
e3
e4

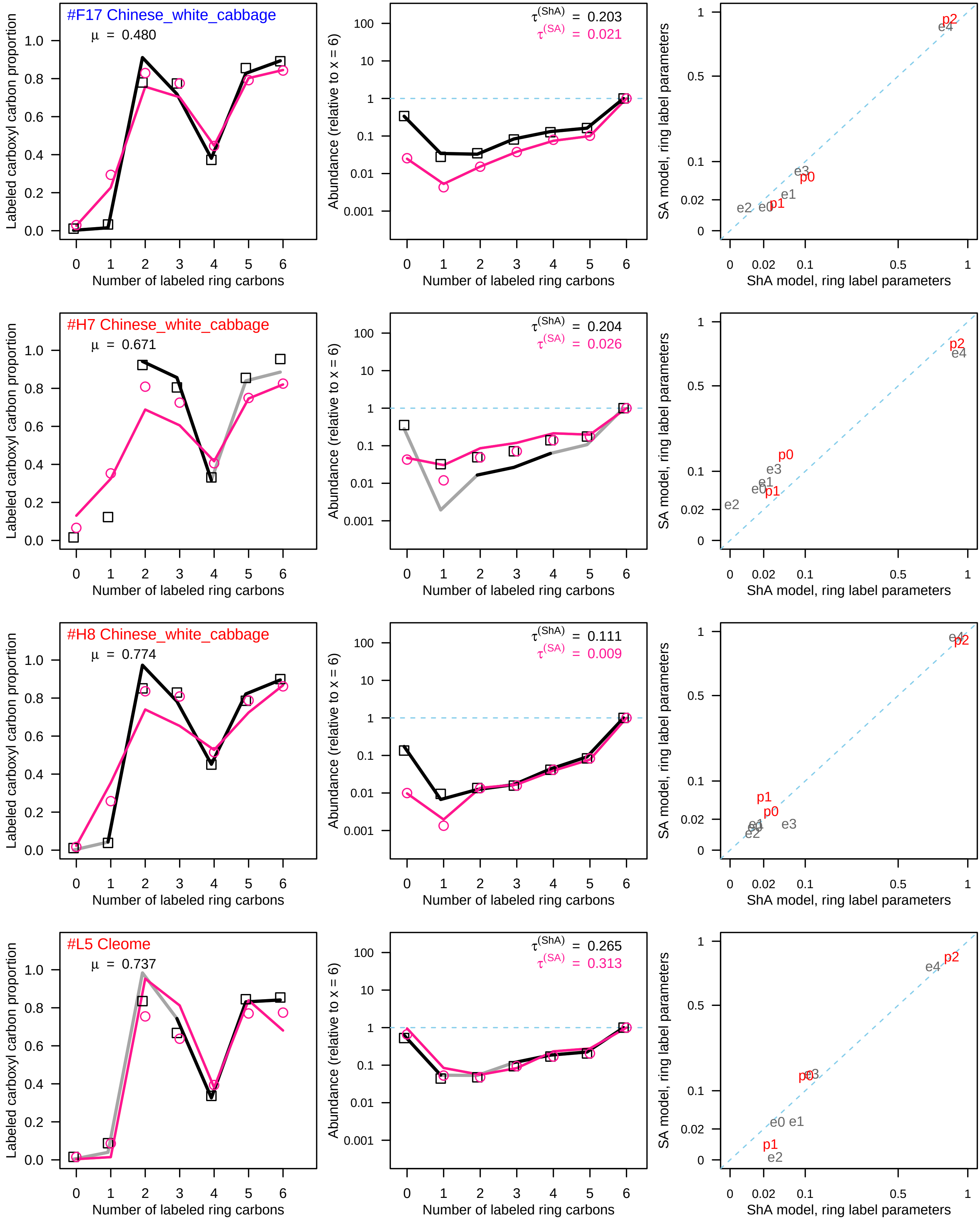
#F17 Chinese_white_cabbage
μ = 0.480
τ(ShA) = 0.203
τ(SA) = 0.021
#H7 Chinese_white_cabbage
μ = 0.671
τ(ShA) = 0.204
τ(SA) = 0.026
#H8 Chinese_white_cabbage
μ = 0.774
τ(ShA) = 0.111
τ(SA) = 0.009
#L5 Cleome
μ = 0.737
τ(ShA) = 0.265
τ(SA) = 0.313
Labeled carboxyl carbon proportion
Abundance (relative to x = 6)
SA model, ring label parameters
Number of labeled ring carbons
ShA model, ring label parameters
p0 p1 p2 e0 e1 e2 e3 e4

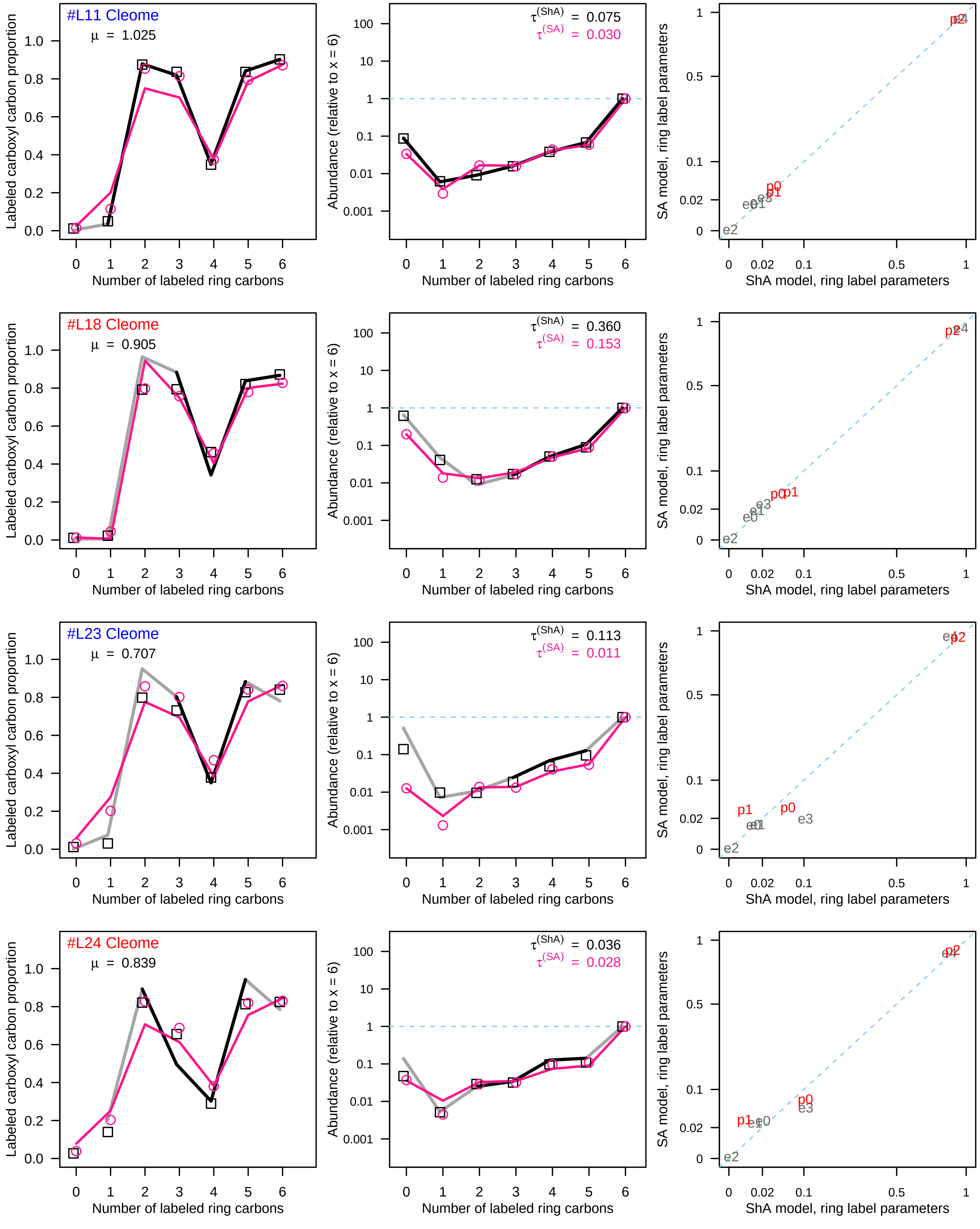

#L11 Cleome
μ = 1.025
τ(ShA) = 0.075
τ(SA) = 0.030
#L18 Cleome
μ = 0.905
τ(ShA) = 0.360
τ(SA) = 0.153
#L23 Cleome
μ = 0.707
τ(ShA) = 0.113
τ(SA) = 0.011
#L24 Cleome
μ = 0.839
τ(ShA) = 0.036
τ(SA) = 0.028
Labeled carboxyl carbon proportion
Abundance (relative to x = 6)
SA model, ring label parameters
Number of labeled ring carbons
ShA model, ring label parameters

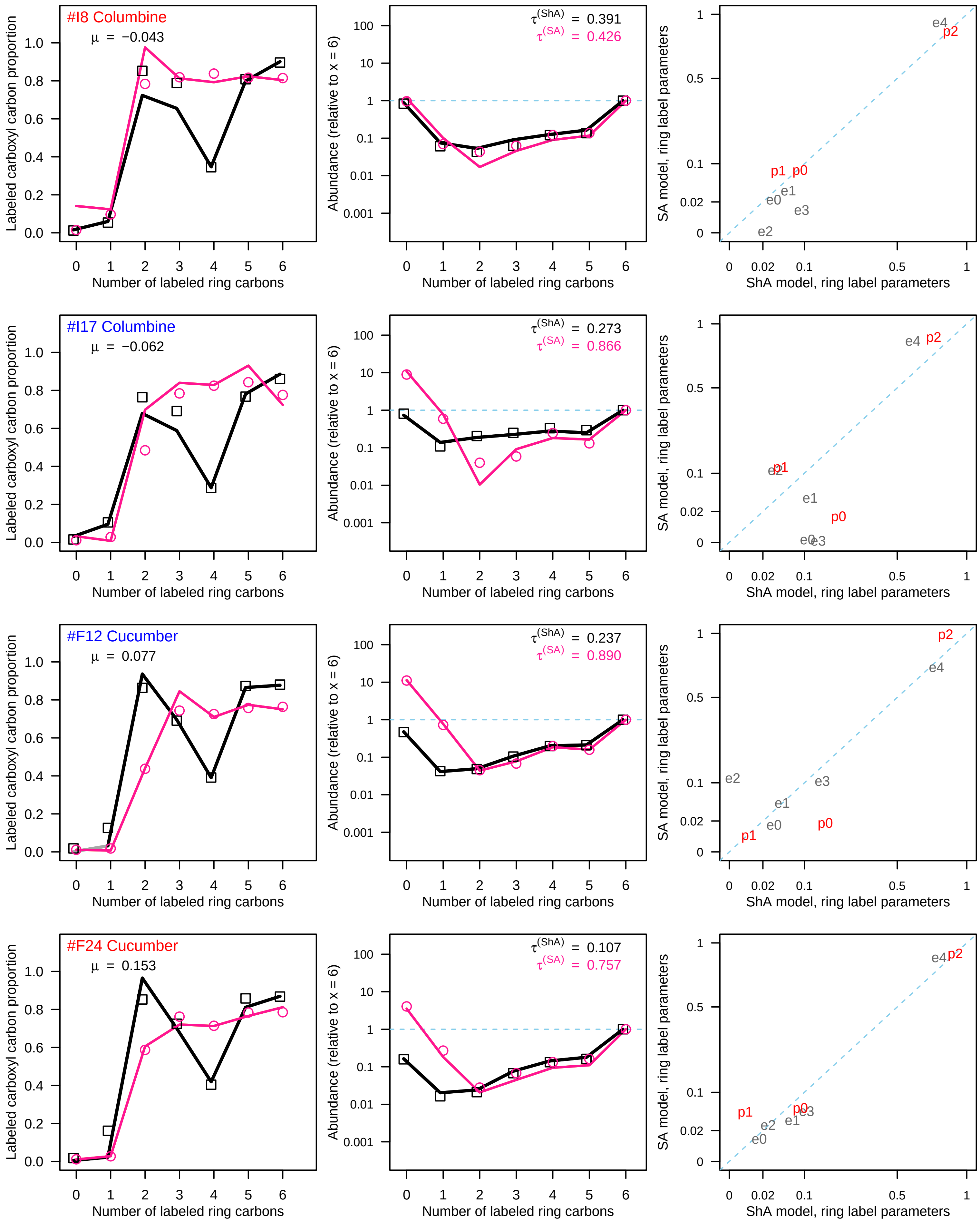
#I8 Columbine
μ = −0.043
τ(ShA) = 0.391
τ(SA) = 0.426
#I17 Columbine
μ = −0.062
τ(ShA) = 0.273
τ(SA) = 0.866
#F12 Cucumber
μ = 0.077
τ(ShA) = 0.237
τ(SA) = 0.890
#F24 Cucumber
μ = 0.153
τ(ShA) = 0.107
τ(SA) = 0.757
Labeled carboxyl carbon proportion
Number of labeled ring carbons
Abundance (relative to x = 6)
SA model, ring label parameters
ShA model, ring label parameters

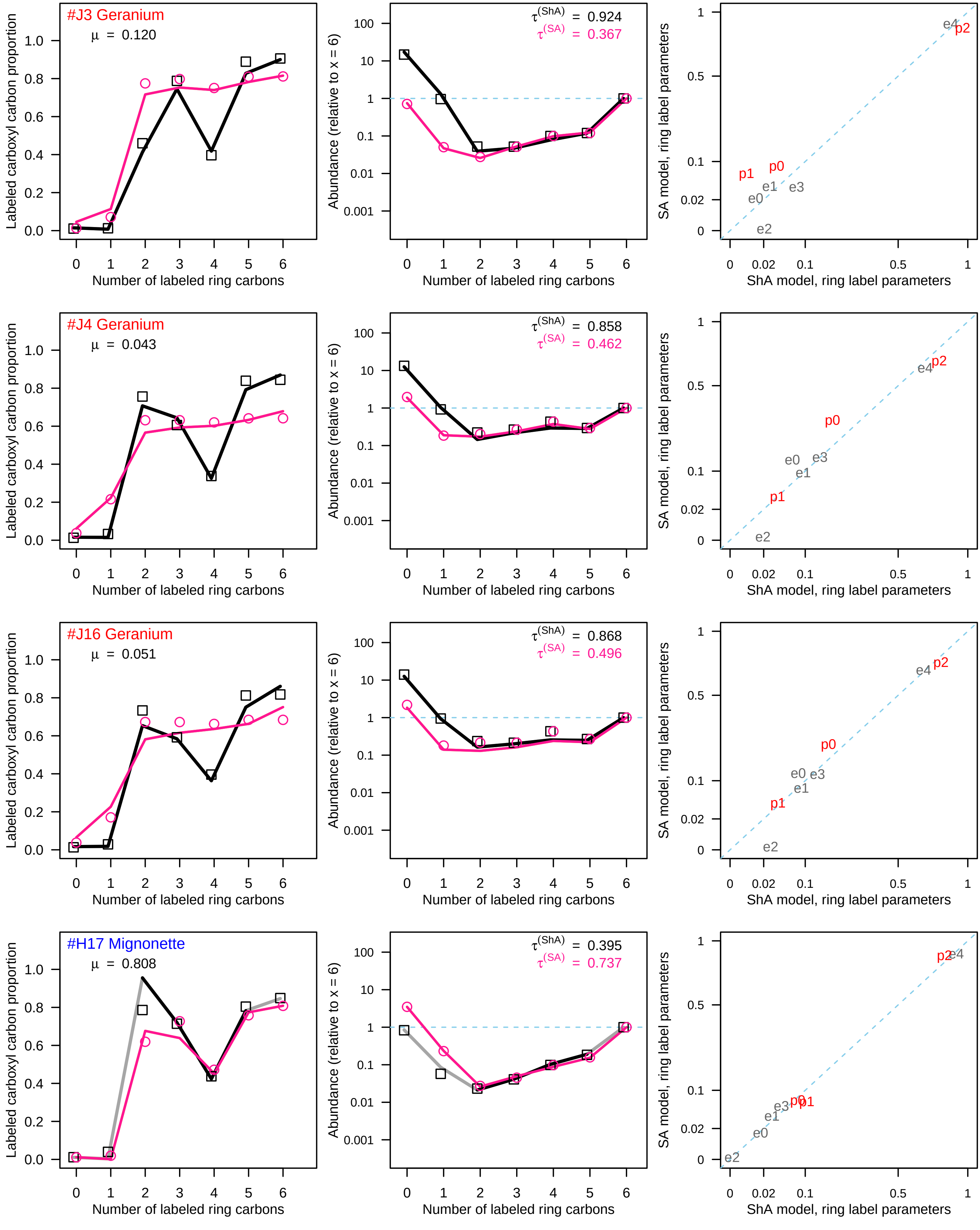

#J3 Geranium
μ = 0.120
τ(ShA) = 0.924
τ(SA) = 0.367
#J4 Geranium
μ = 0.043
τ(ShA) = 0.858
τ(SA) = 0.462
#J16 Geranium
μ = 0.051
τ(ShA) = 0.868
τ(SA) = 0.496
#H17 Mignonette
μ = 0.808
τ(ShA) = 0.395
τ(SA) = 0.737
Labeled carboxyl carbon proportion
Abundance (relative to x = 6)
SA model, ring label parameters
Number of labeled ring carbons
ShA model, ring label parameters
p0
p1
p2
e0
e1
e2
e3
e4

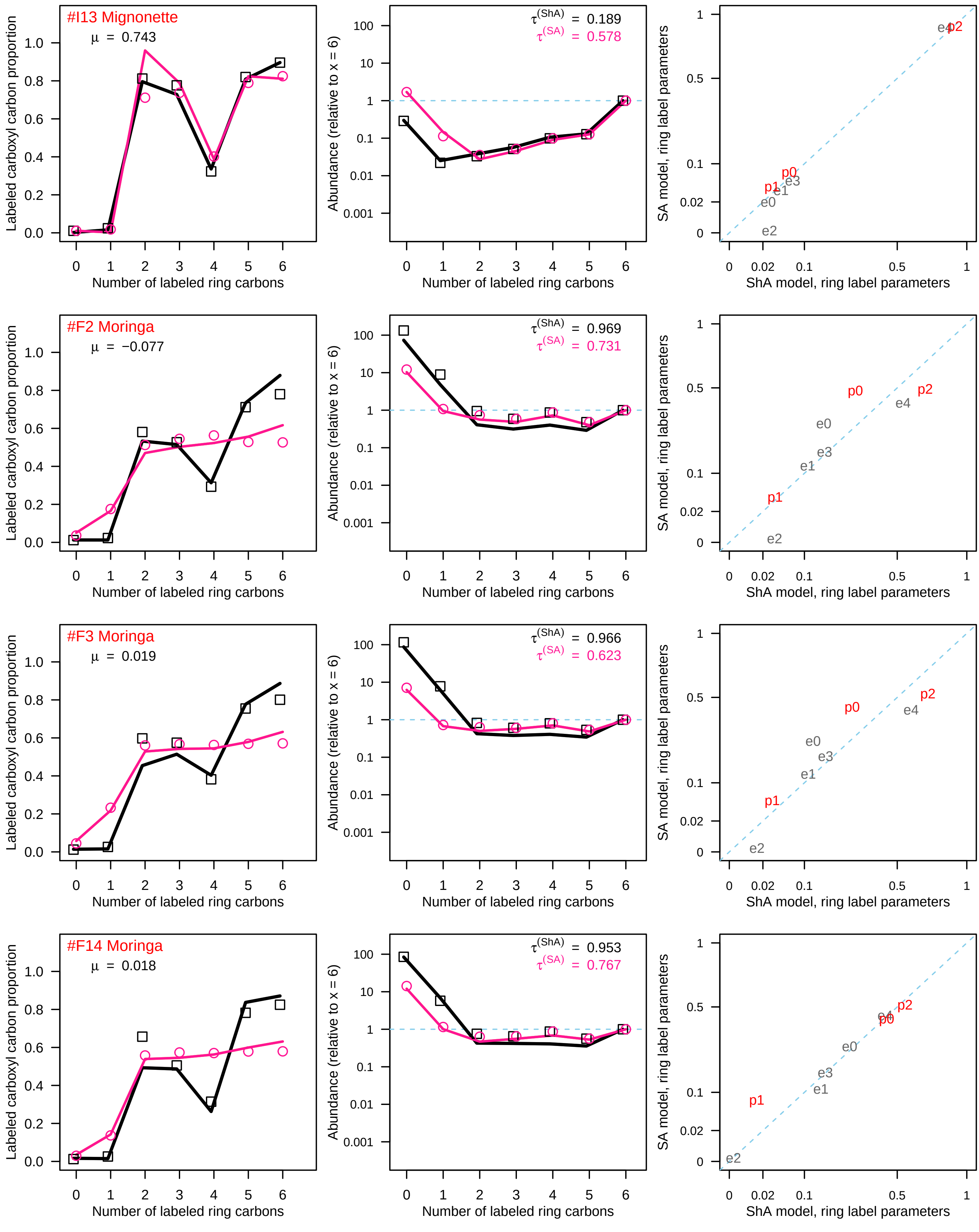
#I13 Mignonette
μ = 0.743
τ(ShA) = 0.189
τ(SA) = 0.578
#F2 Moringa
μ = −0.077
τ(ShA) = 0.969
τ(SA) = 0.731
#F3 Moringa
μ = 0.019
τ(ShA) = 0.966
τ(SA) = 0.623
#F14 Moringa
μ = 0.018
τ(ShA) = 0.953
τ(SA) = 0.767
Labeled carboxyl carbon proportion
Abundance (relative to x = 6)
SA model, ring label parameters
Number of labeled ring carbons
ShA model, ring label parameters
p0
p1
p2
e0
e1
e2
e3
e4

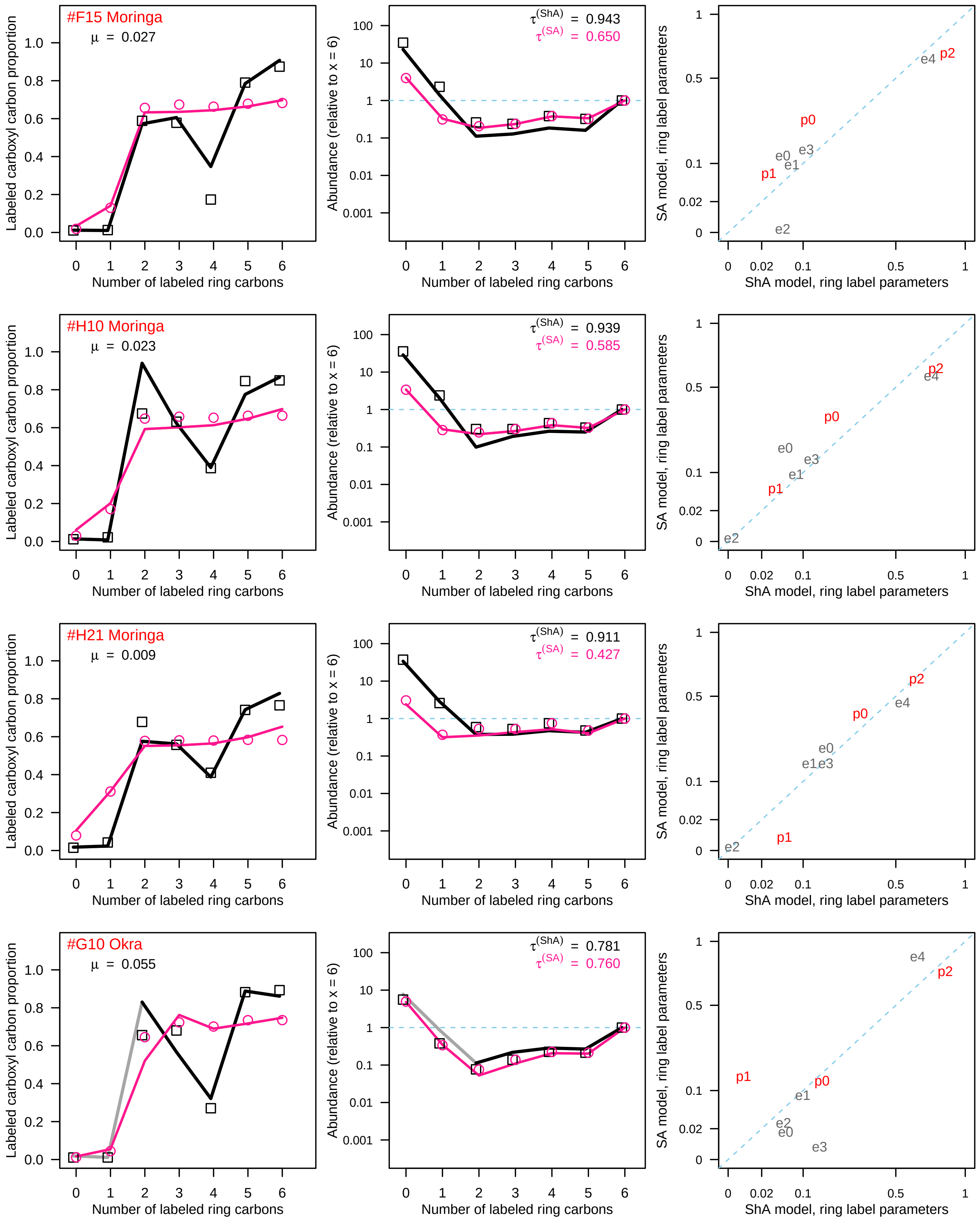
#F15 Moringa
μ = 0.027
τ(ShA) = 0.943
τ(SA) = 0.650
#H10 Moringa
μ = 0.023
τ(ShA) = 0.939
τ(SA) = 0.585
#H21 Moringa
μ = 0.009
τ(ShA) = 0.911
τ(SA) = 0.427
#G10 Okra
μ = 0.055
τ(ShA) = 0.781
τ(SA) = 0.760
Labeled carboxyl carbon proportion
Abundance (relative to x = 6)
SA model, ring label parameters
Number of labeled ring carbons
ShA model, ring label parameters
p0
p1
p2
e0
e1
e2
e3
e4

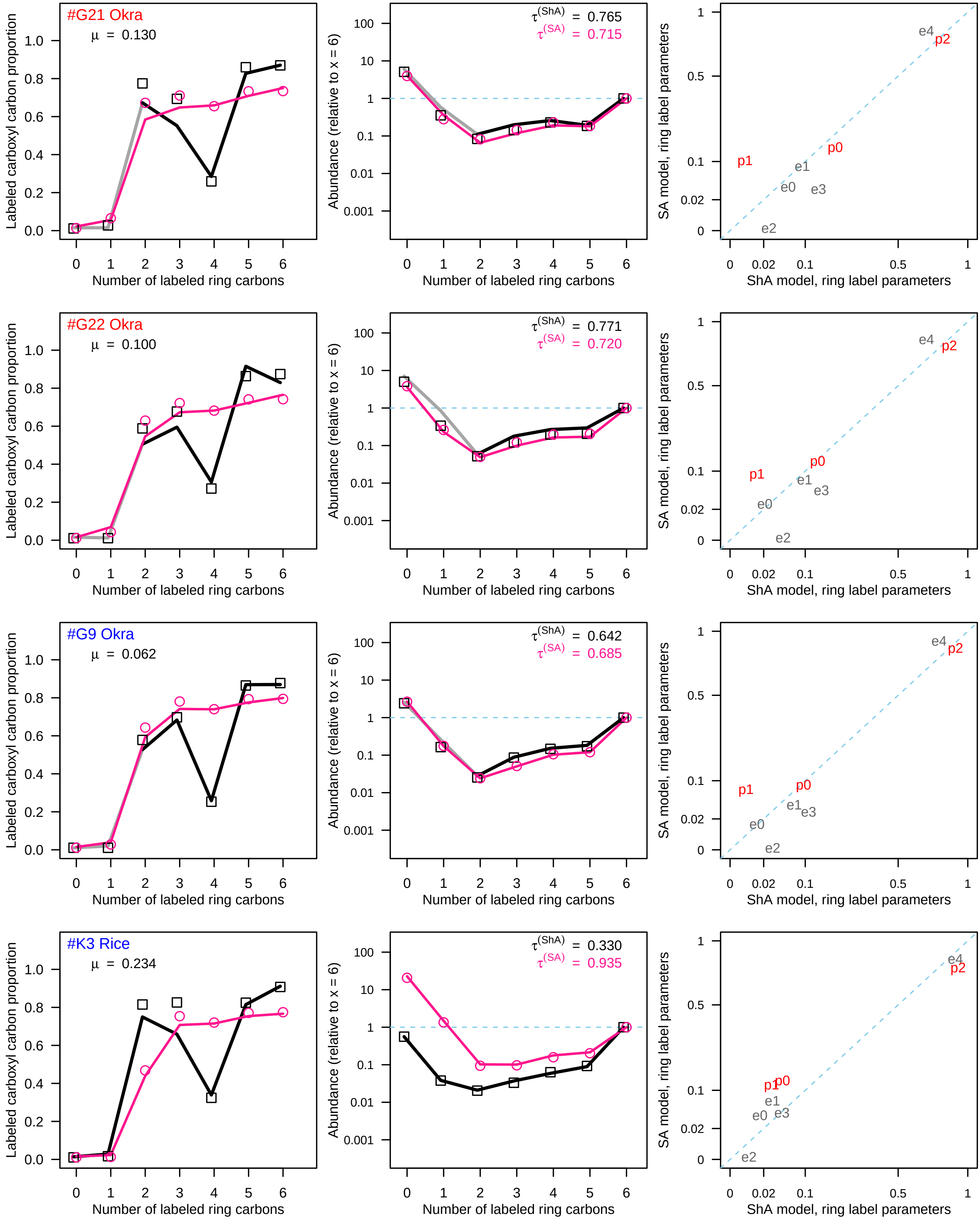

#G21 Okra
μ = 0.130
τ(ShA) = 0.765
τ(SA) = 0.715
#G22 Okra
μ = 0.100
τ(ShA) = 0.771
τ(SA) = 0.720
#G9 Okra
μ = 0.062
τ(ShA) = 0.642
τ(SA) = 0.685
#K3 Rice
μ = 0.234
τ(ShA) = 0.330
τ(SA) = 0.935
Labeled carboxyl carbon proportion
Number of labeled ring carbons
Abundance (relative to x = 6)
SA model, ring label parameters
ShA model, ring label parameters
p0
p1
p2
e0
e1
e2
e3
e4

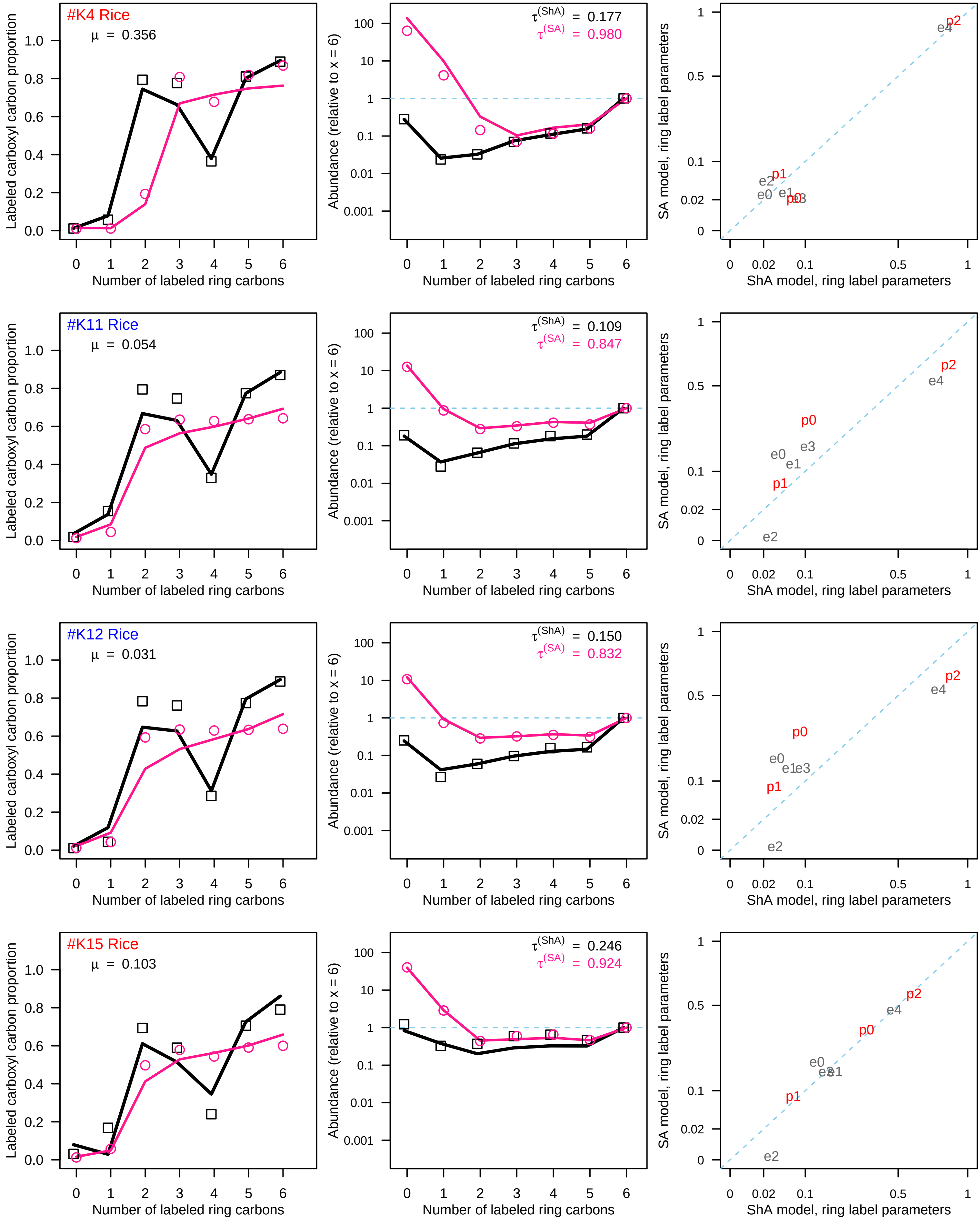
#K4 Rice
μ = 0.356
τ(ShA) = 0.177
τ(SA) = 0.980
#K11 Rice
μ = 0.054
τ(ShA) = 0.109
τ(SA) = 0.847
#K12 Rice
μ = 0.031
τ(ShA) = 0.150
τ(SA) = 0.832
#K15 Rice
μ = 0.103
τ(ShA) = 0.246
τ(SA) = 0.924
Labeled carboxyl carbon proportion
Abundance (relative to x = 6)
SA model, ring label parameters
Number of labeled ring carbons
ShA model, ring label parameters
p0
p1
p2
e0
e1
e2
e3
e4

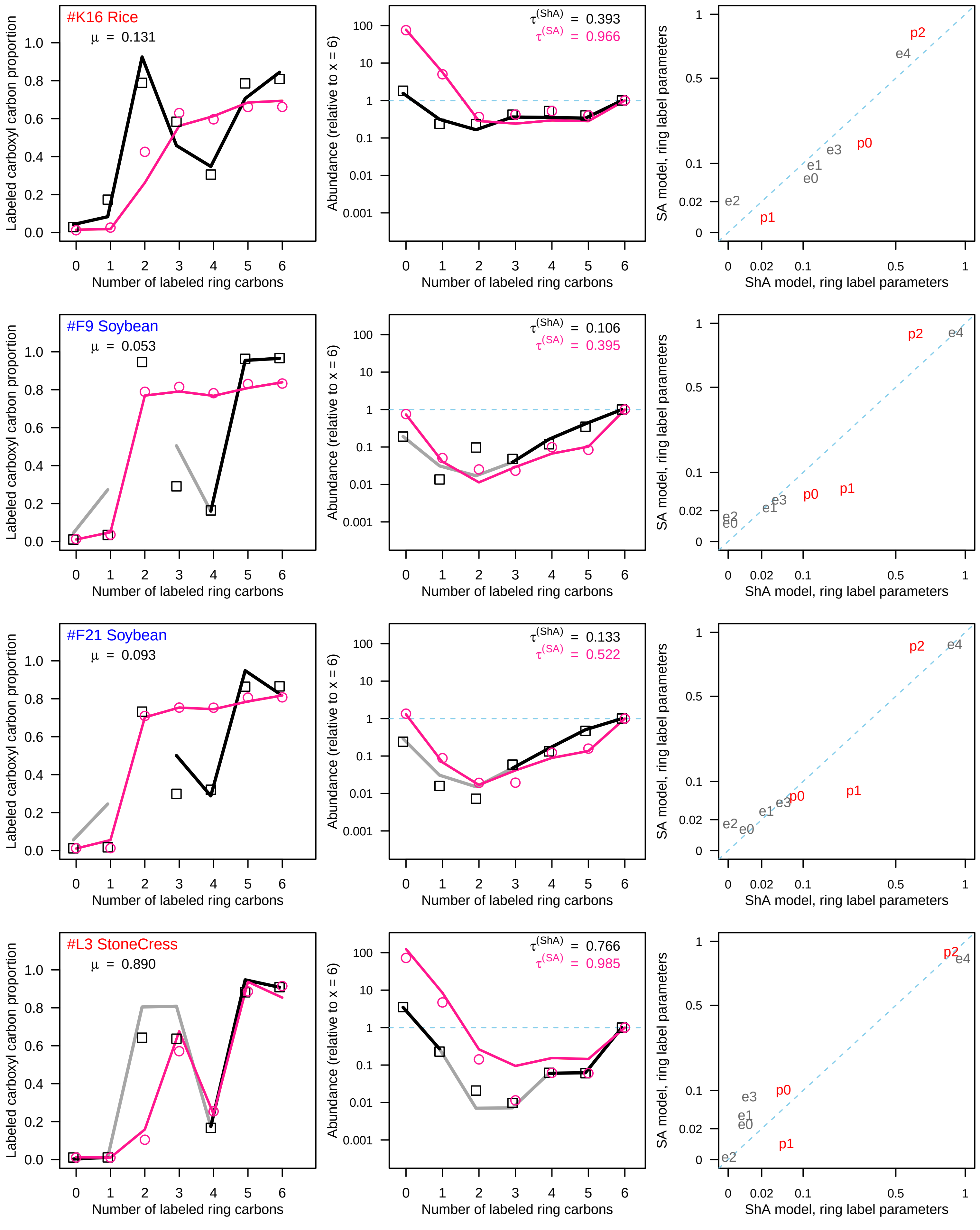
#K16 Rice
μ = 0.131
#F9 Soybean
μ = 0.053
#F21 Soybean
μ = 0.093
#L3 StoneCress
μ = 0.890
Labeled carboxyl carbon proportion
Number of labeled ring carbons
Abundance (relative to x = 6)
τ(ShA) = 0.393
τ(SA) = 0.966
τ(ShA) = 0.106
τ(SA) = 0.395
τ(ShA) = 0.133
τ(SA) = 0.522
τ(ShA) = 0.766
τ(SA) = 0.985
SA model, ring label parameters
ShA model, ring label parameters
p0
p1
p2
e0
e1
e2
e3
e4

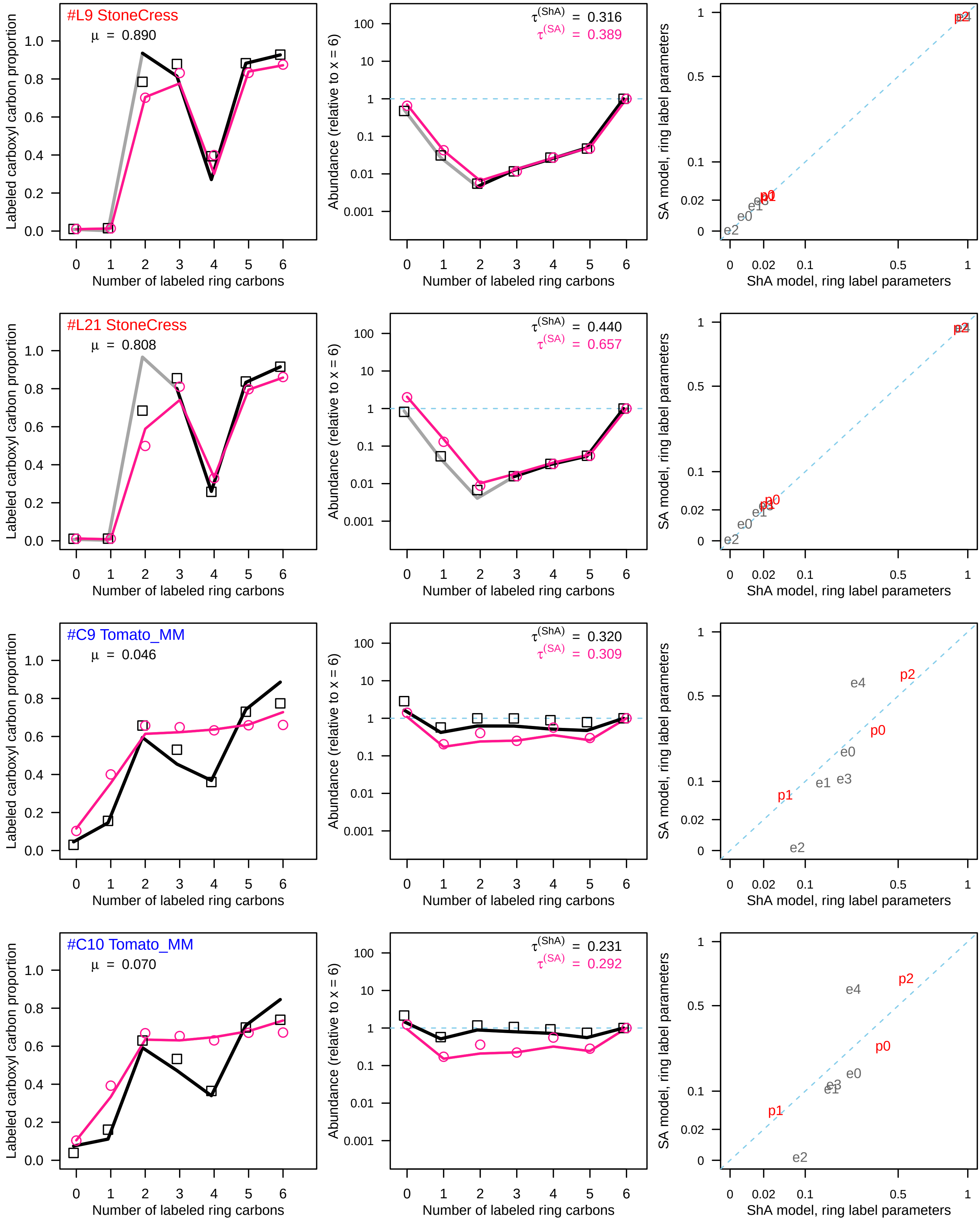

#L9 StoneCress
μ = 0.890
τ(ShA) = 0.316
τ(SA) = 0.389
#L21 StoneCress
μ = 0.808
τ(ShA) = 0.440
τ(SA) = 0.657
#C9 Tomato_MM
μ = 0.046
τ(ShA) = 0.320
τ(SA) = 0.309
#C10 Tomato_MM
μ = 0.070
τ(ShA) = 0.231
τ(SA) = 0.292
Labeled carboxyl carbon proportion
Number of labeled ring carbons
Abundance (relative to x = 6)
SA model, ring label parameters
ShA model, ring label parameters
p0
p1
p2
e0
e1
e2
e3
e4

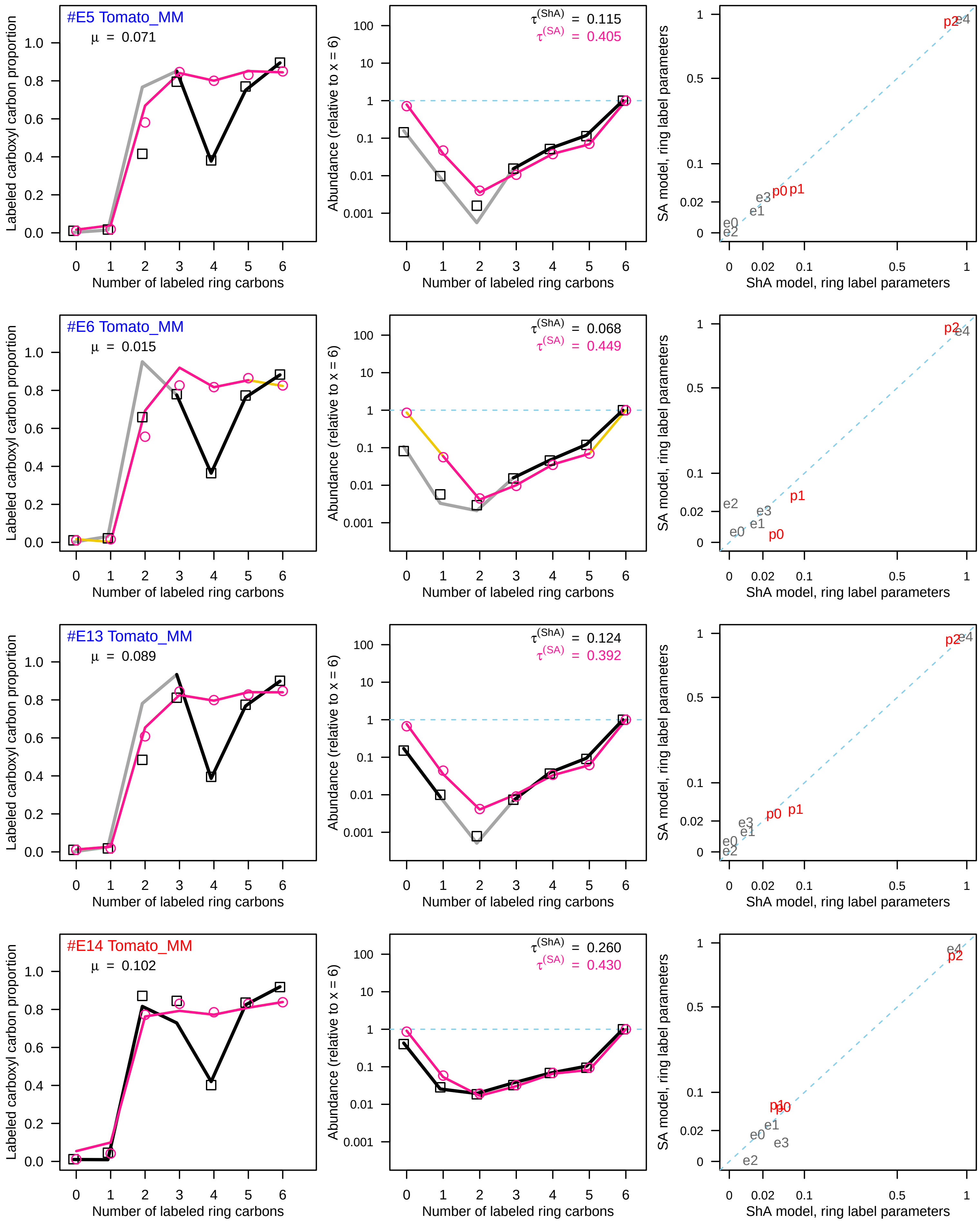
#E5 Tomato_MM
μ = 0.071
$\tau^{(ShA)}$ = 0.115
$\tau^{(SA)}$ = 0.405
#E6 Tomato_MM
μ = 0.015
$\tau^{(ShA)}$ = 0.068
$\tau^{(SA)}$ = 0.449
#E13 Tomato_MM
μ = 0.089
$\tau^{(ShA)}$ = 0.124
$\tau^{(SA)}$ = 0.392
#E14 Tomato_MM
μ = 0.102
$\tau^{(ShA)}$ = 0.260
$\tau^{(SA)}$ = 0.430
Labeled carboxyl carbon proportion
Abundance (relative to x = 6)
SA model, ring label parameters
Number of labeled ring carbons
ShA model, ring label parameters
p0
p1
p2
e0
e1
e2
e3
e4

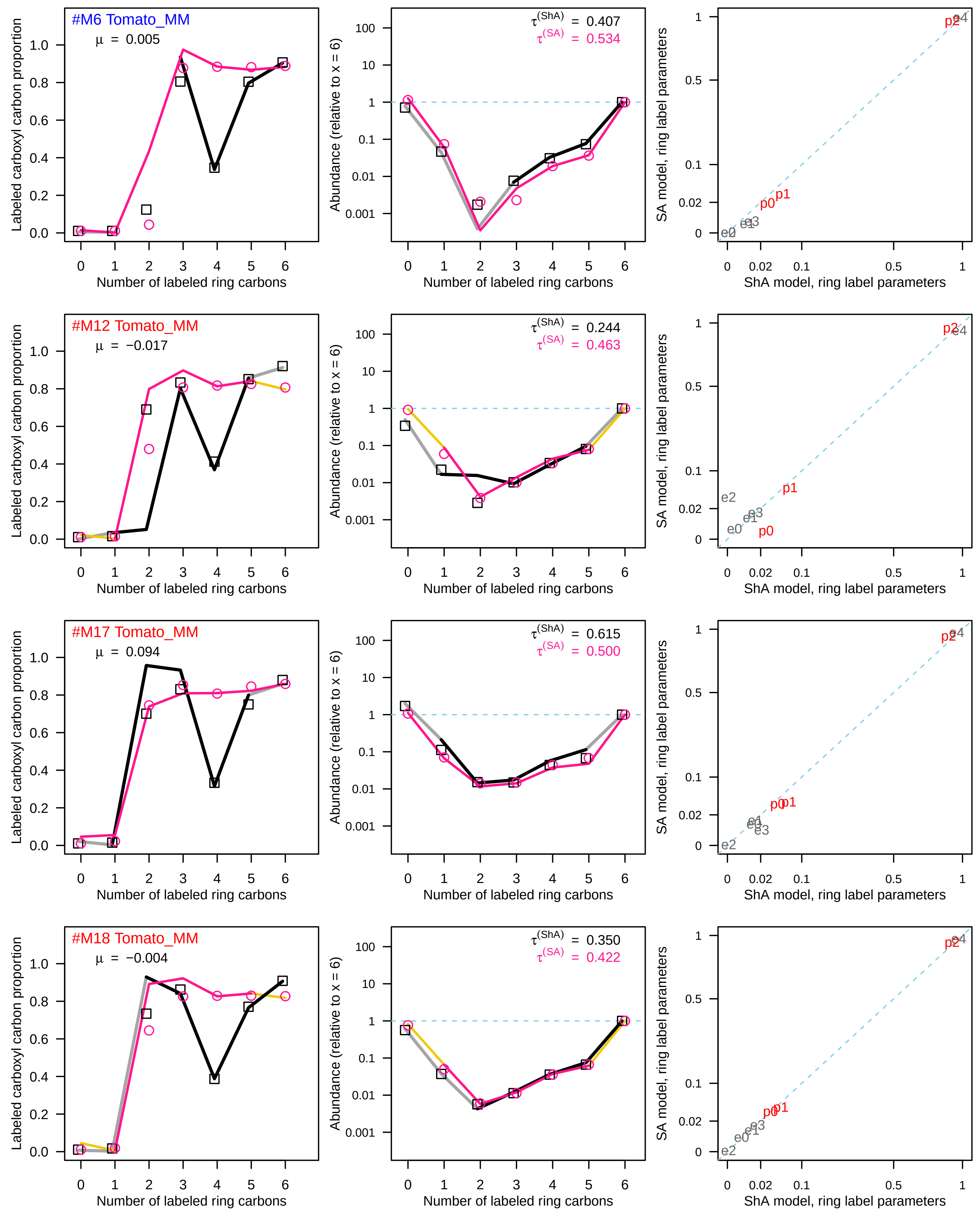
#M6 Tomato_MM
μ = 0.005
τ(ShA) = 0.407
τ(SA) = 0.534
#M12 Tomato_MM
μ = −0.017
τ(ShA) = 0.244
τ(SA) = 0.463
#M17 Tomato_MM
μ = 0.094
τ(ShA) = 0.615
τ(SA) = 0.500
#M18 Tomato_MM
μ = −0.004
τ(ShA) = 0.350
τ(SA) = 0.422
Labeled carboxyl carbon proportion
Abundance (relative to x = 6)
SA model, ring label parameters
Number of labeled ring carbons
ShA model, ring label parameters

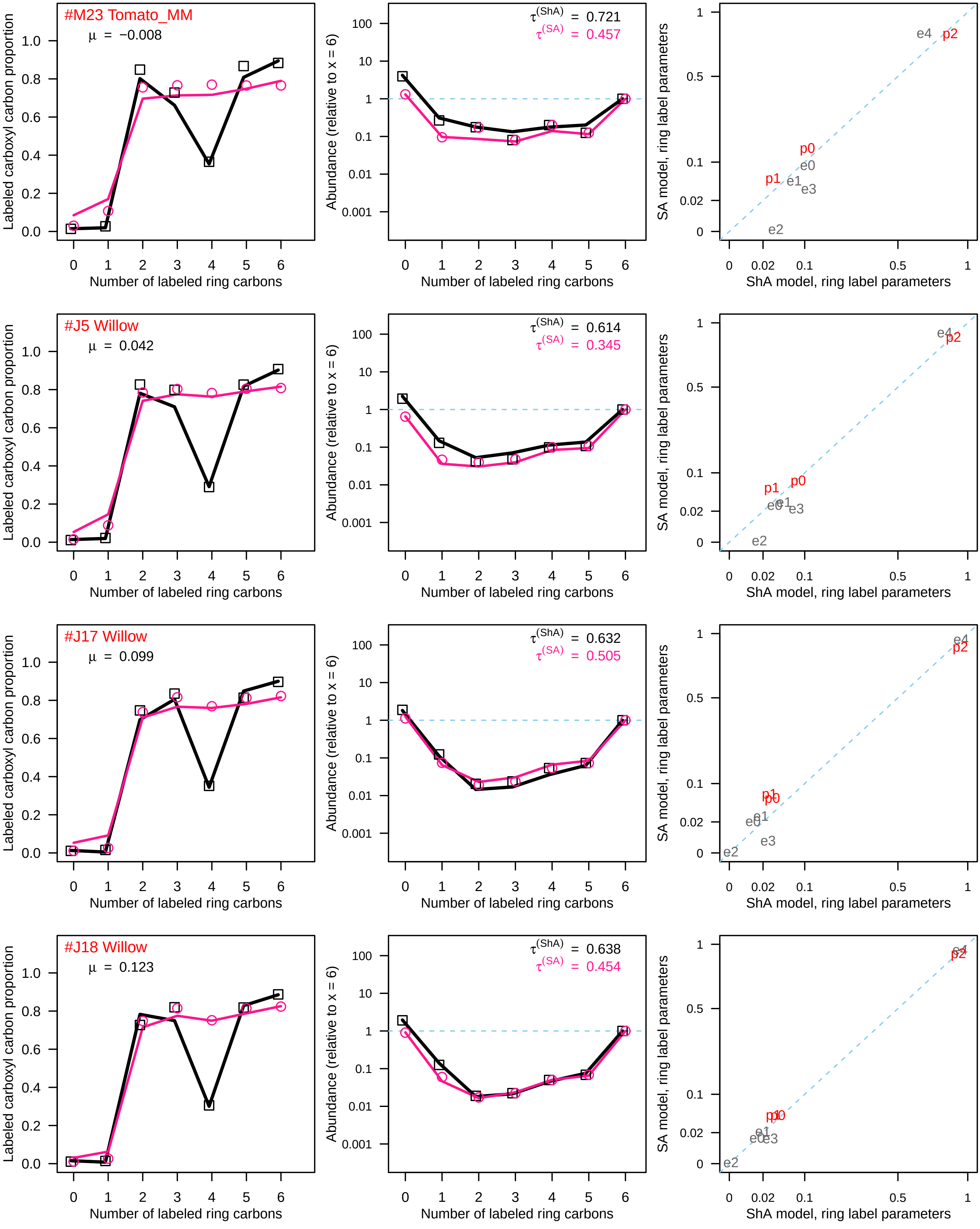
#M23 Tomato_MM
μ = −0.008
τ(ShA) = 0.721
τ(SA) = 0.457
#J5 Willow
μ = 0.042
τ(ShA) = 0.614
τ(SA) = 0.345
#J17 Willow
μ = 0.099
τ(ShA) = 0.632
τ(SA) = 0.505
#J18 Willow
μ = 0.123
τ(ShA) = 0.638
τ(SA) = 0.454
Labeled carboxyl carbon proportion
Abundance (relative to x = 6)
SA model, ring label parameters
Number of labeled ring carbons
ShA model, ring label parameters
p0
p1
p2
e0
e1
e2
e3
e4
