## Supplementary figures and images for "Biosynthesis of a major plant immunity hormone, salicylate, underwent multiple drastic changes during evolution of flowering plants"

### Fig S3

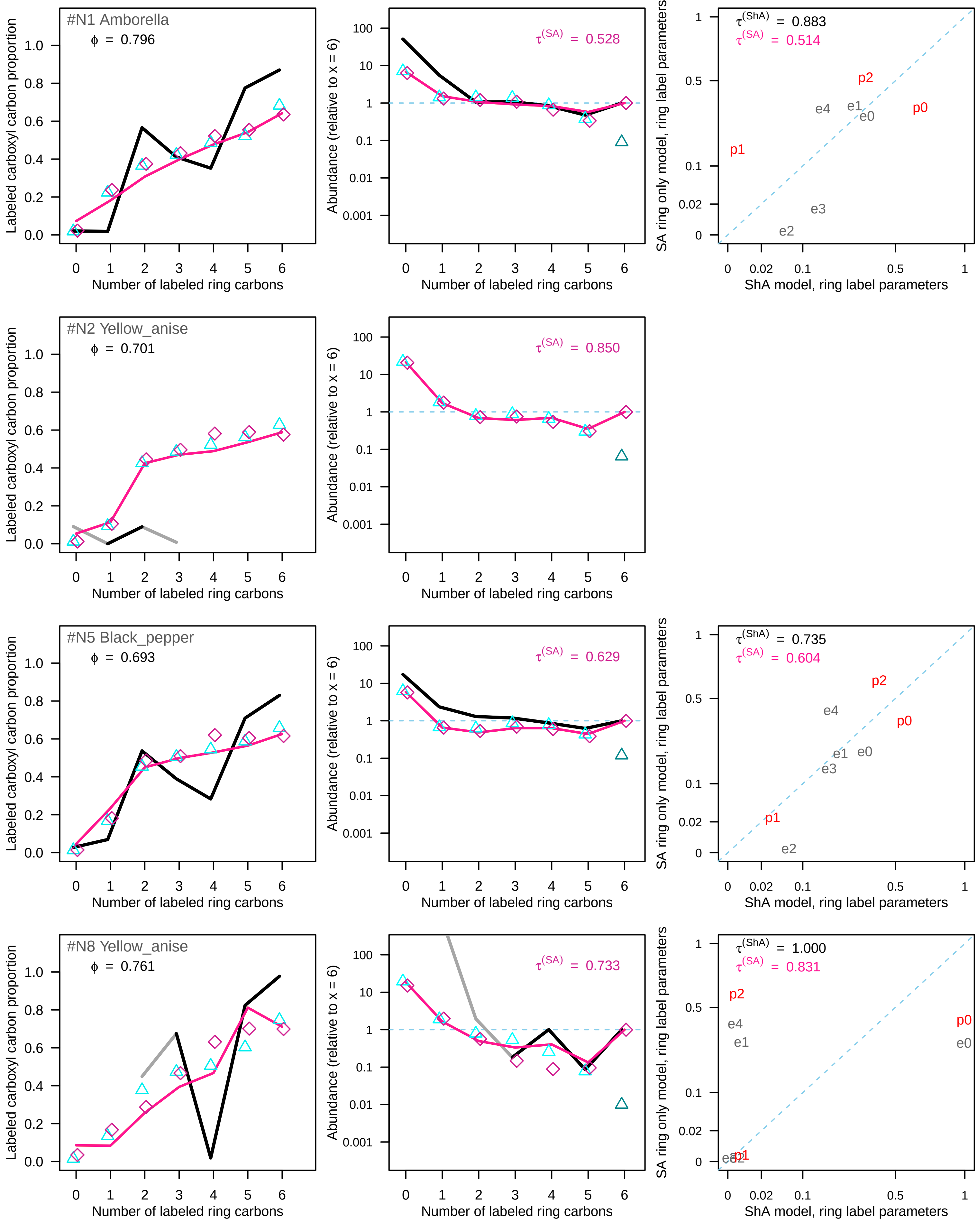

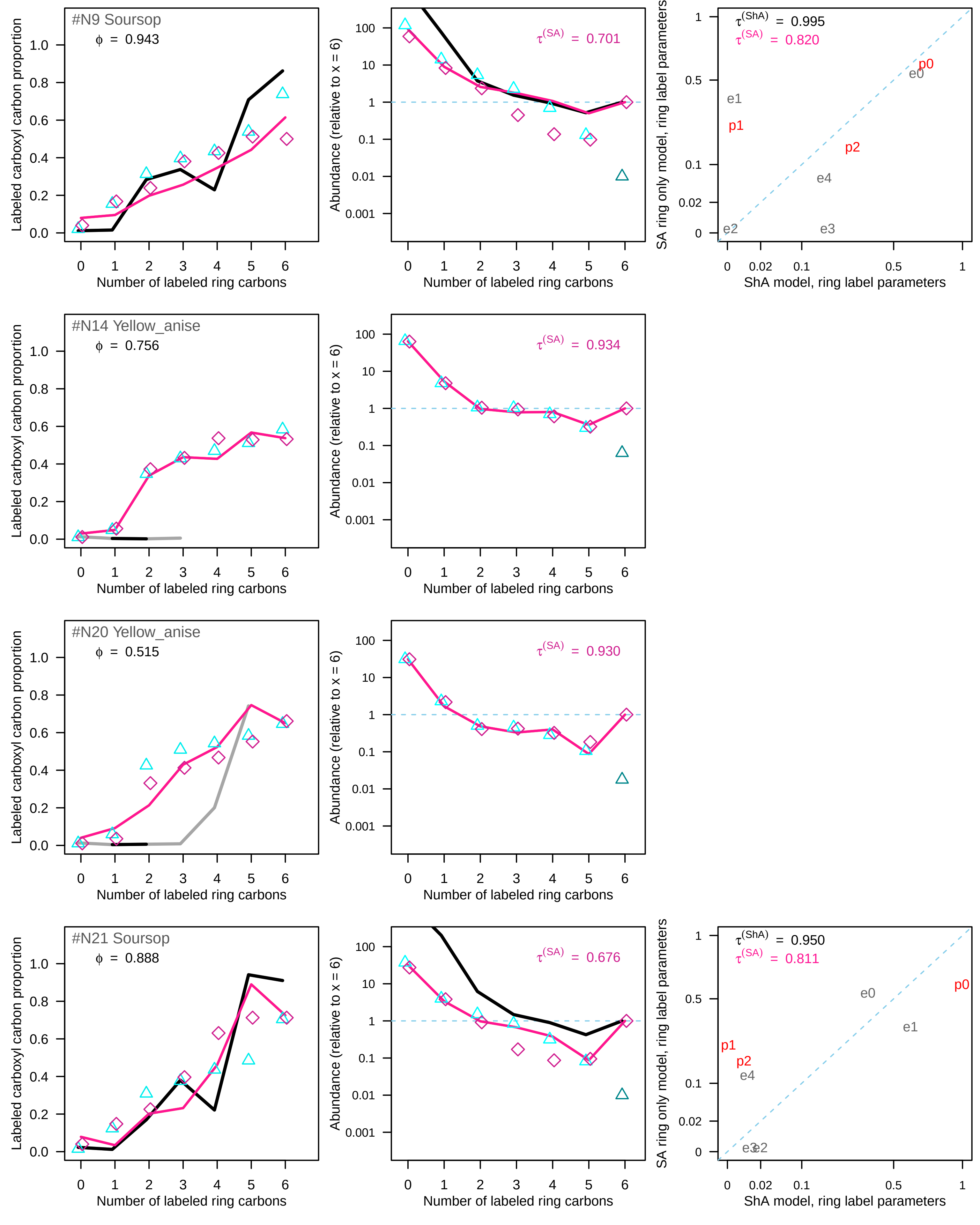

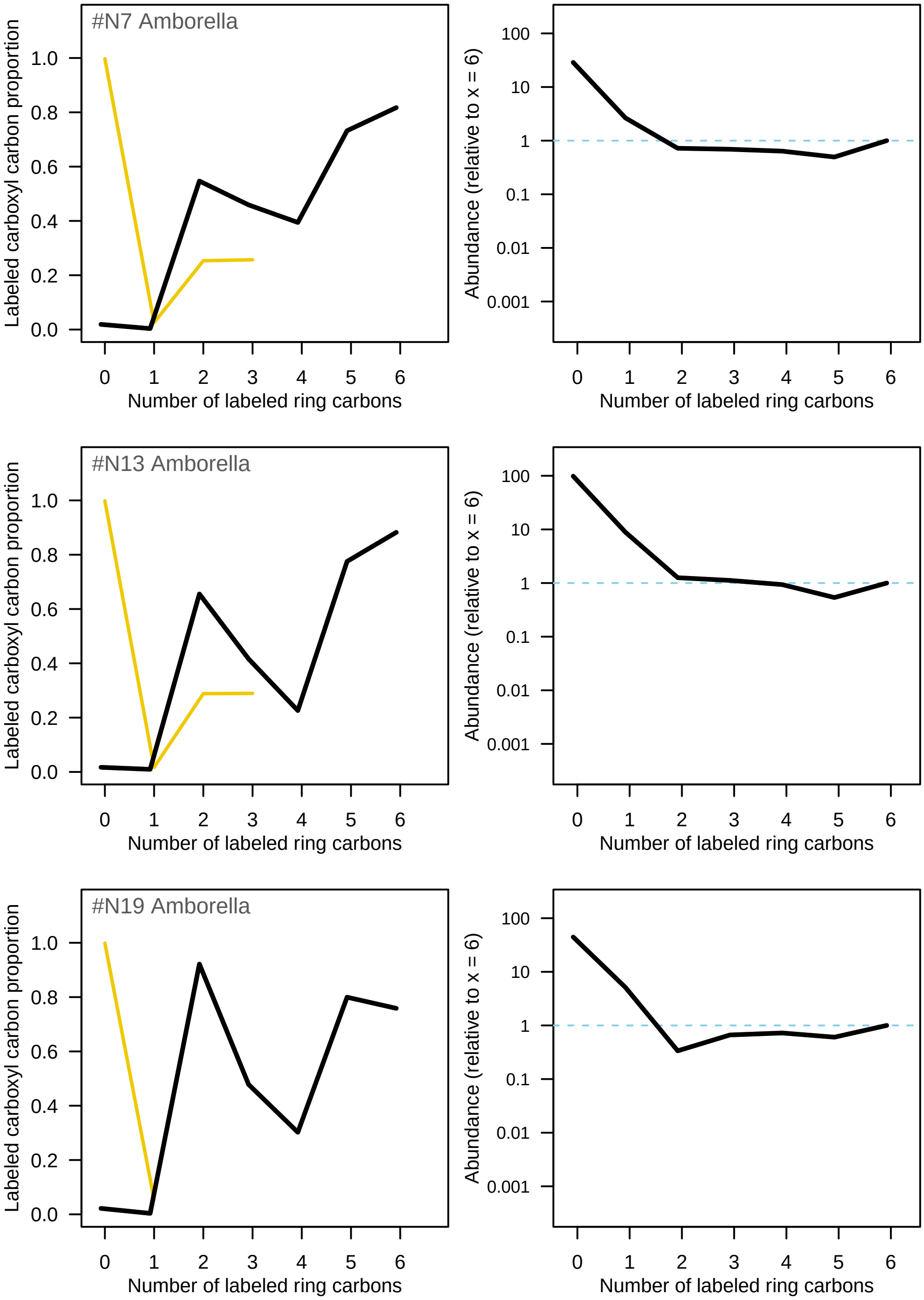
